# Evolutionarily Conserved Amyloid Aggregation in the PACAP Peptide Family Is Controlled by Heparin-Sensitive Lys/Arg Gatekeeper Residues

**DOI:** 10.64898/2026.05.04.722589

**Authors:** Dániel Horváth, Szebasztián Szaniszló, Zsolt Dürvanger, Zsolt Fazekas, Kim Hoang Yen Duong, András Perczel

## Abstract

Functional amyloid formation plays a central role in the storage of several peptide hormones, yet its structural basis remains poorly understood. Here, we investigate amyloid formation within class B1 GPCR-associated peptide hormones by analysing the aggregation propensity of the PACAP family. We show that five of six human PACAP-family peptides form amyloid fibrils not only within the pH range characteristic of secretory granules; however, aggregation is strictly dependent on the presence of glycosaminoglycans such as heparin. We identify conserved Arg/Lys-rich regions as key regulatory elements that act as conditional switches within the PACAP family. Experimental and molecular dynamics simulations show these residues function as aggregation gatekeepers but promote peptide self-assembly upon charge compensation by polyanionic cofactors. Although the glucagon and PACAP families diverged early in chordate evolution, we demonstrate, supported by seven amyloid-like crystal structures, that receptor-binding segments of PACAP peptides also function as aggregation-prone regions despite substantial sequence divergence from their counterparts in the glucagon family. Profiling aggregation in protochordate-derived peptides approximating the ancestral state of the superfamily, combined with sequence analysis of vertebrate PACAP–glucagon peptides, reveals that the intrinsic aggregation profile is remarkably conserved throughout evolution, even as receptor specificity diversifies.

## INTRODUCTION

Besides pathological amyloids associated with neurodegenerative^1^ and systemic diseases^2^, increasing attention has been paid to functional amyloids^3^ across all biological kingdoms. These fibrils may possess inherent physiological roles in various organisms^4–6^, yet their amyloid state can also be harnessed and applied in diverse chemical^7^ and biotechnological^8^ processes, representing a gain of function. Peptide-based hormones have been observed to form amyloids, either as a result of simple misfolding^9^ or as part of a proposed functional mechanism.^10^ Following post-translational modifications, the mature hormones awaiting secretion exist in a self-assembled state within acidic secretory granules.^11^ This transient amyloid state is thought to act as a compartmentalized reservoir system, allowing polypeptides to be selectively stored in a dense, phase-separated, and stabilized form.^12^ The local acidic pH^13,14^, together with the high peptide and glycosaminoglycan (GAG)^15^ concentrations, is suggested to promote amyloid self-assembly in secretory milieu. Upon release into the bloodstream, dilution, the shift from acidic to neutral pH^16^, and shear forces drive fibril disintegration into biologically active monomeric units.

The 3D structures of over 500 pathological amyloid fibrils were recently determined^17^ thanks to the *resolution revolution* in cryo-EM^18^, whereas only about a dozen structures of functional amyloids have been resolved. Despite their generally similar cross-β architectures the fibrillar structure of functional amyloids display a less energetically stable fold. This gives rise to their reversibility and distinct biochemical functions.^19^ To date, we only have atomic-resolution information on the amyloid state of only a few full-length hormone peptides, including β-endorphin^13^, insulin^9^, and glucagon (GCG)^20,21^. In our previous work^14^, we reported the amyloid-like crystal structures of the aggregation-prone regions (APRs) of the glucagon-related human peptides, glucagon-like peptide-1 and 2 (GLP-1 and 2), and glucose-dependent insulinotropic peptide (GIP). These conserved segments were shown to drive amyloid assembly by initiating self-association and stabilizing β-strand packing in the fibrillar state. At the same time, they are well-established determinants of GPCR recognition in the monomeric, bioactive form^22^, highlighting their dual functional role.

In humans, members of the PACAP–GCG (secretin) superfamily are divided into two main branches. The GCG branch comprises GCG, GLP-1, GLP-2, and GIP, whereas the PACAP branch includes secretin (SCT), vasoactive intestinal peptide (VIP), pituitary adenylate cyclase–activating polypeptide (PACAP), peptide histidine methionine/isoleucine (PHM/I), PACAP-related peptide (PRP), and growth hormone–releasing hormone (GHRH).^23^ (**Fig. 1/A, B)** All ten peptides of the superfamily found in humans are encoded by six structurally similar genes and are proposed to have arisen from a common ancestral PACAP-like gene.^24^ **(Fig. 1/A)** The two families diverged early in chordate evolution^25,26^ as a result of a genome duplication event. However, their common ancestral origin is reflected in their sequence and structural homology **(Fig. 1/C)**, their similar tissue distribution of the receptors^22,27^**(Fig. 1/D)** and their pleiotropic metabolic, nervous, endocrine and developmental functions.^28^ Owing to their remarkably diverse, yet still only partially characterized biological activities^29–34^, these peptides are often referred to as the brain–gut peptide superfamily too.^28^ The lengths of their sequences range from 27 to 48 amino acids; however, only the first 27 residues are directly involved in the canonical two-domain class B1 GPCR binding.^27^ Further elongation at the C-terminus and the presence of C-terminal amidation depend on the post-translational processing of the secreting cells and vary across species.^0,35^ Because of their high sequence homology **(Fig. 1/C)**, peptide hormones can bind not only to their dedicated cognate receptors but also to related receptors, often eliciting weaker or distinct biological responses^36,37^, which promiscuity can be therapeutically exploited.^38^

**Figure 1.**
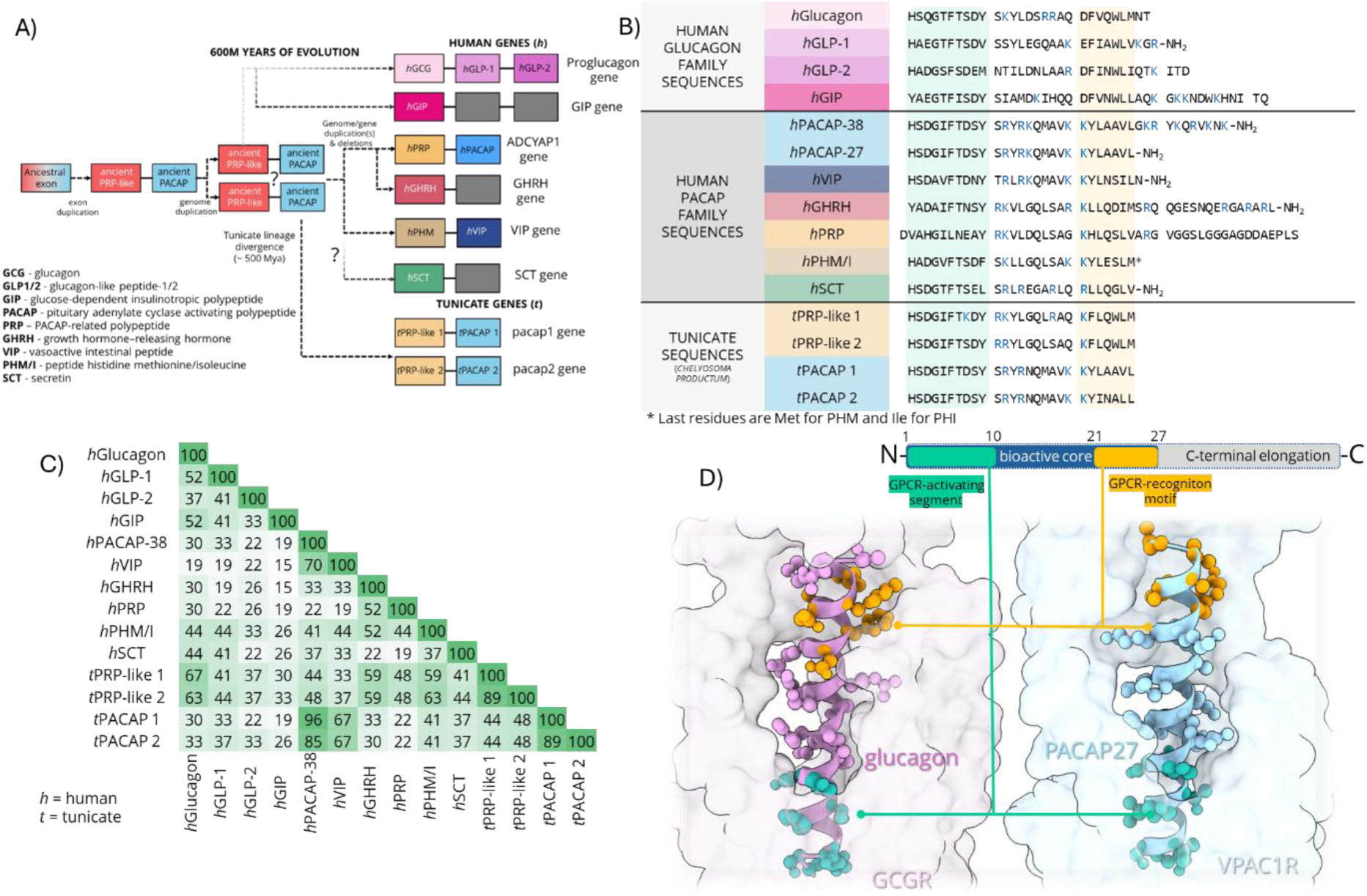
Sequences, evolutionary origin, and general GPCR binding mechanism of the PACAP–glucagon superfamily. **(A)** Proposed evolutionary lineage of the six human genes belonging to the superfamily. Schematic exon–intron structures are shown with peptide products annotated their corresponding coding exons. A primordial PACAP-like gene in early protochordates first underwent exon duplication, giving rise to an ancestral PRP/PACAP gene encoding precursors of the PRP/PHM/GHRH and PACAP/VIP peptide lineages. Subsequently, two rounds of whole-genome duplication expanded this ancestral locus into multiple paralogous genes, which later diverged to form the human PACAP/PRP, VIP/PHM, and GHRH genes (the latter accompanied by a secondary exon loss – grey coloured box). In parallel, an ancestral glucagon gene likely diverged before the genome duplication events, giving rise to GIP, and a later exon duplication within the proglucagon gene produced GLP-1 and GLP-2. The origin of SCT remains uncertain as the peptide is only found in tetrapods.^39^ **(B)** Amino-acid sequences of human and protochordate (*Chelyosoma productum*) PACAP–GCG family peptides. Positively charged Lys and Arg residues are coloured in blue, while GPCR-binding and receptor-activating segments are highlighted in green and yellow. **(C)** Pairwise sequence identity (%) across their 27-residue bioactive cores. In humans, both PACAP-27 and PACAP-38 are bioactive products generated from the same precursor by prohormone convertases. A generalized visualization of the peptides highlights the segments involved in GPCR binding and activation. **(D)** Extracellular views of the ligand–receptor complexes GCG–GCGR (PDB: 6LMK) and PACAP-27–VPAC1R (PDB: 8E3Y). In both complexes, the peptide ligand adopts an α-helical conformation along the whole sequence. The C-terminal segment of the hormone mediates recognition by the extracellular domain, whereas the N-terminal segment inserts into the 7-transmembrane (7TM) helical bundle to activate intracellular signalling. Among the ten human peptides, PRP is the only one without known biological activity, which is attributed to an extra N-terminal Asp residue preventing productive insertion into the receptor 7TM activation pocket.

This study aimed to expand our understanding of amyloid formation among class B1 GPCR-associated peptide hormones by examining the aggregation propensity of the human PACAP family. Specifically, we assessed whether PACAP peptides retain intrinsic aggregation potential despite early divergence from the GCG family, and whether their receptor-binding segments function as APRs despite substantial sequence variability within and between families (**Fig. 1/B, D**). These questions address whether amyloid-forming ability is more deeply conserved than receptor affinity, which shows greater evolutionary plasticity. ^26,28^

## RESULTS AND DISCUSSION

### 1. Screening the amyloid formation of the human PACAP family in presence of heparin *in vitro*

A central question was whether PACAP-family peptides possess an intrinsic capacity to form amyloid, and if so, under what physicochemical conditions this process is enabled. To address this, we systematically screened all six full-length human PACAP-family peptides using circular dichroism (CD) spectroscopy, thioflavin-T (ThT) fluorescence assays, and atomic force microscopy (AFM) **(Fig. 2, SFig. 1-6)** across a broad pH range (2 < pH < 10), both in the absence and presence of the polyanionic glycosaminoglycan heparin (enoxaparin), which mimics the crowded, GAG-rich environment of secretory granules.^13,15^ Initially in absence of heparin, the freshly dissolved peptides exhibit predominantly disordered secondary structure over the entire pH range, as confirmed by CD spectroscopy and subsequent spectral deconvolution. **(SFig. 7)** The sole exception was SCT, which displayed increasing α-helical content at higher pH. This indicates that PACAP-family peptides do not rely on pre-formed secondary structure during self-assembly.

**Figure 2.**
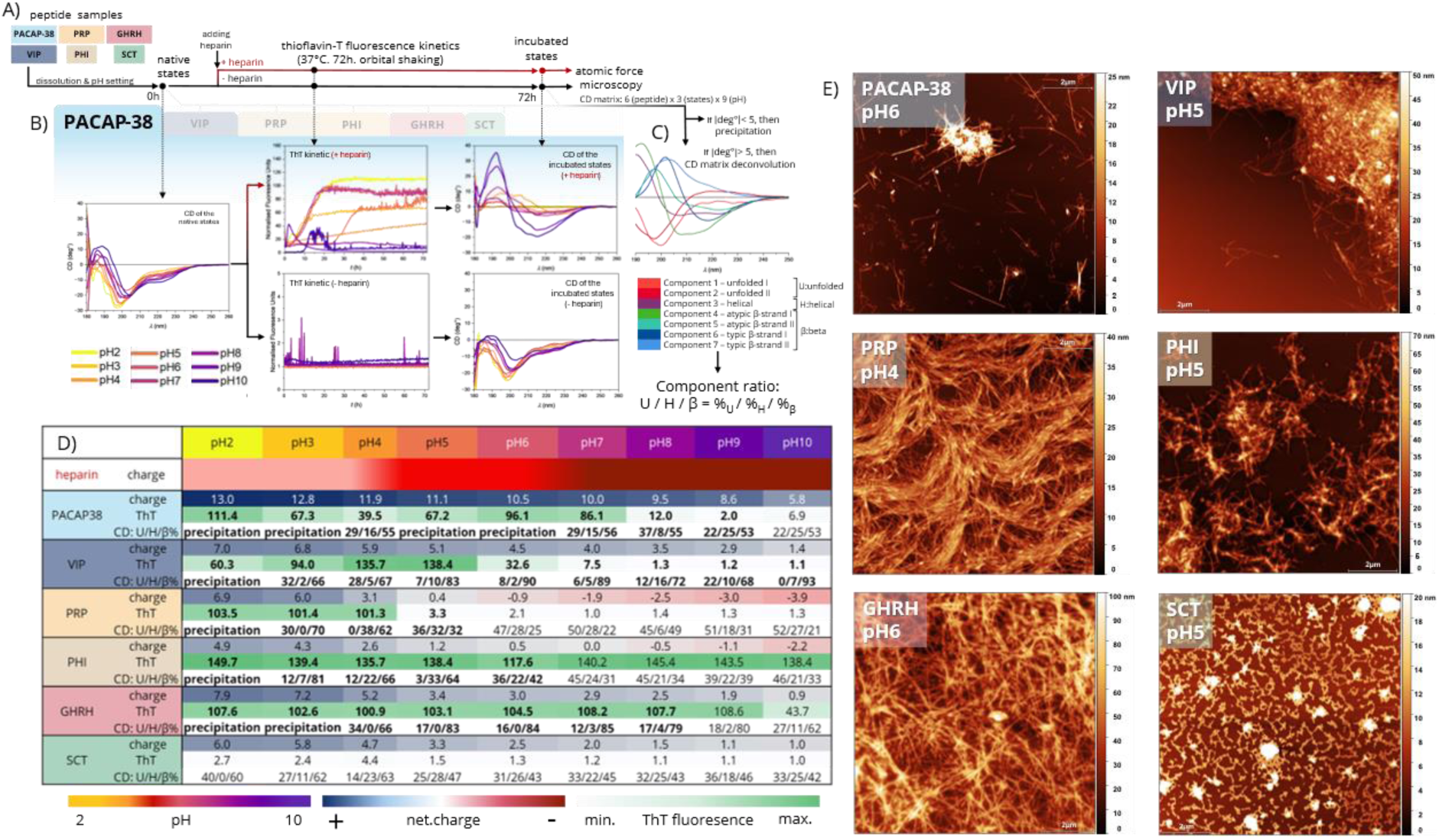
Characterizing the pH-dependent self-assembly of the PACAP family in the presence of heparin. **(A)** General overview of the experimental setup used to monitor amyloid formation. Peptides were applied at a final concentration of 0.35 mM. **(B)** CD spectra were recorded for the initial peptide states prior to incubation. The samples were then divided into two groups and incubated for 72 hours under Thioflavin-T fluorescence monitoring, either in the presence of enoxaparin sodium (2.125 mg mL⁻¹) or in the absence of heparin. The end states were subsequently analysed again by CD spectroscopy. For the rest of screening results see the supporting material. (PACAP-38 – **S**Fig. 1; VIP – **S**Fig. 2; PRP – **S**Fig. 3; PHI – **S**Fig. 4; GHRH – **S**Fig. 5; SCT – **S**Fig. 6) **(C)** The deconvolution of the collected CD curves yields 7 components: two slightly different unfolded components; one helical component and two-two typical and atypical β-sheeted components. For the in-depth review of the CD deconvolution see **S**Fig. 7. **(D)** Overview of the measured biophysical characteristics of the end states extended with the overall charge of the peptides across the examined pH range. ThT values represent the fold increase in fluorescence relative to the corresponding blank, whereas CD values indicate the relative proportions of unfolded, helical, and β-sheet components determined by spectral deconvolution. ThT values and component triplets are highlighted in bold in cases where amyloid fibrils were confirmed by AFM. Overall charges: heparin carries a negative charge under all investigated pH values. The sulfonate groups of polyanionic heparin remain fully deprotonated between pH 2 and 10. In contrast, its carboxyl groups are predominantly protonated at low pH, present in roughly equal protonated and deprotonated proportions near their pK_a_ and become fully deprotonated above pH 7. Approximate net charges of the peptides were calculated using generalized pKₐ values. **(E)** AFM micrographs recorded from the end states of the incubated samples after 20-fold dilution reveal nanoscale amyloid filaments with varying morphologies (**S**Fig. 1**-6****/G** shows additional micrographs), often accompanied by amorphous aggregates. PACAP (pH 6 – Fig. 2**/C**) and GHRH (pH 4 – **S**Fig. 5**/G**) display clear polymorphism within the same sample. In the case of SCT incubated at pH 5, the observed structures are not typical amyloid fibrils; however, they are too structured to be considered mere precipitation.

Addition of heparin resulted in rapid, visually observable precipitation, particularly in samples below pH 6. Samples at low pH (pH 2–4) became the most turbid, leading to a loss of CD signal after the 72-hour incubation period **(Fig. 2/D, SFig. 1–6).** Parallel CD and ThT measurements generally yielded consistent results regarding amyloid formation; however, the signals were not always in agreement (*e.g*., PACAP-38 at pH 9–10; **SFig. 1/B,C**), and CD deconvolution occasionally produced ambiguous or unexpected secondary-structure distributions. To resolve these ambiguities, all samples that tested positive for amyloid formation by either method were subsequently examined by AFM. AFM screenings **(Fig. 2/C, SFig. 1-6/G)** provided direct nanoscale structural evidence that five of the six peptides, with SCT as the sole exception, assembled into amyloid fibrils within 72 hours in the pH 4–6, range characteristic of secretory granules. In several cases (*e.g.*, PACAP-38 at pH 6, **Fig. 2/C,** and GHRH at pH 4, **SFig. 5/G**), polymorphic assemblies coexisted within the same sample, consistent with the heterogeneous β-structure components identified by CD spectral deconvolution, underscoring the complexity of characterizing amyloid samples by CD spectroscopy alone.

**Figure 3.**
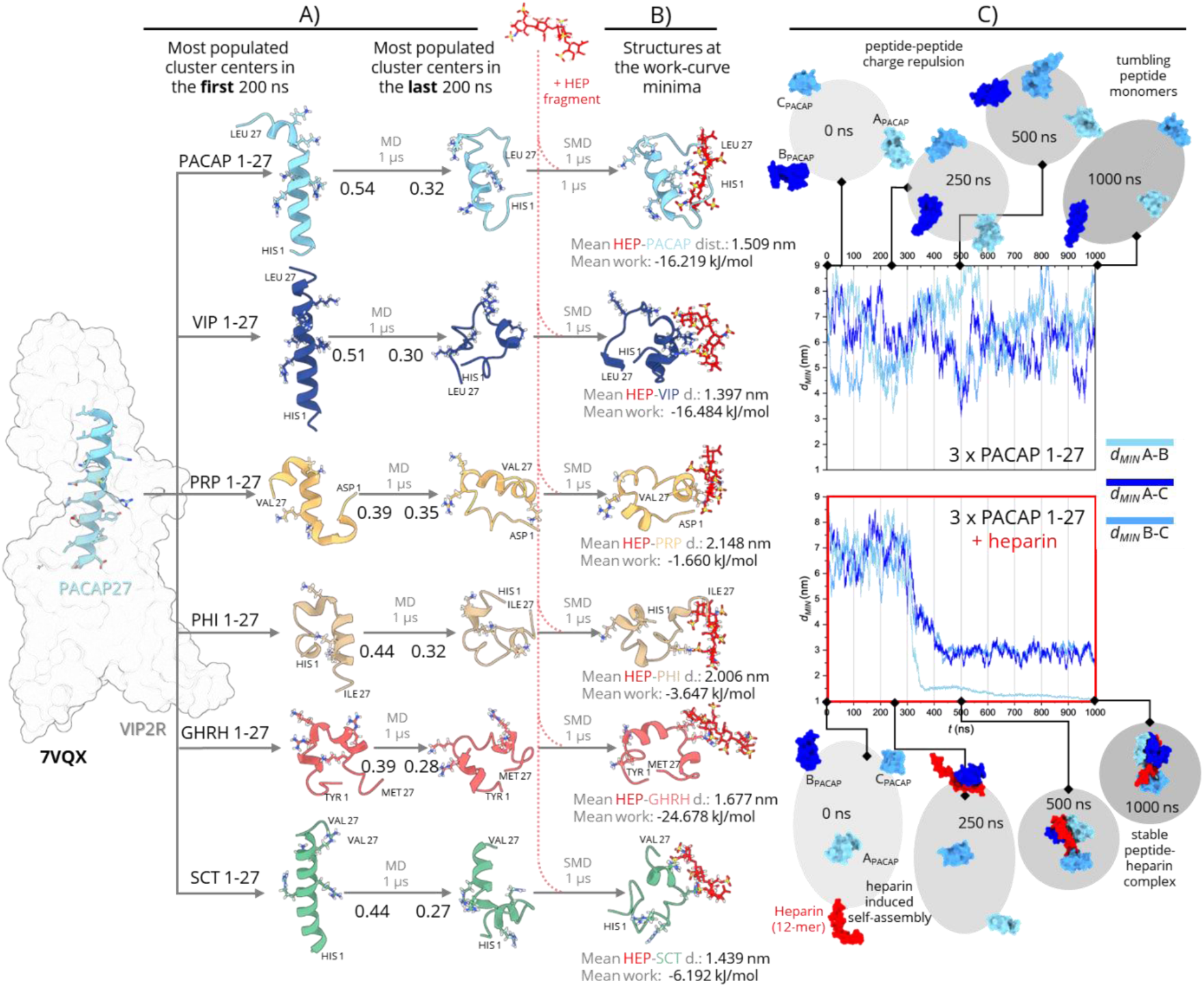
Molecular dynamic (MD) simulation of the PACAP related peptides in presence of heparin. **(A)** The PACAP-27–VIP2R cryo-EM complex structure (PDB ID: 7VQX) served as the starting model for the solution-phase MD simulations of the 27-residue bioactive peptides (C-terminally amidated). Helicity throughout the simulations was quantified using the GMM score, where 1 indicates maximal, and 0 indicates minimal helicity. The conformational ensembles of each peptide were grouped by clustering analysis; the most populated clusters representative of the first and last 200 ns trajectory segments are shown, with arrows indicating the average change in GMM score between these states. Over the course of the simulations, the initially rod-like helices adopted more globular conformations while retaining partial helical elements. In these conformations, Lys/Arg side chains remained preferentially solvent-exposed, generating extended positively charged surface patches, as illustrated by the electrostatic potential maps **(S**Fig. 9). **(B)** These Lys/Arg patches of partially disordered peptides are recognized by a four-sugar unit long heparin fragment (HEP) during steered molecular dynamics (SMD) simulations, leading to the formation of peptide–HEP complexes at pH=7. Representative structures at the minima of the work curves **(S.**Fig 10) are shown, along with the calculated work values and the distances between the geometric centres of the peptide and HEP molecules. **(C)** MD simulations of the self-assembly of three PACAP-27 peptides in the presence and absence of a 12-sugar unit long heparin over 1000 ns provide in silico insight into the additional peptide-recruiting role of GAGs beyond charge compensation. The simulation box size was chosen to approximate experimental peptide concentrations, without added ionic strength except for neutralising counterions. Peptide self-assembly was monitored by the minimum distances (*d_MIN_*) between peptide pairs (A–B, A–C, B–C), as well as peptide–heparin distances (see also **S**Fig. 11). In the absence of heparin (upper panel), the positively charged peptides do not form stable contacts due to electrostatic repulsion, reflected in persistent fluctuations of *d_MIN_* values (∼3–9 nm). In contrast, in the presence of heparin (lower panel), peptide–heparin interactions promote peptide recruitment and charge neutralization, leading to reduced intermolecular peptide-peptide distances (∼0.5-3 nm) and facilitating peptide clustering.

**Figure 4.**
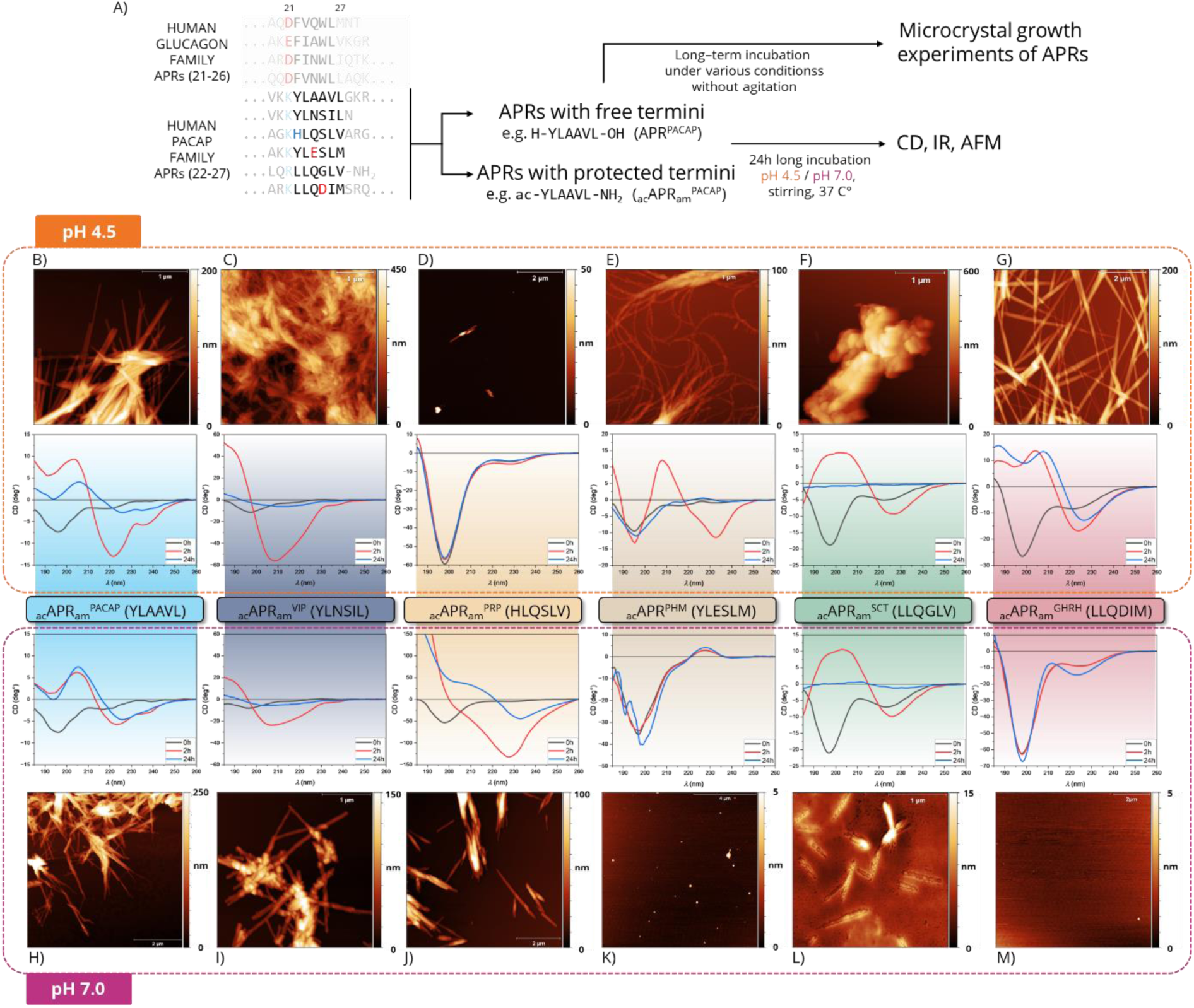
Receptor-binding hexapeptide segments of PACAP family with terminal protection are proved to be APR-s. **(A**) Positions of the investigated APRs within the full-length sequences of the glucagon family (residues 21–26) and the PACAP family (residues 22–27). Negatively and positively charged side chains are indicated in red and blue, highlighting aggregation gatekeeper residues in the vicinity of the APRs. The amyloid formation propensity of six PACAP-related APR hexapeptides was examined under identical experimental conditions, either with free termini or with N- and C-terminal protection. **(B-M)** Structural conversions observed during incubation of the APRs monitored by CD spectroscopy, together with the corresponding AFM micrographs recorded at the end of the incubation. All hexapeptides initially exhibit disordered structural features (black curves); however, variations in spectral intensity, particularly in the negative band at ∼200 nm, indicate differences in solubility despite identical peptide length and concentration. APRs that were subsequently proven to be prone to aggregation at the given pH display lower initial CD intensities, consistent with limited solubility and observable precipitation upon dissolution. In contrast, samples that remain unstructured at the end of the incubation generally exhibit more intense CD signals. **(D, J, M)** Characteristic yet diverse CD signatures indicate the formation of amyloid-like β-sheet–rich nanostructures within hours (red curves), suggesting rapid nucleation, as the intrinsically disordered APRs do not require an unfolding step prior to β-sheet formation. The CD signal intensity of these β-sheet–rich and yet soluble oligomers decreases over time (blue curves), indicating a liquid-to-solid phase separation that ultimately results in the formation of amyloid-like nanocrystals, as observed by AFM.

**Figure 5.**
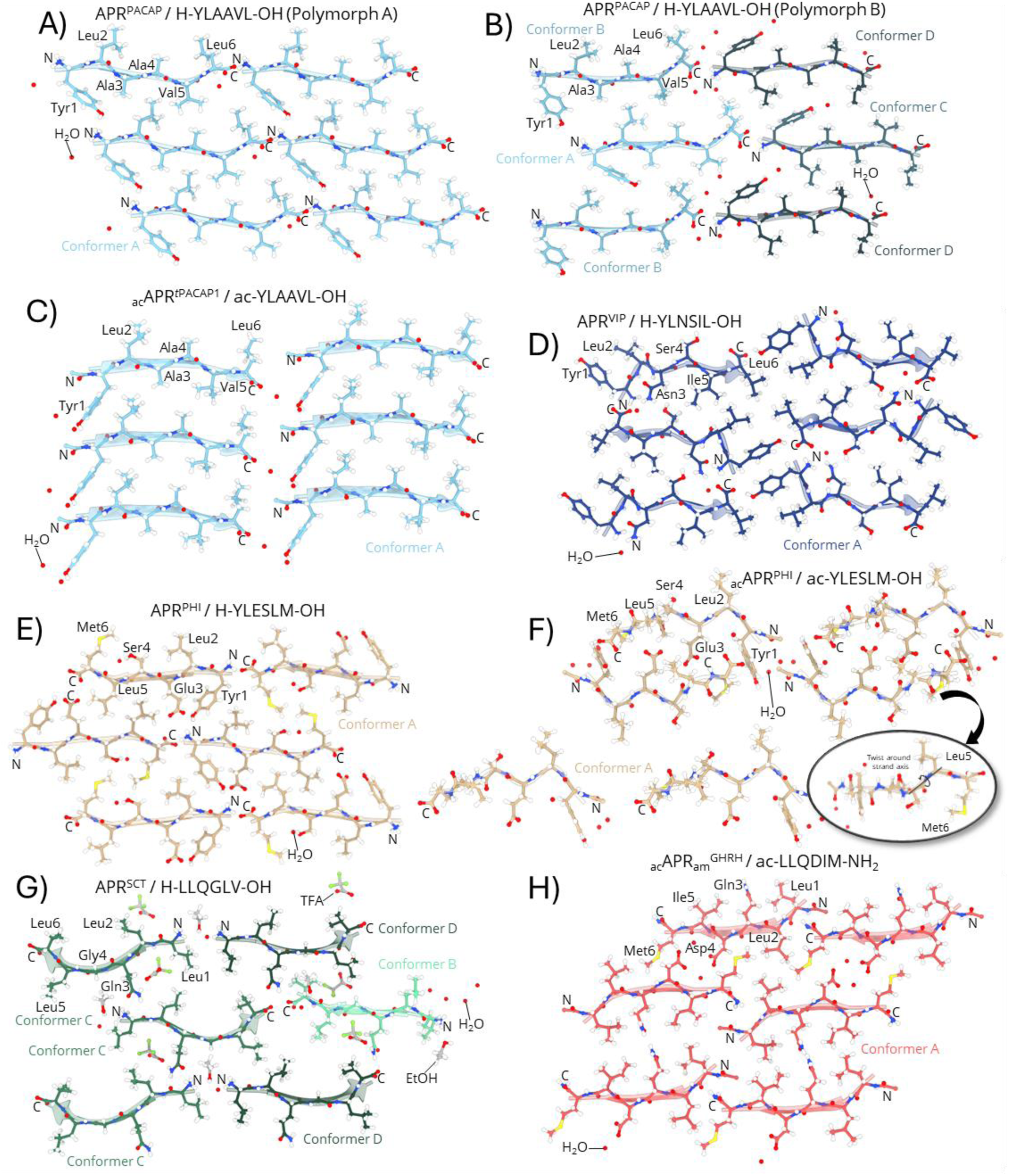
Steric zippers of the amyloid-like crystal structures reveal the major interactions in the APRs of the PACAP-family. (**A-H**) Six adjacent hexapeptides, arranged perpendicular to the fibril axis within the same plane of the crystal-lattice, were selected to visualize all possible side-chain interactions within the steric zipper interfaces of each APRs. β-strand packing along the fibril axis is shown in the Supporting Information **(S**Fig. 15).

Kinetic ThT measurements further revealed that aggregation is rapid in the presence of heparin, typically initiating within the first 24 hours and often exceeding the detection limit of the assay, thereby limiting full kinetic evaluation. Unlike pathological amyloids with prolonged lag phases, these peptides lack stabilizing higher-order structures; thus, the rapid amyloid formation observed in the presence of heparin is consistent with their functional role, as secretory granule maturation requires fast amyloid assembly.

Importantly, amyloid formation was not restricted to conditions mimicking secretory granules (pH ∼5.5) but extended across a broader pH range. Under highly acidic conditions (pH < 4), amyloid formation appeared to be a general feature and was likely accompanied by amorphous aggregation and precipitation. Notably, AFM clearly demonstrated the presence of fibrils formed by VIP, GHRH, and PACAP-38 at pH 7 (and even higher pH), where fibril formation would be unexpected based on current biological models.^10^

Analysis of the screening results obtained in the presence of heparin reveals a clear relationship between net peptide charge and the observed amyloid aggregation landscape. While the polyanionic heparin carries a consistently negative charge across all pH values, the peptides are predominantly positively charged. **(Fig. 2/B, SFig. 1-6/F)** At acidic pH, all PACAP-related peptides carry a maximal positive net charge, and rapid, uncontrolled precipitation competes with organized fibril formation. At physiological and higher pH values (pH > 7), heparin carries its maximal negative charge, while the peptides become less positively charged and PRP with PHI even acquire an overall negative charge; consequently, neither forms fibrils under these conditions. These findings suggest that when the peptides bear a mildly positive charge, favorable electrostatic interactions enable partial charge compensation between the peptides and heparin, thereby promoting controlled self-assembly and the subsequent emergence of fibrillar structures. This example of electrostatic matching between appropriately charged peptides and polyanionic partners suggests a general mechanism that promotes self-assembly toward amyloid formation.

### 2. Conserved Arg/Lys gatekeeper residues in PACAP family act as heparin-binding sites that trigger aggregation upon charge neutralization

Despite their pronounced aggregation propensity in the presence of heparin, none of the PACAP-family peptides formed amyloids under identical conditions in its absence across a broad pH range (2-10) over 72 hours (**SFig. 1-6/D-E)** This highlights the essential role of polyanionic GAGs in initiating amyloid nucleation within the PACAP peptide family. Moreover Arg/Lys residues are abundant across all six peptides exhibiting a conserved distribution pattern within the central 11–26 segment, with additional enrichment in the C-terminal extensions of PACAP and GHRH. **(Fig. 1/B)** We suggest that these residues serve a dual role. On one hand, they act as classical aggregation gatekeepers^39^, as their persistent positive charge under physiological conditions prevents self-association by introducing electrostatic repulsion and imposing an unfavourable entropic penalty through side-chain immobilization. On the other hand, their sequential arrangement closely resembles canonical heparin-binding motifs found in proteins, such as XBBXBX for α-helices or XBBBXXBX for β-sheets, where B denotes lysine or arginine and X a hydrophobic residue^40,41^. In these motifs, the precise spacing of cationic residues is critical for efficient heparin binding.^42–44^

**Figure 6.**
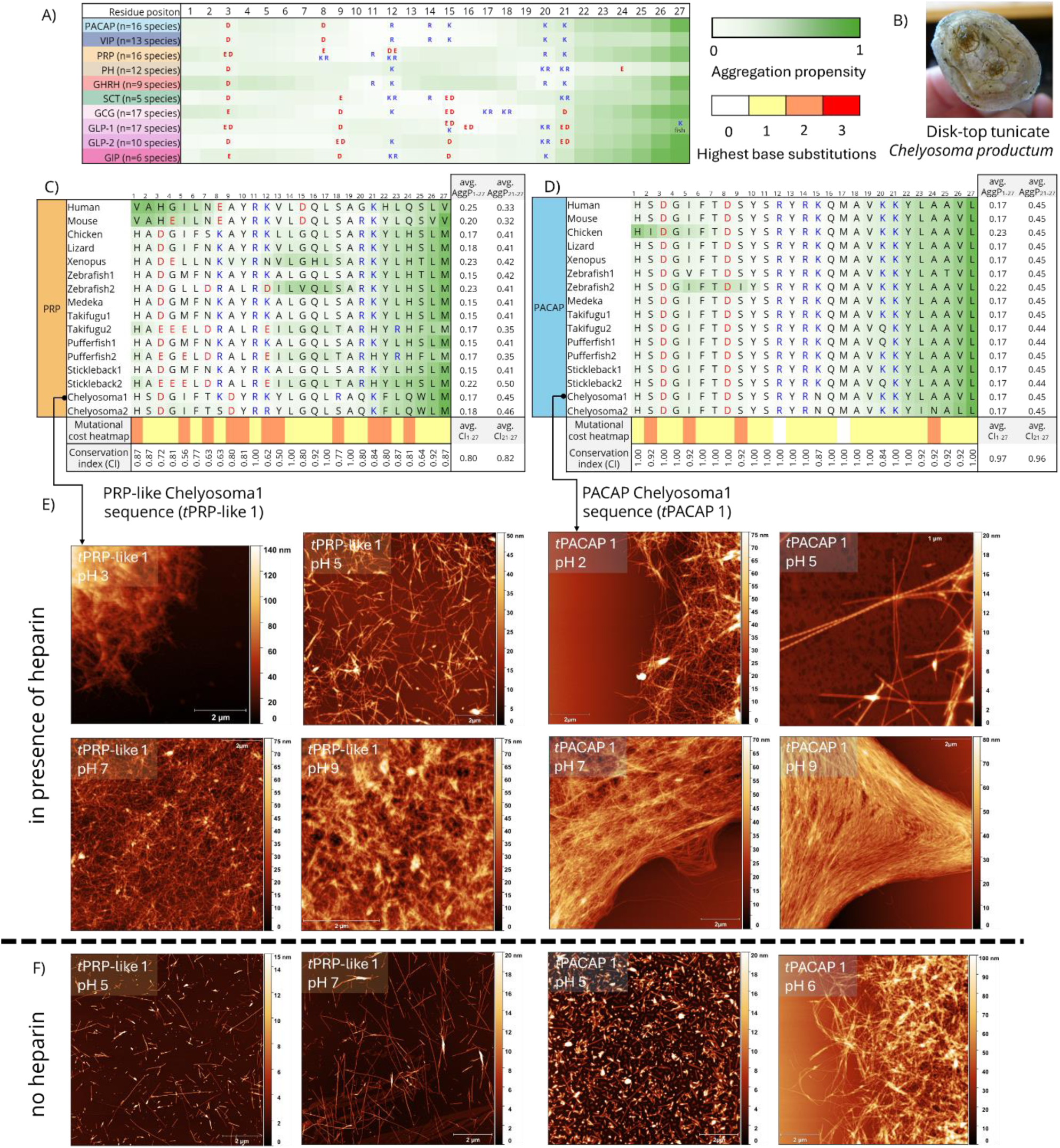
Analysis of the evolutionary conservation of position-specific aggregation propensity and gatekeeper patterns, together with the aggregation properties of the disk-top tunicate *Chelyosoma productum* PACAP-related peptides. (A) Position-specific mean aggregation propensity for each peptide family across the analysed species (n), calculated using AggreProt. Aggregation scores were computed for each sequence, averaged across species, and mapped along the peptide alignment as a heatmap. Distribution of negatively charged Asp/Glu gatekeepers (red) and positively charged Arg/Lys gatekeepers or heparin-binding residues (blue) within the 27-residue bioactive core across the analysed species. (B) *Chelyosoma productum* is a sessile marine tunicate belonging to the subphylum Urochordata, a group of protochordates considered the closest living relatives of vertebrates. Photograph credit: Carita Bergman. Detailed species-level aggregation profiles of the **(C)** PRP-(like) and **(D)** PACAP peptide families. (See **S** Fig. 20 for the remaining eight PACAP-GCG peptides) Species are arranged from bottom to top according to evolutionary order. For each sequence, the residue normalised mean aggregation propensity of the full peptide (avg. _AggP1-27_) and the APR region (avg. AggP_21-27_) is indicated at the line’s end. (Legend continuous on the next page) (Fig.6 **continue)** Position-specific conservation (CI) of the most frequent residues is shown together with a mutational cost heatmap. Mutational cost heatmap indicating the maximal nucleotide substitution distance among the codons observed at each position; the highest observed distance is highlighted even when most codon pairs differ by fewer substitutions. Detailed position-specific mutational pathways are listed in **S.Table 4/1-10**. Although the PACAP amino acid sequence at the APR position is highly conserved, its coding nucleotides have diversified extensively, exploiting nearly all synonymous codon possibilities without altering the protein sequence, indicating exceptionally strong evolutionary constraint, underscoring its critical functional importance, and suggesting an intrinsic, sequence-encoded aggregation propensity. In contrast, PRP and its ancestral sequences appear to have evolved more rapidly. **(S**Fig. 19) Although the APR segments differ considerably in amino acid composition across protochordate, non-mammalian, and mammalian species, these differences are largely separated by single-nucleotide substitutions at the codon level. In PRP, mammalian sequences are particularly divergent from those of non-mammalian species, as also indicated by the hierarchical clustering analysis. This divergence may reflect the loss of biological activity of PRP in mammals—likely due to a one-residue N-terminal elongation—after which its original function was taken over by GHRH, resulting in reduced evolutionary constraints on sequence conservation. Amyloid aggregation mapping of the PRP-like and PACAP peptides encoded by the two exons of the tunicate PACAP1 gene, the closest homolog of the human ADCYAP1 gene. Aggregation of the tunicate-derived full-length *t*PRP-like1 and *t*PACAP1 peptides was examined in the presence **(E)** and absence **(F)** of heparin. Unlike their human counterparts, these peptides readily formed amyloid fibrils across a broad pH range even in the absence of heparin. Detailed screening results are shown in **S**Fig. 22 and 23.

**Figure 7.**
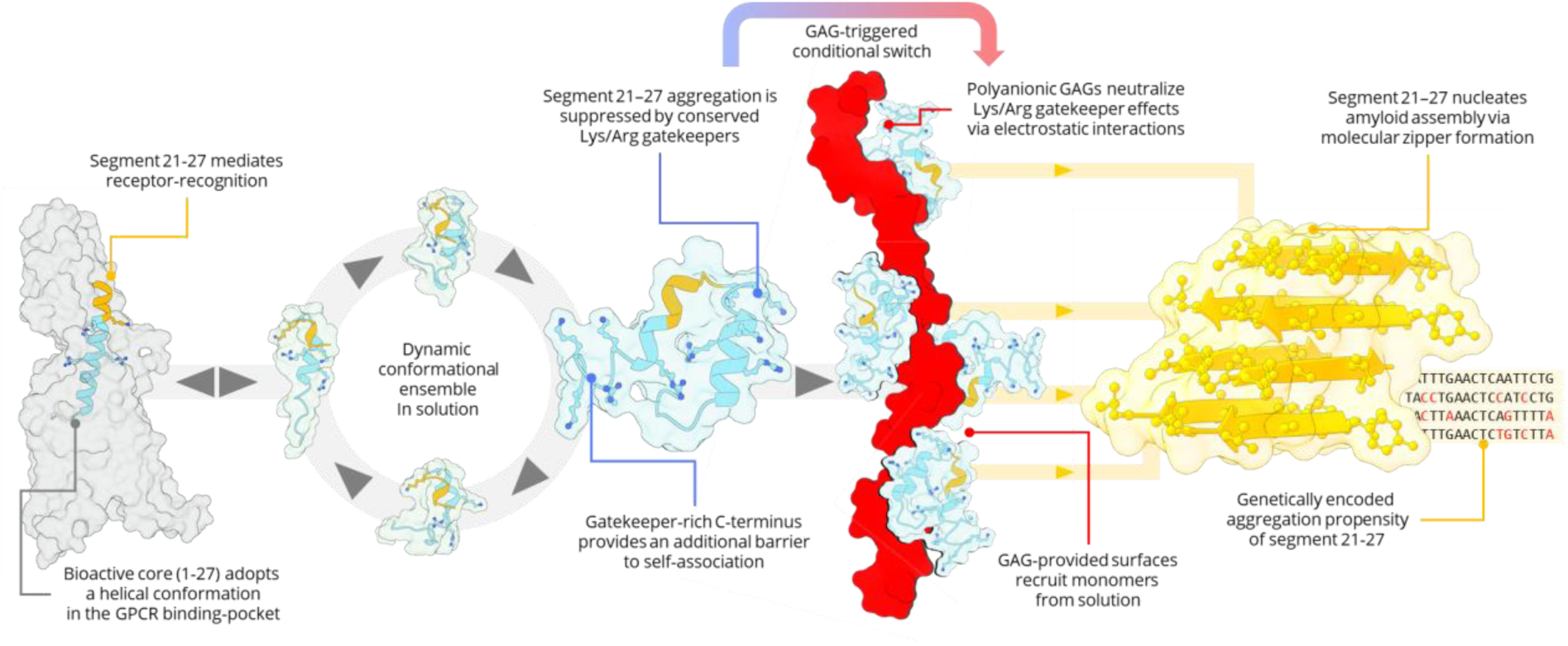
Overview of the conditional regulation of amyloid formation in PACAP-family peptides. Consistently, heparin-assisted pH-scan experiments revealed that aggregation occurs across a broader pH range than that of secretory granules, with the key determinant being peptide net charge. As long as peptides remain positively charged, interactions with negatively charged cofactors promote self-assembly. Thus, aggregation is governed by a combination of intrinsic sequence features and environmental conditions.

To further assess the validity of these putative heparin-binding segments **(SFig. 8**) we performed molecular dynamics (MD) simulations of the 1–27 bioactive regions of all six peptides. Using the fully helical VIP2R bound PACAP-27 peptide as a starting structure, we generated homologous models of the remaining five peptides by *in silico* mutagenesis **(Fig. 3/A)** and performed 1 μs long MD simulations at pH 7 to examine the intrinsic conformational behaviour of these models in and solution in the absence of any binding partners. Helicity was quantified in each frame using the Gaussian Mahalanobis Mean score (GMMs)^45^ **(SFig. 9**), showing that all six peptides progressively lost helical content (GMMs ≈ 0.3) and adopted a molten globule–like state by the end of the simulation. **(Fig. 3/A)** These findings support the experimental CD and deconvolution results, indicating that in solution the peptides do not maintain the canonical α-helical conformation but rather exist as an ensemble of fluctuating disordered states that carry intrinsic helical propensity.

To investigate peptide–heparin interactions, steered molecular dynamics (SMD) simulations were performed at pH 7, in which peptides and a four-sugar unit long heparin fragment were driven toward each other while recording the work along the reaction coordinate. Although these profiles do not represent fully converged free energy curves, they provide insight into relative binding preferences. Despite apparent Arg/Lys-mediated interactions in all cases, the work profiles differ markedly between peptides. (**Fig 3/B**, **SFig. 10**) PRP and PHI, which did not form amyloid at pH 7 in the presence of heparin *in vitro*, exhibit shallow minima (∼–2 to –4 kJ mol⁻¹) at larger (geometric center) separation distances (∼2 nm), indicating a weak and less stable binding. In contrast, PACAP, VIP, and GHRH, which do form amyloids under these conditions, display significantly deeper minima (∼–16 to –24 kJ mol⁻¹) at shorter distances (∼1.5 nm), consistent with tighter peptide–heparin complex formation. These results are in agreement with experimental observations and support a model in which amyloid formation is initiated by peptide–heparin complex formation. This interaction is primarily governed by electrostatics: at pH 7(**SFig. 9**), peptides with net charge closer to neutrality (*e.g.*, PRP, PHI) exhibit weaker binding, whereas more positively charged peptides form stronger complexes with anionic GAGs. Secretin displays intermediate behaviour (∼–6 kJ mol⁻¹), yet does not form fibrils, indicating that charge neutralization alone is not sufficient for amyloid formation. At pH 7, peptides closer to neutrality bind weakly, whereas more positively charged peptides form stronger complexes with anionic GAGs. Furthermore, MD simulations of three separate PACAP-27 peptide chains in the presence and, parallelly, absence of 12-sugar unit long heparin reveal that GAG chains are able to act as scaffolds that recruit peptides from solution, thereby increasing the local concentration of charge-neutralized species and promoting their subsequent self-assembly. (**Fig. 3/C, SFig. 11**).

Together, these findings indicate that Lys/Arg clusters function as conditional switches: they suppress aggregation under physiological conditions but enable rapid self-assembly upon charge compensation through heparin binding.

### 3. Mapping the aggregation-prone region (APR) of the human PACAP family

The receptor-recognition segment of PACAP-family peptides (residues 21–27; **Fig. 1B**) was examined for aggregation-prone properties, as previously demonstrated for the GCG family.^14^ Prediction algorithms (TANGO^46^/AggreProt^47^) identified these regions as weakly aggregation-prone, with substantially lower propensity than those of the GCG family (**SFig. 12**). Another key difference is observed at position 21: while the glucagon family contains a negatively charged Asp/Glu gatekeeper, the PACAP family carries a heparin-sensitive Arg/Lys residue at the same position (**Fig. 4A**).

The aggregation propensity of the six putative APR segments (residues 22–27) **(Table 1.)** was monitored at 5 mg mL⁻¹ under pH 4.5 and pH 7.0. CD **(SFig. 13**) and infrared (IR) spectroscopy **(SFig. 14**) revealed that, all hexapeptides were initially unstructured. After a 24h-long incubation at 37 °C, only APR^VIP^ and APR^PHM^ showed β-sheet–like spectral signatures, indicating limited but intrinsic self-association. **(SFig. 13/C,D,G)**. Strikingly, masking the APR’s terminal charges profoundly altered this weak aggregation potential. N-acylated and C-amidated variants (denoted with the subscript am/ac; **Table 1**), which more closely mimic the chemical environment of the APR segment within the full-length peptide chain, rapidly formed amyloid-like assemblies within a few hours under the same experimental conditions as their non-protected counterparts. (**Fig. 4/B–M**). APRs lacking protonatable side chains (_ac_APR_am_^PACAP^, _ac_APR ^VIP^, _ac_APR ^SCT^) formed amyloid at both pH values, whereas Asp/Glu-containing sequences (_ac_APR^PHM^, _ac_APR ^GHRH^) did so only at pH 4.5, and the His-containing (_ac_APR ^PRP^) exclusively at pH 7.0.These observations demonstrate that neutralization of both terminal and side-chain charges is a key prerequisite for efficient APR self-association. Time-dependent CD measurements revealed initially soluble amyloid-like species with diverse β-sheet–like signatures after two hours of incubation. The subsequent decay of these signals suggests precipitation of the soluble species. Consistently, AFM showed the formation of large (hundreds of nanometers), straight, needle-like amyloid nanocrystals, in contrast to the full-length sequences, which formed fibrils with diameters of ∼10–20 nm. **(Fig. 2/C)** The only exception, _ac_APR^PHM^, produced curved fibrils that assembled into crystal-like superstructures **(Fig. 4/E)**, suggesting that short amyloidogenic segments may prefer rigid nanocrystalline packing over flexible fibrillar architectures.^14,48^

**Table 1.**
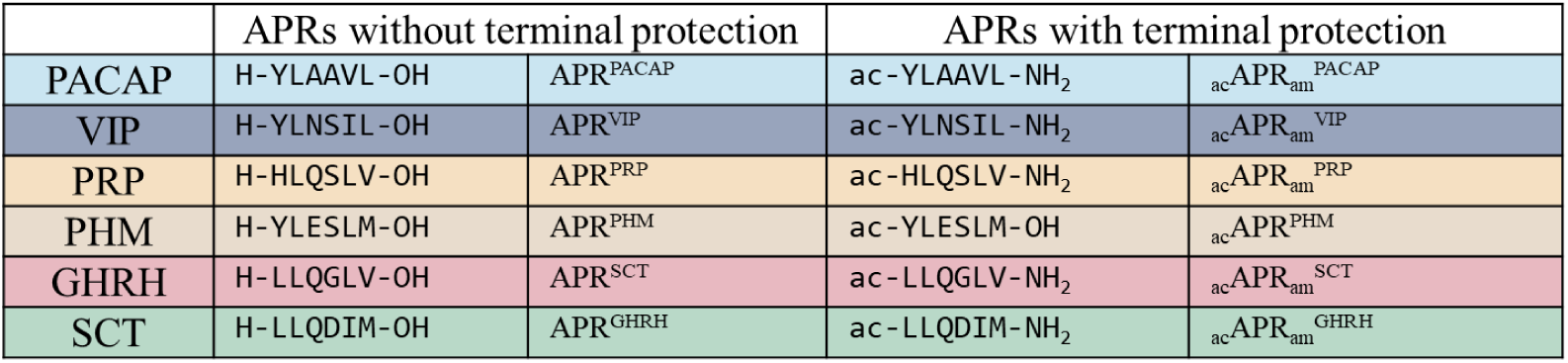
APR hexapeptide sequences of the PACAP-related peptides applied for the aggregation screening experiments. _ac_APR^PHM^ lacks C-terminal amidation, as this APR is located at the native C-terminus of the full-length peptide, where the terminus is naturally free.

These results demonstrate that the receptor-binding segments of the PACAP family also function as APRs contrary to ambiguous prediction outcomes. Comparison of terminally protected and unprotected forms shows that hydration of the zwitterionic termini counteracts the intrinsic aggregation propensity of these segments. The driving force for APR self-association, which is overall weaker than in the GCG family, becomes dominant only upon neutralization of terminal charges. Furthermore, the pH dependence indicates that neutralization of side-chain charges is essential for amyloid-like steric zipper formation, achieved through pH adjustment, charge-masking additives (*e.g.*, heparin), or intermolecular ionic interactions. To characterize steric zipper interactions at the molecular level, PACAP-related APRs were crystallized as amyloid-like microcrystals, previously observed by AFM but here grown to sizes suitable for X-ray diffraction.

### 4. Amyloid-like crystal structures reveal steric-zipper motifs of the PACAP family

Structural features of amyloid-like crystals formed by APR peptides, such as side-chain interactions within steric zipper interfaces, can provide valuable insights into key aspects of amyloid formation in their corresponding full-length proteins.^19,49,50^ We obtained eight diffraction-quality crystals from the APR peptides of five hormones; PACAP, VIP, PHM, SCT, and GHRH; some with and some without terminal protecting groups. **(Table 2**, **Fig. 5, SFig.15**) In the case of PRP, small needle-like crystals formed from _ac_APR ^PRP^; however, these were not suitable for diffraction data collection.

**Table 2.** Overview of the amyloid-like crystals formed by the PACAP family related APRs.

| Peptide | APR | Polymorph | PDB ID. | Crystallisation conditions | Backbone conformer(s) | Co-crystallising reagent(s) | Amyloid class<sup>a</sup> | Shifted assembly mode<sup>b</sup> |
| --- | --- | --- | --- | --- | --- | --- | --- | --- |
| PACAP | H-YLA AVL-OH | A | 29VP | 1 mg ml<sup>-1</sup>, 0.1 M HEPES buffer, pH 7, 37 °C | A<br>B | H<sub>2</sub>O | class 6 | Shifted along the β-spine axis and lateral shift between sheets |
| | | B | 29VQ | 1 mg ml<sup>-1</sup>, 0.2M acetate buffer, 30 v/v% acetonitrile pH 4.07, 37 °C | A<br>B<br>C<br>D | H<sub>2</sub>O | class 4.1 | Every second strand shifted along the β-spine axis and lateral shift between sheets |
| | ac-YLA AVL-OH | A | 29VO | 1 mg ml<sup>-1</sup>, 40% acetonitrile, pH 4 (dissolved at pH 7, filtered and diluted with 0.2M acetate, pH 4 + 40 v/v% acetonitrile), 37 °C | A | H<sub>2</sub>O | class 6 | Shifted along the β-spine axis |
| VIP | H-YLNSIL-OH | A | 29VR | 0.4 mg ml<sup>-1</sup>, 30-50 v/v% methanol, 0.2M acetate buffer, pH 3.77, 37 °C | A | H<sub>2</sub>O | class 4 | No shift |
| PHM | H-YLES LM-OH | A | 29VS | 1 mg ml<sup>-1</sup>, 0.2M acetate buffer, pH 4.67, 37 °C | A | H<sub>2</sub>O | class 1 | Lateral shift between sheets |
| | ac-YLES LM-OH | A | 29VT | 1 mg ml<sup>-1</sup>, 30 v/v% acetonitrile, 37 °C | A | H<sub>2</sub>O | class 7 | Shifted along the β-spine axis and lateral shift between sheets |
| SCT | H-LLQGLV-OH | A | 29WR | 5 mg ml<sup>-1</sup>, dissolved in 30 v/v% ethanol, left to evaporate from an eppendorf with a hole in it), 37 °C | A<br>B<br>C<br>D | H<sub>2</sub>O, ethanol, TFA | class 6 | Shifted along the β-spine axis and lateral shift between sheets |
| GHRH | ac-LLQDIM-NH<sub>2</sub> | A | 29VU | 5 mg ml<sup>-1</sup>, CD sample, shaken overnight at 37 °C, pH 4 | A | H<sub>2</sub>O | class 1 | Lateral shift between sheets |

<sup>a</sup> Amyloid classes are defined based on Eisenberg classification<sup>51</sup>
<sup>b</sup> Adjacent β-sheets may laterally shift relative to one another, altering inter-sheet packing, while β-strand dimers may translate along the $\beta$ -spine axis, interrupting its axial continuity.

#### PACAP

The PACAP-derived APRs (YLAAVL) illustrates how a single sequence can adopt several distinct structural assemblies. In polymorph A of APR^PACAP^, antiparallel β-sheets assemble according to the class 6 architecture. **(Fig. 5/A)** A hydrophobic core is formed by Leu, Ala and Val residues, while the hydroxyl groups of Tyr residues form a hydrogen bond network with chain termini and water molecules. In contrast, polymorph B of APR^PACAP^ **(Fig. 5/B)** cannot be classified within the original steric-zipper class scheme, as it contains four distinct strands per translational unit. This architecture corresponds to class 4.1 in the extended classification of Brumshtein et al.^51^ Beside various hydrophobic interactions, a hydrogen bond system formed by Tyr side chains, water molecules and chain termini can be observed in this crystal form too – however, water molecules here can also be found in the interfaces between sheets. In the third zipper, the N-terminally protected variant of YLAAVL (_ac_APR*^t^*^PACAP1^) **(Fig. 5/C)** preserves the packing mode of its non-protected counterpart, belonging to class 6, displaying an identical side-chain interaction topology within the steric zipper interface.

#### VIP

Although the parallel β-sheet crystal structure of APR^VIP^ **(Fig. 5/D)** formally corresponds to a class 4 steric zipper, it deviates from the canonical alternating side-chain pattern, as Tyr1, Leu2, and Ser4 align on one face, while Asn3, Ile5, and Leu6 occupy the opposite face. This atypical arrangement (**SFig. 16**) enables simultaneous formation of a shared hydrophobic core and an extended H-bond network involving polar residues, termini, and interfacial water molecules.

#### PHM

The crystal structures of PHM-derived peptides (YLESLM) highlight how minimal chemical perturbations can redirect the structural landscape. The unprotected peptide (APR^PHM^) forms a class 1 parallel steric zipper (**Fig. 5/E**) with two distinct regular interfaces: one formed by Glu3, Leu5 and chain termini and another by Leu2, Ser4, Met6 and two water molecules at the interface. Due to a shift between adjacent β-sheets, Tyr1 forms a third, small interface with Leu2. **(Fig. 5/E)** In contrast, the protected variant (_ac_APR^PHM^) adopts a highly unusual backbone conformation (**Fig. 5/F**), in which several torsion angles fall outside the canonical β-region on the Ramachandran plot. (**SFig. 16**)While in canonical β-strands the planes of the peptide bonds are perpendicular to the direction of the strand, here the backbone undergoes an approximately 90° rotation along the chain, enabling H-bonding not only within β-sheets but also between backbone atoms and side chains across neighbouring sheets. Although only a limited segment of the chain participates in classical steric zipper-like interactions, this antiparallel arrangement can be classified as a class 7 steric zipper.

#### SCT

While _ac_APR_am_^SCT^ yielded microcrystals that were not suitable for data collection, APR^SCT^ **(Fig. 5/G)** crystallized as antiparallel β-sheets in a class 6 steric zipper arrangement. This structure exhibits unusual features, as the asymmetric unit contains four distinct strands (A, B, C, and D) that are pairwise conformationally similar (A–C and B–D), yet laterally displaced and not coplanar, introducing an additional level of packing heterogeneity.

#### GHRH

APR of GHRH with both free and protected termini yielded microcrystals, but only crystals of the latter (_ac_APR_am_^GHRH^) were suitable for data collection. _ac_APR_am_^GHRH^ **(Fig. 5/H)** adopts a class 1 parallel β-sheet architecture, in which favourable hydrophobic interactions flank both the inter- and intrasheet H-bond networks. The odd-numbered face (Leu1, Gln3, and Ile5) interacts with the corresponding face of a neighbouring sheet, allowing Gln side chains to form hydrogen bonds at one interface. On the even-numbered face (Leu2, Asp4, and Met6), the protonated Asp4 side chains, together with water molecules, establish a hydrogen-bond network at the opposite interface. Due to terminal protection and the resulting absence of terminal charges, the peptide termini do not participate in the typical interactions seen in many amyloid structures. Instead, the acetyl group contributes to hydrophobic contacts, while the C-terminus forms multiple hydrogen bonds with backbone carbonyls.

Taken together, the diverse sequence composition of PACAP-family APRs suggests that they do not encode a single amyloid fold, but instead populate multiple steric zipper classes, including both canonical and atypical architectures, reflecting a dynamic energy landscape governed by sequence features and environmental conditions.

### 5. Amyloid formation propensity of the PACAP-glucagon family is evolutionary encoded

A central evolutionary question arising from our findings is whether the aggregation-prone nature of the receptor-recognition region and its capacity for functional amyloid formation evolved independently in the PACAP and GCG families or were inherited from a common ancestral peptide and maintained throughout vertebrate evolution. To address this question, we conducted a comprehensive literature survey to identify the amino acid sequences **(S.Table 1**) and their corresponding genomic coding regions of the known members of the PACAP–GCG family. The selected species broadly represent the major vertebrate lineages. Particular attention was given to the conservation of the gatekeeper/heparin-binding regions and APRs within the first 27-residue-long bioactive core. **(S.Table 2**) Based on the classification proposed by Cardoso et al.^52^ and supported by our hierarchical clustering at both the amino acid **(SFig. 17**) and nucleotide levels **(SFig. 18**), the DNA and peptide sequences derived from different species were grouped according to their similarity into the corresponding peptide families. Within each family, we subsequently analysed position-specific amino acid conservation and the diversity of synonymous codons encoding the observed residues. **(SFig. 19**) To quantify positional amino acid conservation, we calculated a normalized Shannon-based conservation index (CI, where 0 < CI < 1), which reflects the degree of residue conservation at each position across the analysed sequences. To obtain length-normalized values, the position-specific conservation index values calculated for each residue were summed and divided by the length of the analysed sequence (avg. CI_1-27_). **(Fig.6 C/D, SFig. 20**)

Overall, the PACAP–GCG peptide families display a high sequence conservation. The lowest average CI_1-27_ was observed for SCT (avg. CI_1-27_ = 0.78), whereas PACAP displayed the highest level of conservation (avg. CI_1-27_ = 0.97). The genomic analyses reveal extensive synonymous codon diversification: sequences encoding highly conserved peptides show substantial nucleotide-level variation while preserving identical or nearly identical amino acid sequences. Highly conserved residues within the same peptide families frequently exploit nearly the entire synonymous codon space. For example, in VIP family the fully conserved Leu27 residue is encoded by five of the six possible leucine codons among the analysed species (n = 13). **(SFig. 19**) Importantly, despite the prominent dual role of the receptor-binding/APR segment, it is not more conserved than the rest of the polypeptide hormone (avg. CI_1-27_ ≈ CI_21-27_) suggesting that its functional role is maintained without strict sequence conservation.

Despite sequence variability, aggregation propensity analysis reveals a conserved APR pattern (**Fig. 6**, **SFig. 20**). While species-specific mutations locally modulate aggregation tendencies, residues 21–27 consistently emerge as the dominant aggregation hotspot across all species and peptide families (**Fig. 6A**), in agreement with the experimentally identified APR in human peptides (**Fig. 4**). Notably, the average aggregation propensity of the human APR (avg. AggP_21–27_) is comparable to, or in some cases lower than, that observed in corresponding regions of other species (**Fig. 6**, **SFig. 20**). Given that this region has been experimentally confirmed to drive aggregation in human peptides, its equal or higher propensity in other species suggests that aggregation is likely preserved across these lineages. Moreover, substitutions at non-conserved positions typically involve residues with similar aggregation-promoting properties, maintaining the intrinsic aggregation potential of the APR. **(Fig.6/B,C SFig.20**). Most mutations at equivalent positions within a given hormone family can be explained by single-nucleotide codon substitutions. (**S.Table 4/1-10**) However, more extensive changes are also observed, involving complete codon replacements, which may partly reflect limited sequence availability or cases of convergent evolution (*e.g.*, SCT). Overall, sequence variation appears constrained by both mutational accessibility and functional requirements, preserving receptor binding while maintaining aggregation propensity.

The positional distribution of charged gatekeeper residues differs between the GCG and PACAP families (**Fig. 6/A**), yet these positions are more strongly conserved than the APR regions themselves. Within the bioactive core, gatekeepers are consistently enriched at positions 3, 8/9, 12, 15, and 20/21, immediately preceding the APR, and show locally reduced aggregation propensity (AggP). In the GCG family, these positions are dominated by negatively charged Asp/Glu residues, whereas in the PACAP family they are predominantly occupied by positively charged Arg/Lys residues associated with observed heparin sensitivity. This difference may explain why PACAP-family aggregation depends on heparin, whereas GCG-family aggregation is less heparin-dependent and more sensitive to pH, typically occurring under acidic conditions where gatekeeper residues are at least partially neutralized.^20,21,53,54^ Although many gatekeeper residues are conserved at the amino acid level, their encoding codons are often highly divergent **(SFig. 19).** This nucleotide-level variability enables multiple single-nucleotide substitutions that can generate alternative gatekeeper residues, including both conservative (Arg↔Lys, Glu↔Asp) and charge-altering changes. A representative example is position 8 in the PRP family, where successive single-nucleotide substitutions can yield multiple residue variants. (*e.g.*, GAG(E) →GAA(E) →AAA(K)→AGA(R)→AGT(S))

The emergence of new charged gatekeepers or charge-altering substitutions is particularly common in species with multiple copies of the exon encoding homologous peptides. In such cases, one copy typically preserves the ancestral gatekeeper pattern, while the other diverges, altering charge distribution and aggregation propensity. This pattern is reflected in pairwise identity heatmaps **(SFig. 17, 18),** where duplicated peptides within a species often show lower similarity than orthologs from related species. Thus, exon duplication provides evolutionary flexibility, enabling functional conservation alongside diversification, as illustrated by the GCG exon serving as a template for the emergence of GLP-1 and GLP-2 families. ^26^

The earliest traceable members of the PACAP–glucagon superfamily originate from protochordates that diverged from the vertebrate lineage ∼600 million years ago. In the tunicate *Chelyosoma productum* (**Fig. 6/B**), two full-length cDNAs encode four 27-residue peptides closely related to vertebrate PACAP–GCG family members **(Fig. 1**).^55^ Notably, one peptide differs from human PACAP-27 only by a single residue, indicating that the PACAP-encoding exon represents an ancient and highly conserved element of the superfamily, preserved from early chordates to humans^25,52,56,57^ (**Fig. 6/D**). In contrast, another exon in the tunicate gene likely corresponds to the ancestral precursor of the PRP and GHRH families (commonly referred to as PRP-like or GHRH-like/GRF-like, i.e., growth hormone–releasing factor) and shows greater sequence variability (**Fig. 6/C**), although a shared evolutionary origin is still supported (**SFig. 17, 18**). Sequence analyses further reveal that the tunicate PRP-like peptide shows distinct relationships at different levels, clustering with the PRP/GHRH lineage at the amino acid level **(SFig. 17, S.Table 3**), but with GCG sequences at the nucleotide level. **(SFig. 18, S.Table 3**) This suggests that it may represent an evolutionary link between the PACAP and GCG families.

Consistent with the one-to-four rule^28,58–60^ describing early gene expansion, the modern PACAP–GCG peptide families can be traced back to these tunicate peptides, which approximate the ancestral state of the superfamily. Having established the aggregation properties of human peptides (*h*), tunicate-derived sequences (*t*) were examined experimentally to determine whether this propensity was already present in the ancestral lineage and how it differs from their human counterparts.

### 6. Tunicate-derived (*t*) ancestral peptides form amyloids with weaker heparin dependence than their human (*h*) counterparts

First, the three distinct APR hexapeptides corresponding to the free C-terminal residues 21–27 encoded by the two tunicate PACAP cDNAs were investigated using the same experimental framework as their human counterparts. Amyloid aggregation of _ac_APR*^t^*^PACAP^ ^1^ (ac-YLAAVL-OH) was expected, as it differs from human constructs (APR^PACAP^, _ac_APR_am_^PACAP^) only in terminal protection and adopts a similar steric-zipper morphology (**Fig. 5/A**). Another PACAP-related hexapeptide, _ac_APR*^t^*^PACAP^ ^2^ (ac-YINALL-OH), encoded by the homologous exon, as well as the _ac_APR*^t^*^PRP-like^ ^1/2^ (ac-FLQWLM-OH), identical in both exons, formed microcrystals and showed a pH-dependent transition from disordered to β-sheet conformations by CD spectroscopy **(SFig. 21),** though diffraction-quality crystals could not be obtained. Notably, _ac_APR^tPRP-like1/2^ more closely resembles the human APR^GCG^ (FVQWLM) than APR^PRP^ (HLQSLV) at both the genomic and amino acid levels. This relationship is reflected in the CD spectra, which display a characteristic negative–positive band pair around 220 nm, indicative of aromatic Phe–Trp interactions previously linked to aggregation in GCG-family peptides.^14^

As a next step, we investigated the aggregation propensity of the full-length 27-residue peptides, as *t*PRP-like 1 and *t*PACAP 1 represent ancestral forms from which all human peptide family members evolved but lack the C-terminal elongation present in their human counterparts. **(Fig. 6/E, SFig. 22, 23)** Both tunicate-derived peptides rapidly formed amyloid in the presence of heparin across a broad pH range (pH 2 to 10), where the net peptide charge remains positive throughout. This confirms that peptide–heparin interactions facilitate self-assembly by charge neutralization, as observed in human peptides. Strikingly, unlike their human analogues, both tunicate peptides were capable of forming amyloid fibrils in the absence of heparin over a more limited pH range (pH 5–10). (**Fig. 6/F**, **SFig. 22, 23**) Below pH 5, the net charge is highly positive (+4 to +7), and in the absence of negatively charged heparin, this uncompensated electrostatic repulsion prevents peptide–peptide interactions, highlighting the dominant role of electrostatics in aggregation.

However, charge control alone does not explain why tunicate peptides aggregate without heparin, whereas human PACAP-38, PRP, and GHRH do not. The tPRP sequence shares low similarity with the bioactive cores of *h*PRP and *h*GHRH (48% and 59%; **Fig. 1/B,C**), but higher homology to *h*GCG (64%), which aggregates without heparin.^13,21,61^ Accordingly, the heparin-independent aggregation of *t*PRP, together with the observed Phe–Trp interaction in _ac_APR^tPRP-like1/2^, links it to the GCG lineage at the physicochemical level rather than the genomic level. These features appear preserved in the GCG branch but lost in the PRP/GHRH lineage.

In contrast to *t*PRP, the *t*PACAP1 sequence is highly similar to the *h*PACAP-27, sharing 96% sequence identity and differing at only a single position (Asn15 instead of Lys15). Although this substitution may introduce an additional gatekeeper residue in the human PACAP variants, we propose that the major difference in the observed heparin-limited aggregation arises from the Lys/Arg-rich C-terminal extension of hPACAP-38. While the shorter *t*PACAP sequence, resembling *h*PACAP-27, can aggregate even in the absence of heparin, the extended C-terminus of *h*PACAP-38 (–GKRYKQRVKNK) introduces an additional, extremely positively charged gatekeeper region that likely acts as an additional heparin-binding motif, restricting peptide self-association and amyloid formation primarily to conditions where heparin is present (**SFig. 1**).

This example highlights that in peptide lineages with C-terminal extensions beyond the 27-residue bioactive core (*e.g.*, PACAP, PRP, GHRH - **Fig. 1, S.Table 1**), these regions, although not directly involved in receptor activation, can modulate functional amyloid formation. These extensions are enriched in gatekeeper residues—Lys/Arg in PACAP, Glu/Arg/Gly in GHRH, and Glu/Asp/Gly/Pro in PRP—suggesting a role in limiting aggregation. A similar effect was observed in the exendin family, a GLP-1–related lineage in reptiles, where the proline-rich C-terminus of exendin-4 acts as a β-sheet breaker, and amyloid formation occurs only upon its truncation.^62^

## CONCLUSIONS

Phylogenetic and comparative studies suggest that ancestral class B1 GPCR peptide hormones were highly versatile, affecting multiple signalling systems. With increasing organismal complexity, gene duplication enabled functional diversification, while preserving core features essential for hormone function.^28^ In this context, our results show that although receptor–ligand interactions evolved plastically, the aggregation profile of the PACAP–glucagon peptide family remained remarkably conserved, likely reflecting the functional importance of amyloid formation in hormone storage. Importantly, sequence evolution predominantly involved substitutions accessible through single-nucleotide codon changes, maintaining intrinsic aggregation propensity while allowing receptor specificity to diversify.

A key determinant of this balance is the co-evolution of aggregation-prone segments with charged gatekeeper residues. In the PACAP family, Lys/Arg-rich regions suppress spontaneous self-association through electrostatic repulsion but enable aggregation in the presence of polyanionic cofactors such as heparin, which neutralize these interactions. This establishes a regulatory mechanism in which aggregation is conditionally activated. In contrast, such motifs are less dominant in the glucagon family, contributing to differences in aggregation behaviour. Furthermore, optional C-terminal extensions enriched in gatekeeper residues introduce an additional regulatory layer, modulating aggregation propensity without affecting receptor activation.

Molecular simulations and experimental observations together suggest that glycosaminoglycans, such as heparin, play a dual role in this process. Beyond electrostatic charge neutralization, their negatively charged polymeric structure acts as a scaffold that recruits positively charged peptides, locally increasing concentration and promoting amyloid nucleation. A similar recruiting function of heparin has been described in mast cells, where it facilitates the maturation and storage of positively charged proteases^63^, as well as in the regulation of certain inflammatory processes^64^. Notably, the evolutionary origin of heparin can be traced back to protochordate tunicates^65,66^, implying that glycosaminoglycans were already present during the early emergence of peptide hormone families.

Despite the clear evolutionary relationship among PACAP-related peptides, secretin (SCT) occupies a distinct position within the family. Its evolutionary origin remains unclear, as SCT has so far only been identified in tetrapods, with the earliest representatives in amphibians. Available sequences differ markedly from other family members at both the amino acid and, more prominently, the genomic level, and even within the SCT family, many codon transitions would require multiple nucleotide substitutions, indicating a greater evolutionary distance. Consistent with this divergence, SCT also exhibits distinct aggregation behavior: while the isolated APR^SCT^ segment forms amyloid-like structures, the full-length peptide does not aggregate under comparable conditions, even in the presence of heparin. Together, these observations suggest that SCT represents a functionally and evolutionarily distinct branch, and that similarities with PACAP-related peptides may reflect convergent rather than shared evolutionary origins.

Overall, the PACAP–glucagon peptide family emerges as a class of structurally adaptable, “chameleon-like” molecules capable of adopting multiple conformational states depending on their molecular environment. **(Fig. 7**) While disordered in solution and helical upon receptor binding, they can transition into β-sheet–rich amyloid structures in the presence of appropriate cofactors. This conformational plasticity and intrinsic aggregation propensity appear to be evolutionarily constrained features that cannot be eliminated without compromising function. Recognizing this constraint is particularly important for drug development and long-term product safety, especially given the rapidly expanding clinical use of class B1 GPCR agonists such as GLP-1–based anti-obesity therapeutics including semaglutide.^67^

## MATERIALS AND METHODS

### Peptide synthesis

Full-length PACAP-related peptides and their corresponding APR hexapeptides were synthesized using our in-house developed flow chemistry-based solid-phase HPPS-4000 flow peptide synthesizer (METALON Ltd.) a modified version of the standard Jasco LC-4000 series high-performance liquid chromatography (HPLC) system. The primary modification consists of an additional valve integrated into the PU-4180 HPLC pump, enabling precise control of solvent flow. The entire process was fully automated using ChromNAV2 software (version 2.04.06). During the synthesis the Fmoc/tBu strategy was used.^68,69^ Preloaded Wang TentaGel resin (containing the first C-terminal amino acid) was used for peptides with a non-protected C-terminus, while RAM TentaGel S resin (0.24 mmol/g, Rapp Polymere GmbH) was applied for peptides with a protected C-terminus. Coupling reactions were performed using OxymaPure and DIC as reagents, with DMF as the solvent. For Fmoc-deprotection, a solution of 20% (v/v) piperidine in DMF was used. The resin was washed with DMF between each step. Protected and special amino acids were dissolved in N-methyl-2-pyrrolidone (NMP) containing 1 equivalent of OxymaPure. The activating agent, diisopropylcarbodiimide (DIC), was added immediately prior to coupling and was injected via the autosampler. The reactions were carried out at 80 °C under a pressure of 8–12 maintained by a backpressure regulator MPa. Peptides containing Met, His, or Arg residues were cleaved from the resin using a mixture of 0.25 g phenol, 2.5 v/v% ethane-1,2-dithiol, 5 v/v% thioanisole, 5 v/v% water, 1.2 v/v% triisopropylsilane, and 86.3 v/v% TFA. For other peptides, cleavage was performed using a mixture of 2.5 v/v% triisopropylsilane, 2.5 v/v% water, and 95 v/v% TFA. Cleavage reactions were carried out at room temperature under continuous stirring for 4.5 h or 3 h, respectively. TFA was removed using a rotary vacuum evaporator, and the peptides were precipitated in cold diethyl ether. After sedimentation, the ether was decanted and the peptide pellet was washed again with fresh ether. This washing cycle was repeated three times, followed by vacuum drying. The crude oligopeptides were dissolved in 5:95 v/v% ACN:H₂O and filtered through a PTFE membrane (45 μm). Prior to filtration, the pH of APR hexapeptides containing Asp or Glu residues was adjusted to pH 7 to increase solubility. APR hexapeptides were purified by reverse-phase HPLC using a C12 column (Jasco LC-2000Plus system equipped with a Jupiter 10 μm Proteo 90 Å column, 250 × 10 mm), whereas full-length peptides were purified using a C18 column (Kinetex 5 μm EVO C18, 100 Å, 150 × 21.1 mm) with gradient elution (ACN/water containing 0.1% TFA). Collected fractions were immediately frozen and lyophilized. Aliquots of the APR peptides were reserved for crystallization. Analytical purity was confirmed by analytical HPLC (Jasco LC-2000 system equipped with a Jupiter 4 μm Proteo 90 Å column, 250 × 4.6 mm) using gradient elution (ACN/water containing 0.1% TFA, 5–95% B over 30 min). Mass spectra were recorded on a Bruker Esquire 3000+ tandem quadrupole mass spectrometer equipped with an electrospray ionization source.

### Amyloid sample preparation of the APRs

Lyophilized hexapeptides with and without terminal protection were dissolved in distilled water at a nominal concentration of approximately 5 mg mL^-1^. The exact peptide concentrations could not be accurately determined due to limited solubility. In addition, for the LLQDIM, LLQGLV, and HLQSLV peptides, concentration determination by UV spectrophotometry at 280 nm was not possible because these sequences lack aromatic chromophores. The pH of each sample was adjusted using 0.01, 0.1, or 1 M NaOH and HCl solutions, measured with an Orion Star A211 pH meter (Thermo Scientific). Samples were continuously stirred using a magnetic stirrer (500–600 rpm) and incubated at 37 °C for 24 h in the case of human APRs and for 72 h in the case of tunicate APRs.

### Amyloid sample preparation of the full-length peptides monitored by Thioflavain-T (ThT) fluorescence

The peptides were dissolved in distilled water and filtered through a PTFE membrane (45 μm). The peptide concentrations were determined using a NanoDrop Lite spectrophotometer (Thermo Scientific) at 280 nm, applying the appropriate molar extinction coefficients for each peptide. (ε_hPACAP_ = 6990 M^-1^ cm^-1^, ε_tPACAP1_ = 4470 M^-1^ cm^-1^, ε_hVIP_ = 2980 M^-1^ cm^-1^, ε_hPRP_ = 1490 M^-1^ cm^-1^, ε_tPRP-like_ = 8480 M^-1^ cm^-1^, ε_hPHI_ = 1490 M^-1^ cm^-1^, ε_hGHRH_ = 2980 M ^-1^ cm^-1^. SCT is lack of aromatic chromophores thus the concentration was determined at 214 nm with ε_SCT_=35821 M^-1^ cm^-1.^ The samples were subsequently diluted to concentration of 0.42 mM. The pH of each sample was adjusted using 0.01, 0.1, or 1 M NaOH and HCl solutions. No buffer systems were used, as the presence and composition of buffers can influence amyloid aggregation. In addition, buffer components interfere with CD measurements (180-190 nm) and complicate AFM imaging due to crystallization of buffer salts during sample drying. Incubations were performed in black-walled, flat-bottom 96-well microplates (Greiner Bio-One). Each well contained 160 µL of 0.42 mM peptide solution (or distilled water for blank samples), 20 µL of pH-adjusted 50 µM thioflavin T (ThT) stock solution, and either 4.25 µL of enoxaparin sodium (as synthetic heparin) solution with 15.75 µL distilled water (for heparin-containing samples) or 20 µL distilled water (for heparin-free samples). The enoxaparin sodium solution was obtained from a commercially available prefilled syringe (Clexane^®^, Sanofi-Aventis), containing 20 mg enoxaparin sodium in 200 µL. The final peptide concentration in each well was 0.35 mM, corresponding to approximately 1 mg mL^-1^. Thioflavin T (ThT) stock solutions (Acros Organics, Thermo Fisher Scientific) were prepared in distilled water, with the pH adjusted between 2 and 10 in increments of one pH unit. The solutions were filtered through a 0.45 µm membrane filter. The exact ThT concentration was determined by UV spectroscopy (Jasco V-660 spectrophotometer) at 412 nm using an extinction coefficient of 31 600 M^-1^ cm^-1^. The stock solutions were stored in the dark at 4 °C. The plate was hermetically sealed to prevent evaporation. A SpectraMax iD3 microplate reader (Molecular Devices) was used to control the experimental conditions (37 °C, orbital shaking at medium intensity) and to record fluorescence data for 72 h. The excitation and emission wavelengths were set to 445 and 490 nm, respectively. ThT fluorescence intensity was measured from the bottom of the microplate using medium photomultiplier tube (PMT) sensitivity with an integration time of 400 ms. Under these conditions, the fluorescence signal of some amyloid-forming samples exceeded the upper detection limit of the detector. In such cases, the last correctly detected maximum value was assigned to subsequent data points, resulting in an apparent plateau phase. Blank samples contained all components except the peptide. For each condition (with or without heparin), blank measurements were performed in triplicate, whereas peptide-containing samples were measured once. The fluorescence intensity of each peptide sample was normalized to the mean signal of the corresponding blank measurements at each time point. The resulting blank-normalized fluorescence increase was plotted as a function of time.

### CD measurements of the APR hexapeptides and the full-length hormone peptides

Circular dichroism (CD) measurements were performed using a JASCO J-1500 spectropolarimeter. The temperature of the cuvette holder was controlled by a Peltier-type heating system. Data acquisition and processing were carried out using JASCO Spectra Manager v2.0 software. Aliquots (40 µL) taken directly from the incubated samples were measured in 0.1 mm path-length quartz cuvettes at 25 °C. Each spectrum represents the average of three scans recorded with a spectral scanning speed of 50 nm min⁻¹, a bandwidth of 1 nm, and a step resolution of 0.2 nm over the wavelength range of 180–260 nm (far-UV region). All spectra were baseline-corrected by subtracting the solvent (with or without heparin) spectrum and smoothed using the Savitzky–Golay method with a convolution width of seven. The raw ellipticity values (deg°) were plotted as a function of wavelength. During the interpretation of the CD spectra, conversion of the raw ellipticity values to mean residue molar ellipticity units was not performed. This was due to uncertainties in the effective peptide concentration caused by aggregation during incubation, as well as increased light scattering arising from sample turbidity. Furthermore, in our analysis the overall shape of the CD spectra was considered more informative than the normalized intensities of spectral minima and maxima.

### Deconvolution of full-length peptide CD spectra

For spectral deconvolution, CD spectra of the full-length peptides were normalized by the number of amino acid residues. In the case of peptides with an amidated C-terminus, the residue number was increased by one to account for the additional amide group. Normalization by peptide concentration was not applied, as all samples were prepared from the same initial concentration. CD measurements in which the absolute ellipticity was below 5 deg° were excluded from further analysis, as such low signal intensity indicated substantial peptide loss due to precipitation and the resulting decrease in effective concentration. For the deconvolution analysis, only the spectral region between 190 and 260 nm was used, since measurements at lower wavelengths frequently showed increased noise. The CD spectral matrix was decomposed by CCA+^70^ (Convex Constraint Algorithm) v2.42 software using an increasing number of components. Evaluation of the resulting basis spectra indicated that a seven-component model provided the most meaningful description, yielding basis curves corresponding to the canonical CD signatures of disordered, α-helical, and β-sheet secondary structures, along with four additional spectra that resemble variants of these canonical spectral profiles.

### Fourier-transform infrared spectroscopy (FTIR) measurements

FTIR measurements of the incubated human APR hexapeptides were performed using a Bruker Equinox 55 FTIR spectrometer equipped with a bio-ATR (attenuated total reflectance) cell with a ZnSe internal reflection element. The ZnSe photoelastic modulator of the instrument was set to 1600 cm^-1^, and an optical filter with a transmission range of 1900–1200 cm^-1^ was used to enhance sensitivity in the amide I–II spectral region. The MCT (mercury–cadmium–telluride) detector was cooled with liquid nitrogen. Each FTIR spectrum was recorded by averaging 128 scans in the range of 4000–850 cm^-1^ with a spectral resolution of 4 cm^-1^ using an aperture of 3000 µm. IR spectra were calculated from the single-channel DC spectra. Baseline correction was applied to each measurement by subtracting the corresponding blank solvent spectrum. Data processing and export were performed using OPUS 6.5 software.

### Atomic Force Microscopy (AFM)

Five microliters of incubated APR and full-length peptide samples were diluted 20-fold with appropriately pH-adjusted water, immediately deposited onto freshly cleaved mica surfaces, and dried overnight in a vacuum desiccator. AFM imaging was performed on the dried samples; however, in some cases small residual water patches were still observed on the surface. AFM imaging was performed using a FlexAFM microscope system (Nanosurf AG) operating in dynamic mode, controlled by Nanosurf Control software C3000 version 3.10.4. Measurements were carried out using Tap150GD-G cantilevers (BudgetSensors Ltd.) with a nominal tip radius of less than 10 nm. Samples selected for AFM analysis included those incubated under biologically relevant pH conditions (pH 5 and pH 7) and samples in which either CD spectroscopy or ThT fluorescence measurements indicated amyloid formation. Low-resolution pre-screening scans were performed prior to high-resolution imaging to identify fibril-like structures and to avoid areas with large surface height variations. When optically dense aggregates were observed on the camera, AFM images were collected in close proximity to these regions. In the absence of such aggregates, pre-screening scans were used to accelerate the mapping of the sample surface. If low-resolution pre-screening of three independent surface locations indicated the presence of amyloid fibrils, high-resolution data were subsequently acquired, and representative datasets were selected for illustration. If fibrillar structures were not detected in the first three screened regions, three additional pre-screening scans were carried out, followed by the acquisition of at least one high-resolution micrograph. It should be noted that the presence of fibrillar structures is generally easier to confirm than their absence. This consideration also applies to cases where fibrils were not detected by AFM, although CD or ThT measurements suggested amyloid formation, potentially including false-positive indications. Micrographs with a scan size of 10 µm × 10 µm and a resolution of 512 pixels per line were recorded. In several cases, additional images of selected regions of interest were acquired at higher resolution. AFM data were processed and micrographs exported using Gwyddion 2.62 software.

### Approximate protonation state distribution and net charge calculations

The protonation state distribution (microspecies distribution) and net charge of the peptides were calculated over the pH range of 2–10 using a site-specific acid–base equilibrium model. Each ionizable group (N-terminus, C-terminus, and ionizable side chains) was treated as an independent titratable site characterized by its intrinsic pKₐ value. Approximate net charges of the peptides were calculated using generalized pKₐ values: Asp (3.9), Glu (4.2), His (6.0), Cys (8.3), Tyr (10.1), Lys (10.5), Arg (12.5), N-terminus (8.0), and C-terminus (3.1). This approximation is justified in the present context, as the peptides are short and predominantly disordered in solution, as indicated by CD measurements, and thus less likely to exhibit strong site–site interactions, electrostatic coupling, and potential environment-dependent shifts in pKₐ values. For each pH value, the fractional protonation of individual sites was calculated using the Henderson–Hasselbalch formalism. For acidic groups, the deprotonated fraction was computed as:

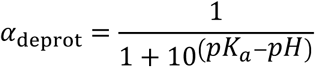

while for basic groups, the protonated fraction was calculated as:

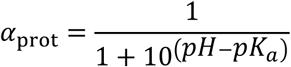

Microspecies populations were determined by enumerating all possible protonation states of the peptide and calculating their probabilities as the product of the corresponding site-specific protonation or deprotonation fractions. The net charge of each microspecies was obtained by summing the charges of all ionizable groups in their respective protonation states. The overall net charge at a given pH was calculated as the population-weighted average over all microspecies.

### Hydrophobicity and hydrophobic moment calculations

The average hydrophobicity (⟨H⟩) of each peptide was calculated using the Eisenberg hydrophobicity scale^71^, by taking the arithmetic mean of the residue-specific hydrophobicity values along the sequence. The hydrophobic moment (μH), describing the amphipathic character of the peptides, was calculated assuming an ideal α-helical geometry with a periodicity of 100° per residue. Each amino acid was represented as a vector with magnitude corresponding to its Eisenberg hydrophobicity value and direction defined by its angular position in the helix. The vector components were summed over the entire sequence, and the magnitude of the resulting vector was normalized by the number of residues:

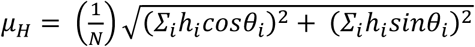

where *h_i_* is the hydrophobicity value of residue ^i^, θ*i* is its angular position, and N is the sequence length.

### Molecular dynamic simulation of PACAP-related peptides and their complexes with heparin

The Cryo-EM structure of human vasoactive intestinal polypeptide receptor 2 (VIP2R) in complex with PACAP27 (PDB ID: 7VQX^72^) was downloaded from the protein data bank (PDB) and the 27 long helical PACAP peptide was extracted from it using PyMol. (v2.0 Schrödinger, LLC.) As the receptor-bound form of all investigated peptides are α-helical^27^, all other simulated peptides were constructed from this structure using PyMol’s mutation tool. Conformers of the mutated residues were selected from the backbone-dependent rotamer library of PyMol with visual assessment. A C-terminal amide protection was also constructed for all peptides. During all simulations, arginines, lysines and unprotected N-termini were positively charged, aspartic and glutamic acid side chains were negatively charged.

The peptides were simulated by themselves and, separately, with a 4-sugar unit heparin molecule fragment with molecular dynamics (MD). The single peptide simulations were started from the aforementioned structures using the GROMACS 2024.4 software^73^. The AMBER99SB-star-ILDNP^74^ force field was used with the OPC^75^ water model. All systems were placed at the centre of a dodecahedral box with a side-length of 7.179 nm. Chloride ions were used to neutralize the peptides, but no additional salt was present in the solution. Four energy minimization steps were performed with the conjugate gradient method using 1000, 500, 100 and 0 kJ·mol^-1^·nm^-2^ positional constraints subsequently. Temperature equilibration was performed using four, 400 ps long NVT runs with the v-rescale thermostat and the same positional constraints. Then, after a 100 ps long NPT pressure equilibration step (using the Parrinello-Rahman barostat), a 1 μs long production NPT simulation was run for all peptides. Step size was set to 2 fs for all simulations.

After the single peptide production runs, water was removed from the trajectories, and the trajectory was centred to the peptide. Helicity was assessed using our in-house developed Gaussian Mahalanobis Mean (GMM) technique^45^. This metric measures the density of (Φ, Ψ) dihedral angle pairs in the α-helical region of the Ramachandran map. It reports this density as a number between 0 and 1, where 0 means the lowest possible helicity, while numbers near 1 indicate a highly helical peptide. Clustering analysis was performed on the first and last 200 ns parts of the full trajectory using the gromos clustering algorithm, a cutoff of 0.2 nm and a Δt of 100 ps. The most populated cluster’s central structure was used for the simulations that involved the 4-sugar unit heparin fragment.

The solution structure of an 18-sugar unit heparin (PDB ID: 3IRI^76^) was downloaded from the PDB and a 4-unit long part was extracted from it using PyMol. Missing hydrogens were added to the structure (except for the deprotonated sulfate and carboxylate moieties) and it was energy minimized using the xtb version 6.4.1 software and the GFN2-xTB method^77^. The constructed molecule has a net charge of-8. After minimization, the ACPYPE^78^ (v2023.10.27) software was used to construct the molecule’s AMBER force field parametrization. The parametrized molecule was incorporated into the AMBER99SB-star-ILDNP force field, and a standard MD simulation was performed using the same minimization and equilibration protocol as for the single peptide runs. The last 200 ns of the simulation was, again, clustered to obtain a representative equilibrium structure.

Systems were built using the representative peptide structures and representative, 4-unit heparin fragments in a 1:1 molecule ratio. The corresponding peptide was placed at the centre of the box, while the heparin molecule was placed at the most distant face of the box. Both molecules were oriented so that their principal axes aligned with the box sides. The box was a rectangular cuboid with side lengths of 5.5, 5.5 and 7.5 nm. The box was solvated, ions were added to neutralize the system, NVT and NPT equilibrated, and then a 1 μs long steered MD (SMD) simulation was performed using the Colvars^79,80^ plugin of GROMACS. During equilibration, a 1000 kJ·mol^-1^·nm^-2^ distance constraint was defined between the geometric centre of the heparin and the peptide with an equilibrium distance of 3.5 nm. Then, during the production SMD run, the equilibrium distance was slowly shifted from 3.5 nm to 0.5 nm, where the last distance constraint was enforced at 1 μs. The work performed by the moving constraint was recorded with the *outputAccumulatedWork* keyword set “on” in the Colvars configuration file.

For the parametrization of the 12-sugar unit heparin the structure of the molecule was assembled using RDKit^81^ (2025.03.06) and ETKDG v3^82^. After assembly, using the openff-nagl-models library (version 2025.9.0)^83^, AM1-BCC^84^ charges were approximated with NAGL charges^85^ using the openff-gnn-am1bcc-1.0.0 model. Bonded and nonbonded parameters were extracted from the openff-2.2.0.offxml file, encoding the parameters of the Sage force field^86^ (2024.04.18). Using in house Python scripts, the parametrization was transferred to the AMBER99SB-star-ILDNP force field, through new entries in the atomtypes.dat and ffnonbonded.itp files, as well as a new heparin.rtp file. The structure of heparin was exported as a gro file using the openff-interchange (version 0.5.2) library. Next, heparin alone was MD simulated using GROMACS using the above-described protocol for 1 μs and the last 200 ns of the trajectory was clustered. The centroid molecule of the most populated cluster centre was used for the simulation of the 3:1 = PACAP:HEP system.

In the 3:1 = PACAP:HEP system the molecular components were placed in a cube with a side length of 12 nm, so that the minimum pairwise distance between the components was maximized. This resulted in an effective concentration of 3 mM (calculated for the peptide). After that, a standard MD simulation with the already described protocol was performed. For the simulation of the 3 PACAP system (without HEP) this was also the starting structure, just with the HEP molecule removed. Intermolecular distances were calculated, after centring and putting molecules back into the box, using the *gmx pairdist* command.

### X-ray diffraction measurements and structure determination

Single-crystal X-ray diffraction data were collected at 100 K on a Rigaku XtaLAB Synergy-R rotating anode diffractometer using Cu Kα radiation or at beamline I03 at Diamond Light Source (YLNSIL). Data collection and reduction were performed using CrysAlisPro and AutoPROC^87^. The phase problem was solved by molecular replacement using idealized 5–6-residue-long β-sheet fragments or polyalanine chains derived from our previously determined amyloid structures as search models. Molecular replacement was carried out using Phaser (Phenix package)^88^ or, when sufficiently high-resolution data were available, Fragon^89^ (CCP4 package)^90^. Manual model building was performed in COOT^91^, followed by structure refinement using Phenix.refine^92^ and Buster^93^. Figures of atomic models were generated using PyMOL (v2.3.2) and ChimeraX (v1.8).

### Crystallization of hexapeptides

Ac-YLAAVL-OH: the peptide was dissolved in water at pH 7 at a concentration of 5 mg mL⁻¹, then diluted to approximately 1 mg mL⁻¹ with 40% (v/v) acetonitrile in 0.2 M acetate buffer (pH 4) and incubated at 37 °C. YLAAVL polymorph A: the peptide was dissolved in 0.1 M HEPES buffer (pH 7) at a concentration of 1 mg mL⁻¹ and incubated at 37 °C. YLAAVL polymorph B: the peptide was dissolved at approximately 1 mg mL⁻¹ in 0.2 M acetate buffer containing 30% (v/v) acetonitrile and incubated at 37 °C. YLNSIL: the peptide was dissolved at a concentration of 0.4 mg mL⁻¹ in 30% (v/v) methanol with 0.2 M acetate buffer (pH 3.77) and incubated at 37 °C. Ac-YLESLM-OH: the peptide was dissolved at 1 mg mL⁻¹ in water containing 30% (v/v) acetonitrile (pH 4) and incubated at 37 °C. YLESLM: the peptide was dissolved in 0.2 M acetate buffer (pH 4.67) at a concentration of 5 mg mL⁻¹ and incubated at 37 °C. LLQGLV: the peptide was dissolved at a concentration of 5 mg mL⁻¹ in 30% (v/v) ethanol in an Eppendorf tube with a perforated cap and incubated at 37 °C. Crystals grew on the wall of the tube upon solvent evaporation. Ac-LLQDIM-NH₂: crystals formed on the wall of the tube during overnight incubation at 37 °C from a 5 mg mL⁻¹ solution at pH 4, typically at the air–liquid interface.

### Comparative sequence analysis and evolutionary clustering

Pairwise sequence identity analyses were performed on both amino acid and nucleotide sequences corresponding to the N-terminal 27 residues of PACAP–glucagon family peptides. Peptide sequences and transcript IDs are contained in the Source Data and were used as input for all subsequent analyses. Pairwise identity was calculated by direct position-by-position comparison of the 27-residue peptide sequences and their corresponding 81-nucleotide coding regions. For each sequence pair, the number of identical characters at corresponding positions was determined. Heatmaps were generated using Python. For non-clustered heatmaps, sequences were displayed in the original order of the input dataset. Colour scales represented raw identical position counts (0–27 for amino acids; 0–81 for nucleotides). To assess similarity relationships among sequences, a distance matrix was computed from the percentage identity matrix using the transformation: “*distance*” = 100 − “*identity* (%)”, this produced a symmetric distance matrix with zero diagonal values. Hierarchical clustering was performed using the average linkage (UPGMA) method implemented in the SciPy library. The resulting linkage matrix was used to generate dendrograms representing relative sequence divergence. Dendrograms were plotted from the hierarchical clustering linkage matrix using SciPy’s dendrogram function. Branch lengths correspond to sequence divergence as defined by the distance metric (100 − *identity* %).

### Mutational pathway reconstruction of observed codons and mutational cost heatmap

Codon variability was analysed in a position-specific manner for each peptide family using aligned sequence data. At each position, only codons directly observed in the dataset were retained and mapped to their corresponding amino acids. Pairwise mutational distances between codons were quantified using the Hamming distance, defined as the number of nucleotide mismatches between two codons, corresponding to the minimal number of single-nucleotide substitutions required for interconversion. For positions with multiple observed codons, a hypothetical minimal mutational pathway was defined as the ordering of codons that minimizes the cumulative Hamming distance between consecutive elements. This was determined by evaluation of all possible permutations of the observed codons, selecting the sequence with the minimal total distance. These pathways represent computationally derived, hypothetical minimal routes within the observed codon space and do not imply evolutionary directionality or temporal ordering.

For each peptide family, position-specific mutational cost heatmaps were derived from the corresponding minimal mutational pathways. At each position, the mutational cost was defined as the maximum Hamming distance observed between any pair of consecutive codons along the optimal pathway, representing the largest single substitution step (bottleneck) within the inferred route. Accordingly, the heatmap reflects the maximum local mutational barrier within the hypothetical minimal pathway at each position irrespective of the presence of multiple alternative transitions involving fewer base changes.

### Aggregation propensity estimation

Aggregation propensity of the 27-residue-long biological active core of the PACAP–glucagon family was determined using AggreProt^47^. The publicly available code (https://github.com/loschmidt/aggreprot-predictor) was run locally. The training dataset and model weights were provided by the authors upon request.

### Shannon conservation index calculation

Positional amino acid conservation within each peptide family was quantified using a normalized Shannon conservation index.^94,95^ For each position of the analysed peptide region, the frequency of each amino acid was calculated from the aligned set of sequences. Let *p*_*i*_ denote the relative frequency of amino acid *i* at a given position The Shannon entropy

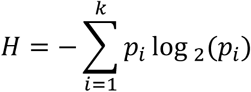

where *k* represents the number of distinct amino acids observed at that position. To allow comparison across positions, the entropy values were normalized by the theoretical maximum entropy corresponding to a uniform distribution of the 20 canonical amino acids:

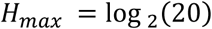

The normalized Shannon conservation index (CI) was subsequently calculated as:

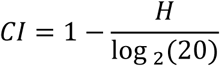

In this formulation, values approaching **1** indicate strong positional conservation, whereas values approaching **0** reflect higher variability. The conservation index was calculated independently for each peptide position based on the observed amino acid frequency distribution across all analysed sequences within the corresponding peptide family.

### Declaration on the use of generative AI

Generative AI (ChatGPT, OpenAI; GPT-5.3) was used to assist in language editing, improving clarity and readability of the manuscript, as well as generating and refining code for data processing and figure preparation. In the comparative sequence analysis and evolutionary clustering section, AI-assisted code was used, and all such code is provided in the accompanying Source Data. However, it was not involved in the generation or validation of primary scientific data. All results and conclusions were critically evaluated by the authors, who take full responsibility for the scientific content of the manuscript.

## DATA AVAILABILITY

The processed CD, FTIR, and ThT data generated in this study are provided in the Source Data file. Raw data of the FTIR and CD spectra, as well as the unprocessed AFM pictures, will be shared upon request by contacting the corresponding author. Novel structural coordinates have been deposited online in the Protein Data Bank under accession codes: 29VU, 29WR, 29VT, 29VS, 29VR, 29VQ. 29VP. 29VO. List of peptide sequences and transcript IDs used for the comparative sequence analysis, as well as the AI-generated codes applied in the analysis, are provided in the Source Data. Trajectories and cluster centres from the molecular dynamic simulations are also included in the Source Data.

## ACKNOWLEDGEMENTS

We thank Gergő Gyulai for providing access to the AFM instrument, and Joan Planas-Iglesias for supplying the files required for local execution of AggreProt. We acknowledge Diamond Light Source for providing beamtime at beamline I03. We thank Gergely Nagy for collecting data at Diamond Light Source. We also thank Dániel Borbély and Tamás Gabnai for their contributions to performing the experiments during their BSc/MSc studies.

## CONTRIBUTIONS

D.H. designed and coordinated the project. D.H., Sz.Sz., and Y.D. synthesized and purified the peptides. D.H., together with Sz.Sz., prepared the samples and performed CD and ThT measurements. D.H. measured the FTIR spectra and collected the AFM micrographs. D.H. processed the CD, ThT, AFM, and FTIR data. D.H. and Zs.D. crystallized the samples. Zs.D. measured, processed, and analysed the crystal structures. Sequence data analysis was performed by D.H. with the assistance of generative AI tools. Molecular dynamics simulations were set up and analysed by Zs.F. The manuscript was written by D.H., Zs.D., and Zs.F., with contributions from all authors. A.P. provided funding and extensive technical and instrumental support over the years.

