## Supplementary Data for "Evolutionarily Conserved Amyloid Aggregation in the PACAP Peptide Family Is Controlled by Heparin-Sensitive Lys/Arg Gatekeeper Residues"

### Supplementary Figures, Tables, Data

Dániel Horváth[1,2,*], Szebasztián Szaniszló[1,2], Zsolt Dürvanger[1,2], Zsolt Fazekas[2], Kim Hoang Yen Duong[2], András Perczel[1,2,*]

[1] HUN-REN–ELTE Protein Modeling Research Group, ELTE Eötvös Loránd University, Pázmány Péter sétány 1/A, H-1117 Budapest, Hungary.
[2] Laboratory of Structural Chemistry and Biology, ELTE Eötvös Loránd University, Pázmány Péter sétány 1/A, H-1117 Budapest, Hungary

**LIST OF CONTENT**

General description of Supplementary Figure 1-6

**S.Figure 1.** Monitoring of the amyloid formation for PACAP-38 in presence and absence of heparin

**S.Figure 2.** Monitoring of the amyloid formation for VIP in presence and absence of heparin.

**S.Figure 3.** Monitoring of the amyloid formation for PRP in presence and absence of heparin.

**S.Figure 4.** Monitoring of the amyloid formation for PHI in presence and absence of heparin.

**S.Figure 5.** Monitoring of the amyloid formation for GHRH in presence and absence of heparin.

**S.Figure 6.** Monitoring of the amyloid formation for SCT in presence and absence of heparin.

**S.Figure 7.** Color-coded bar chart analysis of the secondary-structure components obtained from CD spectral deconvolution.

**S.Figure 8.** Putative heparin binding sites localised in the PACAP and glucagon families.

**S.Figure 9.** Characterization of the solvated states of PACAP-family peptides by molecular dynamics simulations.

**S.Figure 10.** Steered molecular dynamics (SMD) simulation of peptide-HEP complexes.

**S.Figure 11** Molecular dynamics (MD) simulation of the self-assembly of three PACAP-27 peptides in the absence of heparin

**S.Figure 12.** Prediction of aggregation-prone regions along the sequences of the PACAP–glucagon superfamily.

**S.Figure 13.** Monitoring the aggregation propensity of the non-terminally protected APR hexapeptide cores by CD spectroscopy.

**S.Figure 14.** IR spectra of the terminally protected and non-protected APR-s after 3-day long incubation.

**S.Figure 15.** Molecular packing of two adjacent β-sheet planes, each containing four β-strands stacked along the fibril axis.

**S.Figure 16.** Ramachandran diagram of the amyloid-like crystal structures of APRs in the PACAP family.

**S.Figure 17.** Pairwise amino acid identity heatmap of the PACAP–glucagon peptide family and hierarchical clustering dendrogram based on amino acid sequence identity (first 27 residues).

**S.Figure 18.** Pairwise nucleotide identity heatmap of the PACAP–glucagon gene family and hierarchical clustering dendrogram based on nucleotide sequence identity (first 81 bp).

**S.Figure 19.** Position-specific amino acid conservation and codon usage diversity across peptide families of the PACAP–glucagon superfamily.

**S.Figure 20.** Position-specific conservation and aggregation propensity across species in the PACAP family.

**S.Figure 21.** Amyloid aggregation analysis of PACAP-related APR hexapeptides from *Chelyosoma productum*.

**S.Figure 22.** Monitoring of the amyloid formation for tunicate derived PRP-like 1 peptide in presence and absence of heparin.

**S.Figure 23.** Monitoring of the amyloid formation for tunicate derived PACAP 1 peptide in presence and absence of heparin.

**Supplementary Table 1.** Members of the PACAP peptide family across vertebrate species.

**Supplementary Table 2.** The 27-residue bioactive segments and their corresponding coding sequences from the PACAP peptide family were used to sequence and genomic analyses in this work.

**Supplementary Table 3.** Nearest-neighbour sequence similarity analysis of PACAP family peptides at the nucleotide and amino acid levels.

**Supplementary Table 4/1.** Position-specific mutational pathways in PACAP sequences.

**Supplementary Table 4/2.** Position-specific mutational pathways in VIP sequences.

**Supplementary Table 4/3.** Position-specific mutational pathways in PRP sequences.

**Supplementary Table 4/4.** Position-specific mutational pathways in PH sequences.

**Supplementary Table 4/5.** Position-specific mutational pathways in GHRH sequences.

**Supplementary Table 4/6.** Position-specific mutational pathways in SCT sequences.

**Supplementary Table 4/7.** Position-specific mutational pathways in GCG sequences.

**Supplementary Table 4/8.** Position-specific mutational pathways in GLP-1 sequences.

**Supplementary Table 4/9.** Position-specific mutational pathways in GLP-2 sequences.

**Supplementary Table 5.** X-ray data collection and refinement statistics.

**General description of Supplementary Figure 1-6: (A)** CD spectra of the initial peptide states as a function of pH. **(B)** Thioflavin-T fluorescence kinetics monitored over 72 h at 37 °C with orbital shaking in 96-well plates in the presence of heparin. **(C)** CD spectra of the corresponding incubated end states. **(D)** Thioflavin-T kinetics recorded in the absence of heparin. **(E)** CD spectra of the corresponding end states without heparin. **(F)** Relative abundance of the charged microspecies (right y-axis) and overall charge (left x-axis) along the pH. The graph highlights how the relative abundance of differently charged microspecies corresponds to the observed presence or absence of amyloid self-assembly at a given pH. **(G)** Representative AFM micrographs of incubated samples, indicating pH and the presence of heparin. Peptide specific features and observations are detailed individually for PACAP (**SFig. 1**), VIP (**SFig. 2**), PRP (**SFig. 3**), PHI (**SFig. 4**), GHRH (**SFig. 5**) and SCT (**SFig. 6**).

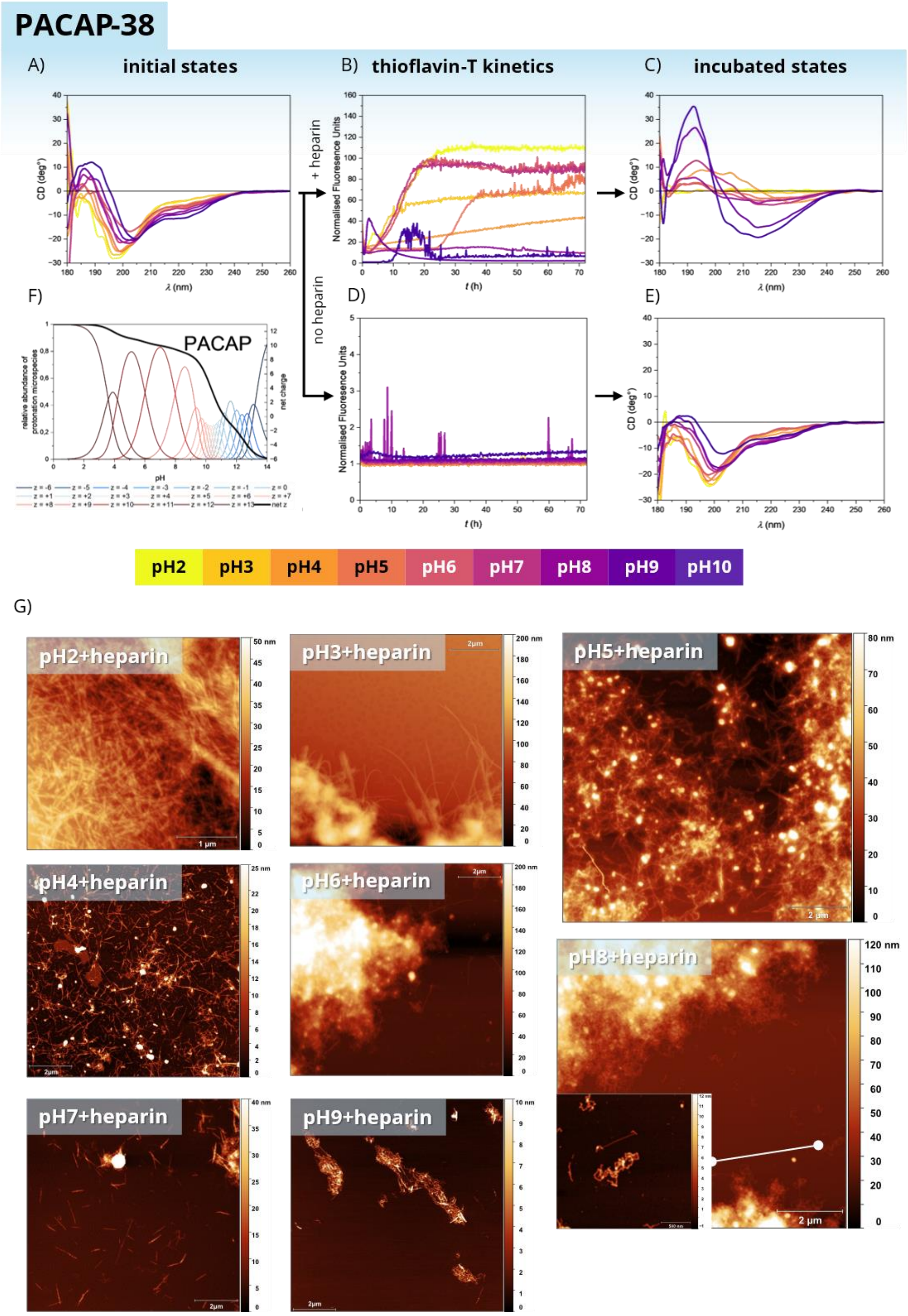


**Supplementary Figure 1. Monitoring of the amyloid formation for PACAP-38 in presence and absence of heparin.** For figure legend see general description at page 2. In the presence of heparin at pH > 8, ThT kinetics did not unambiguously indicate amyloid formation based on fluorescence changes; however, CD spectroscopy revealed a pronounced β-structure in the incubated samples within the same pH range, which was further confirmed by AFM. PACAP-38 is exceptionally rich in basic amino acids and therefore exists exclusively as positively charged microspecies across the examined pH range. According to the screening results, amyloid formation occurs only in the presence of heparin, whereas no self-assembly is observed in its absence. This suggests that negatively charged heparin is required to neutralize the excess positive charge of PACAP-38, thereby enabling the initiation of self-association.

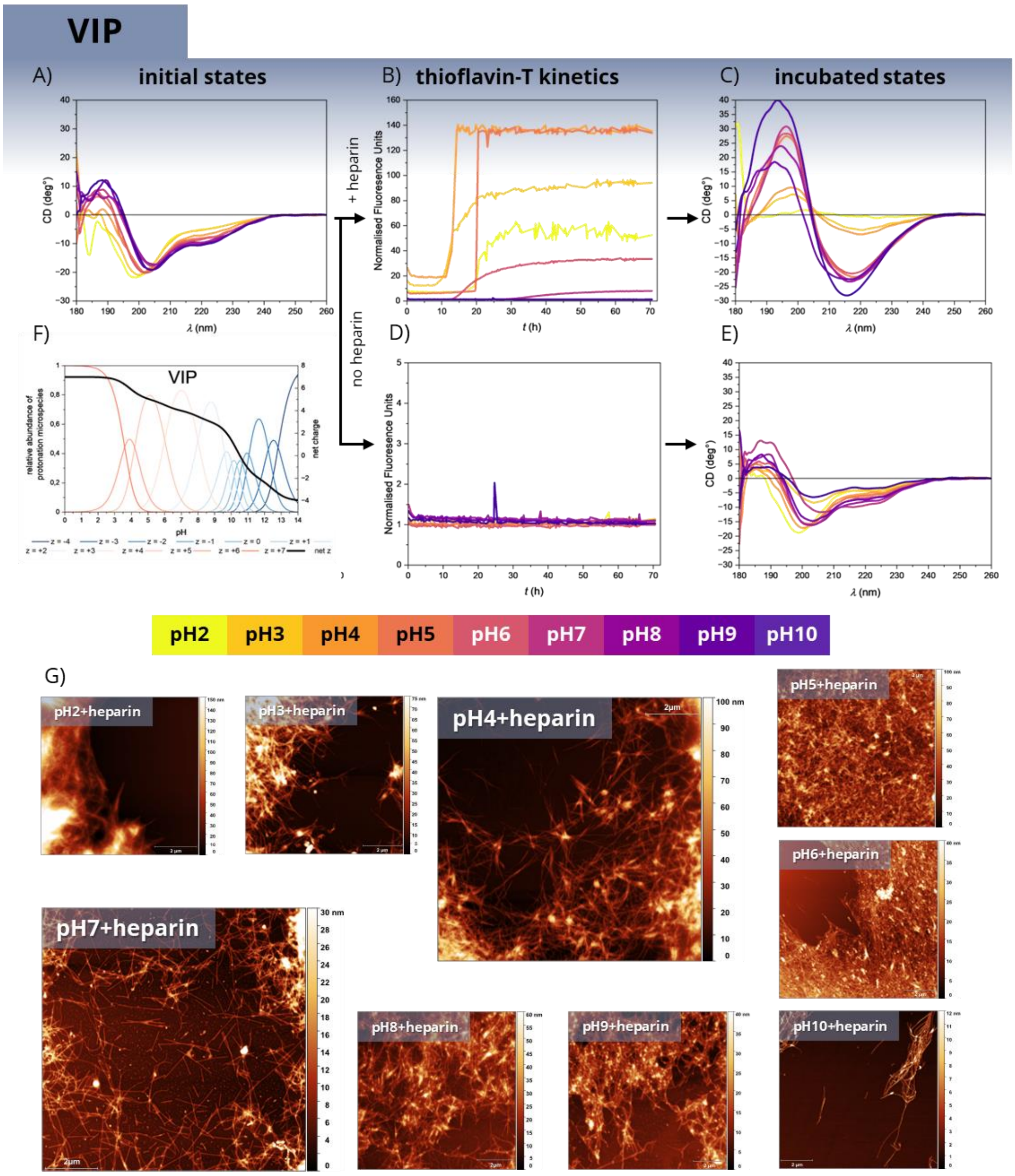


**Supplementary Figure 2. Monitoring of the amyloid formation for VIP in presence and absence of heparin.** For figure legend see general description at page 2. VIP formed amyloid across the entire examined pH range (+1 to +7 net charge) in the presence of heparin. Although ThT fluorescence did not show a pronounced increase under alkaline conditions (pH > 8), CD spectra exhibited strong β-sheet signatures, and AFM screening confirmed the presence of fibrillar structures.

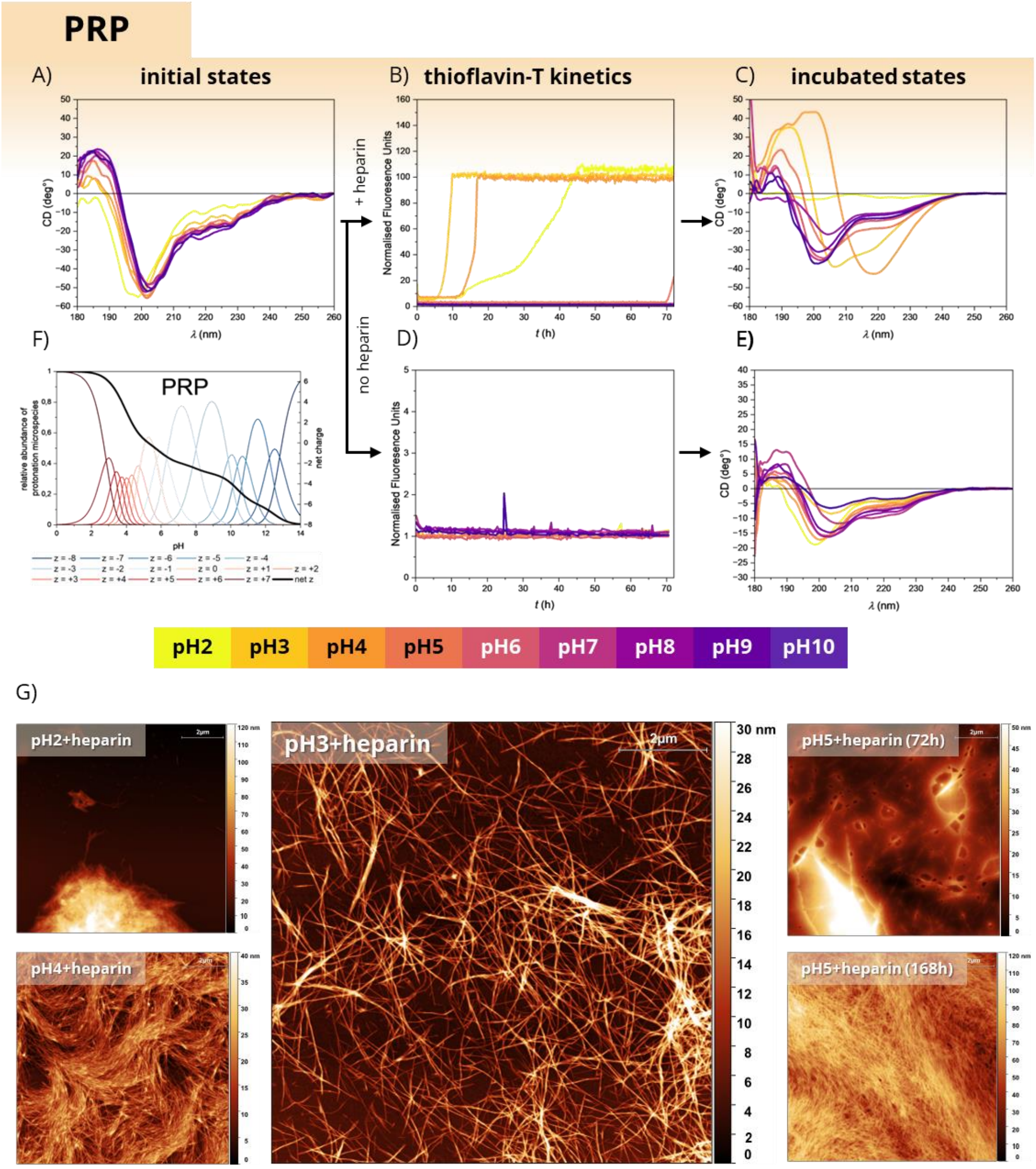


**Supplementary Figure 3. Monitoring of the amyloid formation for PRP in presence and absence of heparin**. For figure legend see general description at page 2. For PRP, CD and ThT measurements consistently indicated amyloid formation within the same pH range (pH 2–5). The apparent plateau values in the ThT fluorescence kinetics lie beyond the upper dynamic range of the detector. After 72 hours of incubation, the sample at pH 5 exhibited a CD spectrum characteristic of pronounced β-structure; however, ThT fluorescence and AFM suggested only partial fibril formation at this time point, which became clearly detectable after one week of incubation. Within this aggregation-prone pH range, the predominant microspecies carry individual charges between +2 and +7, consistent with a model in which Coulombic peptide–heparin interactions facilitate subsequent peptide self-association. In the absence of heparin, no significant structural changes were observed for PRP over 72 hours.

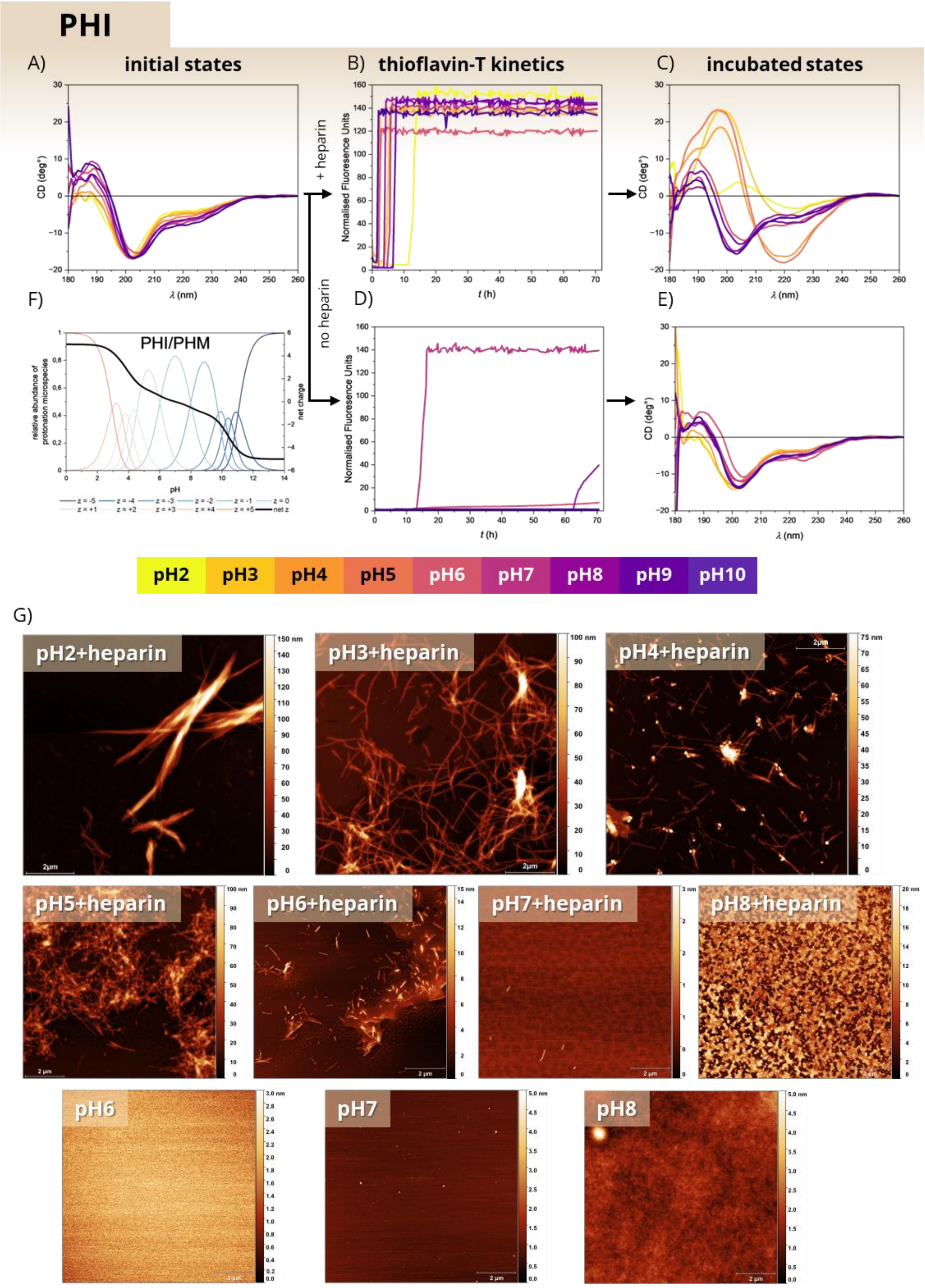


**Supplementary Figure 4. Monitoring of the amyloid formation for PHI in presence and absence of heparin.** For figure legend see general description at page 2. Amyloid formation for PHI was observed within the pH range of 2–5, where the predominant microspecies carry individual charges between +5 and +1. ThT kinetics showed a rapid and pronounced increase in fluorescence; however, CD and AFM analyses confirmed that at pH > 7 in the presence of heparin, the ThT signal represents a false-positive result. Similarly, in the absence of heparin at pH 7, the increased ThT fluorescence was not accompanied by structural changes, suggesting an artefactual signal. The apparent plateau values in the ThT fluorescence kinetics lie beyond the upper dynamic range of the detector.

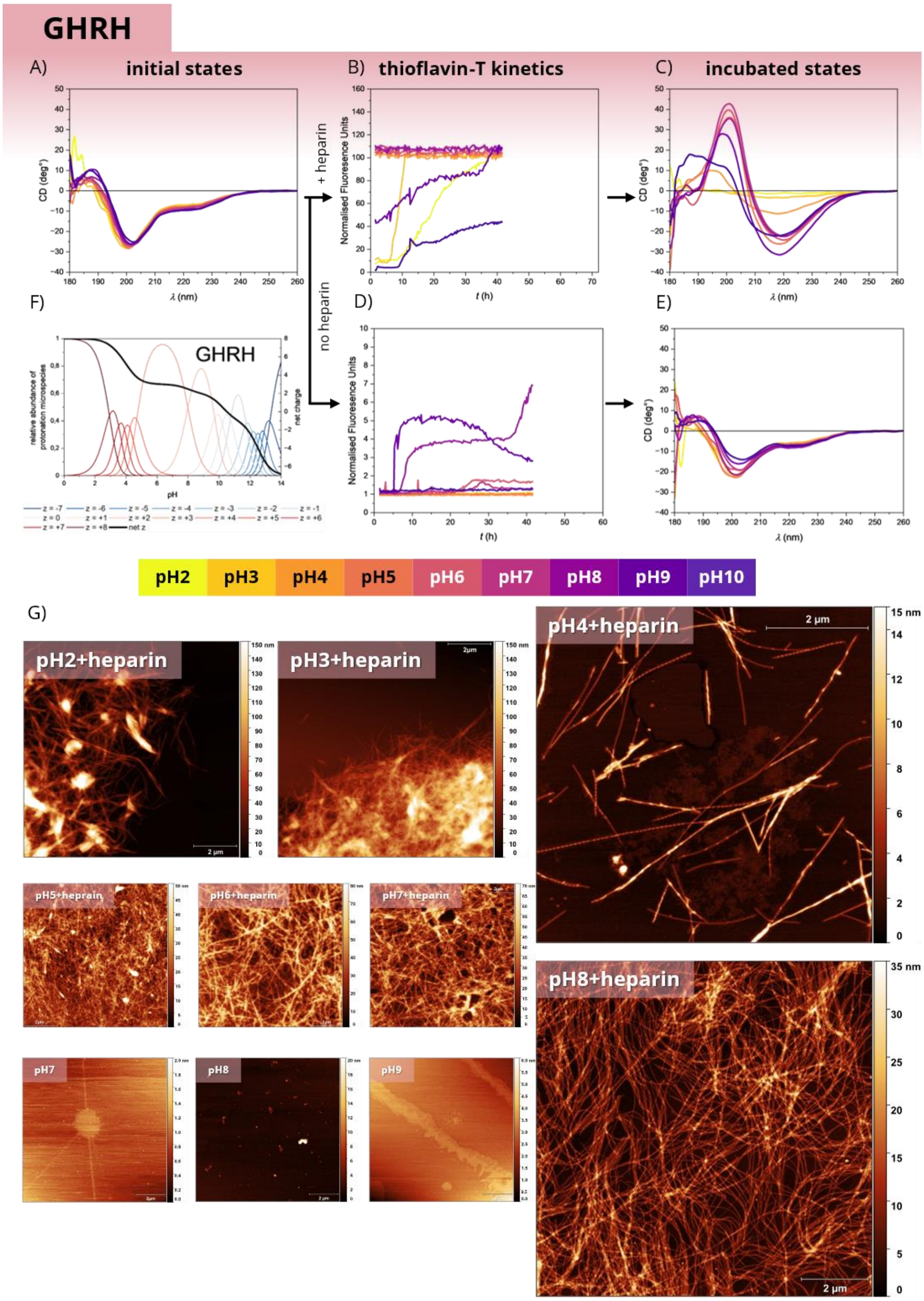


**Supplementary Figure 5. Monitoring of the amyloid formation for GHRH in presence and absence of heparin.** For figure legend see general description at page 2. GHRH formed amyloid convincingly across the pH range 2–8, as confirmed by all applied techniques. Within this interval, the predominant microspecies carry individual charges between +8 and +3. At pH > 8, both CD and ThT measurements suggested amyloid formation; however, AFM screening did not reveal fibrillar structures. In heparin-containing samples, the apparent plateau values in the ThT kinetics exceeded the upper dynamic range of the detector. In the absence of heparin, ThT produced false-positive signals in the pH 7–9 range, as neither CD nor AFM confirmed the presence of β-sheet–rich amyloid fibrils. During ThT measurements, the detector malfunctioned after 40 hours and failed to record further signals; nevertheless, the incubation program (temperature control and orbital shaking) continued uninterrupted.

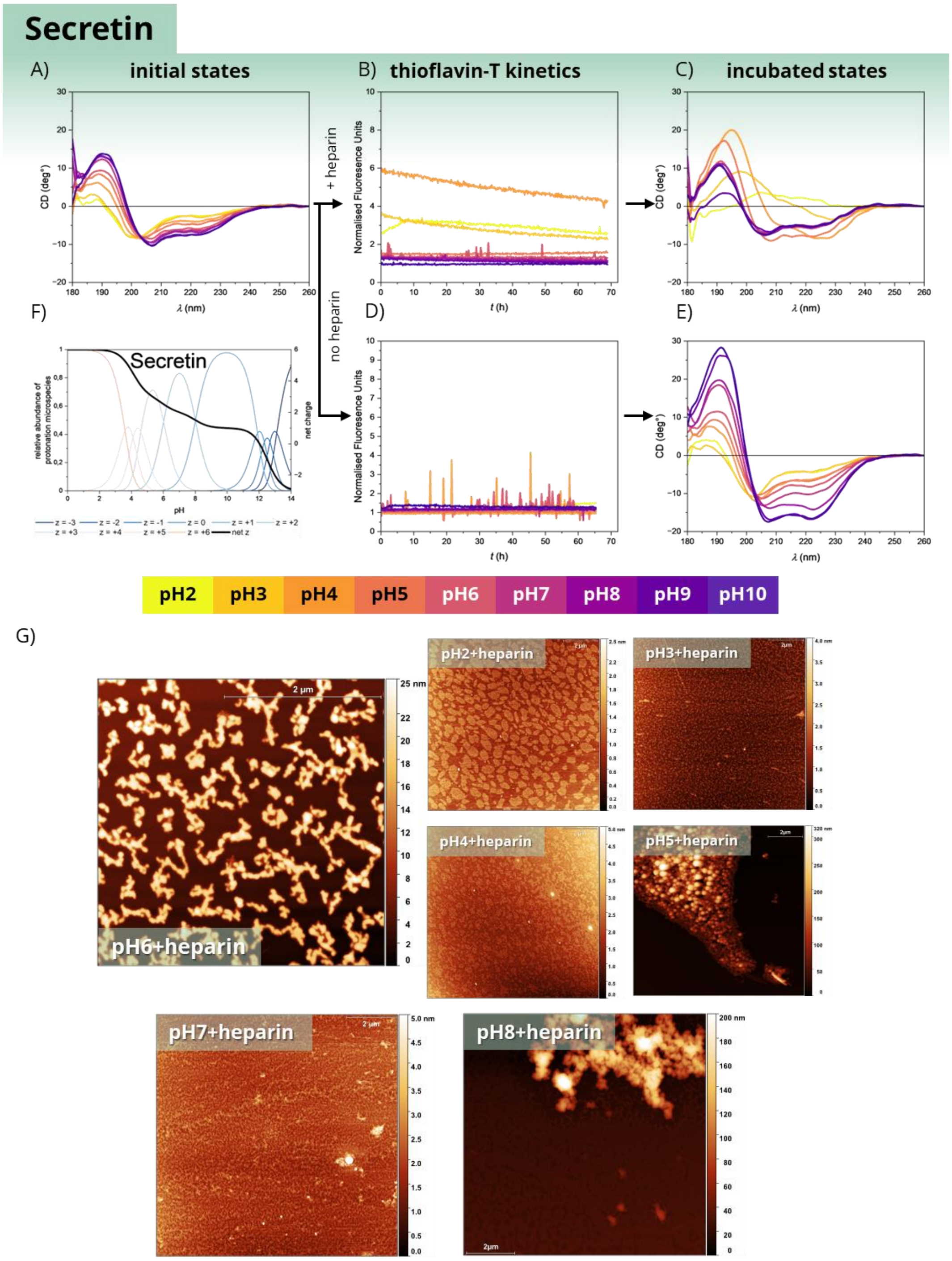


**Supplementary Figure 6. Monitoring of the amyloid formation for SCT in presence and absence of heparin**. For figure legend see general description at page 2. Secretin did not exhibit unambiguous amyloid aggregation under any of the tested incubation conditions within 72 hours. In contrast to the other five peptides, its initial CD spectra revealed a pronounced pH dependence of secondary structure: predominantly disordered under acidic conditions and increasingly α-helical at alkaline pH. Upon incubation in the absence of heparin, this pH-dependent structural behaviour became even more distinct. In the presence of heparin, the elevated but decaying ThT fluorescence observed in the pH 2–4 range reached only a fraction of the intensity measured for the amyloid-forming states of the other peptides. Although CD indicated β-like structural rearrangement in this acidic range, AFM micrographs did not reveal the characteristic fibrillar morphology observed for the other five peptides. At pH 6, the detected aggregates displayed a relatively uniform height profile, inconsistent with simple protein precipitation; however, a β-sheet–rich amyloid nature could not be conclusively confirmed based on the available data.

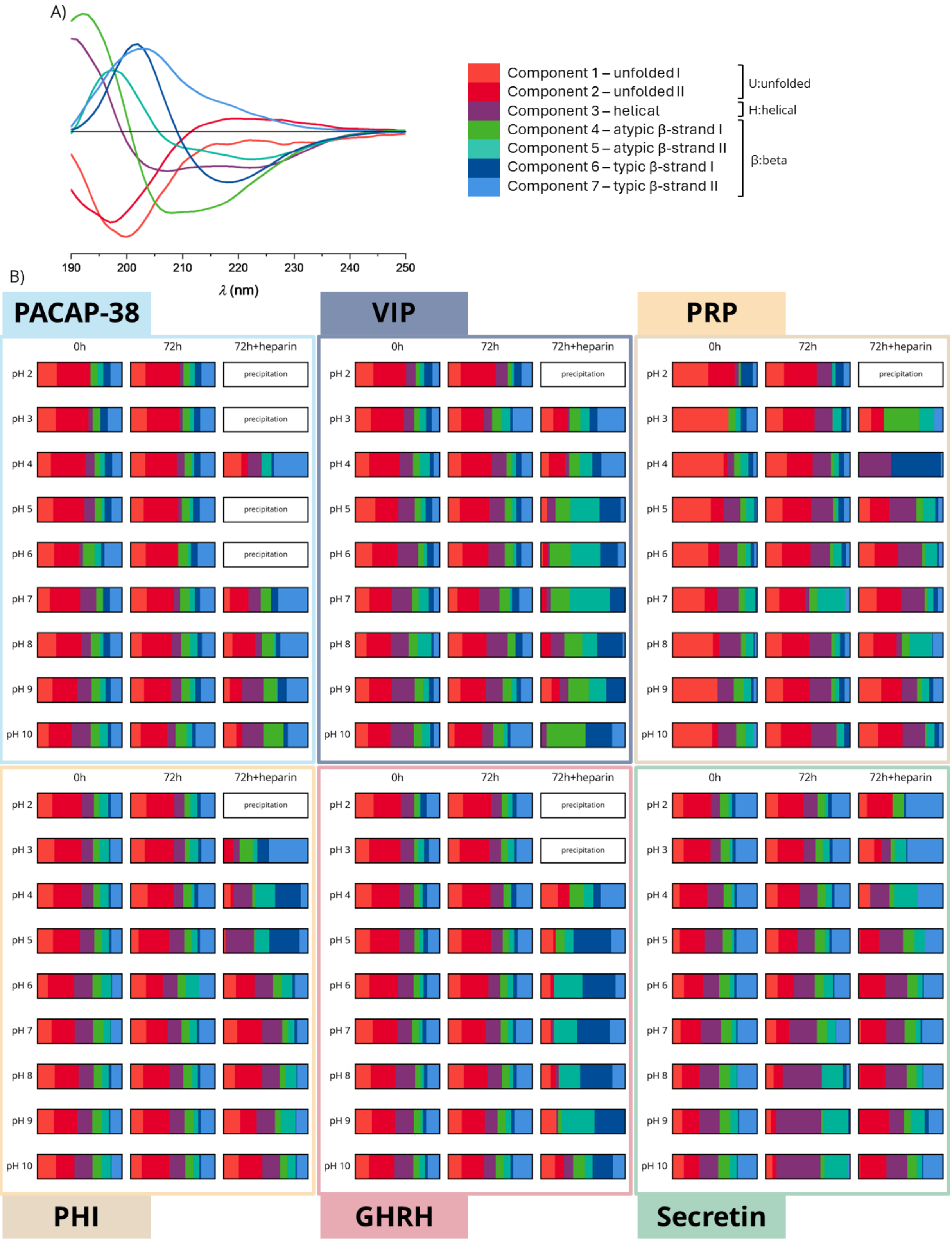


**Supplementary Figure 7. Color-coded bar chart analysis of the secondary-structure components obtained from CD spectral deconvolution. (A)** A total of 153 CD spectra were analysed by deconvolution into seven components. The seven-component model provided the most optimal set of basis curves, including at least one representative curve characteristic of disordered, α-helical, and β-sheet secondary structures. During deconvolution, CD spectra were normalized only to amino acid residue number, as the initial solutions were prepared at identical molar concentrations. Because peptide self-association during incubation alters the soluble peptide concentration—thereby affecting CD intensity and potentially the accuracy of deconvolution—samples showing substantial aggregation (reflected by a marked decrease in CD intensity) were excluded from the deconvolution matrix. No significant difference was observed between the two unfolded basis curves (shown in red shades). Unfolded I exhibits a minimum at 200 nm, whereas Unfolded II shows a slightly hypsochromic shifted minimum and is overall hyperchromic shifted relative to Unfolded I. The characteristic helical component (purple), displaying a maximum at 190 nm and minima at 205 and 225 nm, is present only in low proportion in the non-incubated samples despite the intrinsic helix propensity of the peptides. Its most pronounced contribution is observed for secretin at pH > 8 following incubation in the absence of heparin. In amyloid aggregation, the resulting β-sheet architectures can adopt a range of CD spectral signatures, often deviating from the canonical β-sheet spectrum, thereby complicating data interpretation[1]. In the present deconvolution, four β-like basis spectra were distinguished: two typical and two atypical components. Typical β-strand I corresponds to a CD profile characteristic of parallel β-sheets and is most prominently represented in PRP at pH 4 and in the amyloid state of GHRH. Typical β-strand II reflects an antiparallel β-sheet CD signature and is most evident in PACAP, secretin, and in VIP/PHI incubated with heparin at pH < 4. Atypical β-sheet I displays a maximum around 195 nm reminiscent of α-helical structure but exhibits only a single pronounced minimum near ~205 nm. This component is most abundant in heparin induced samples of VIP across the pH range and in PRP samples at pH 3.Atypical β-sheet II shows a hypochromic shifted maximum relative to the canonical α-helical spectrum. **(B)** Relative proportions (%) of the seven deconvoluted components as a function of pH and incubation conditions. Interpretation of component ratios in a single sample can be misleading; therefore, it is more informative to examine changes in component distribution as a function of pH or incubation. In the initial samples, disordered structures account for approximately 40–50%, the helical component for 10–20%, and the remaining fraction consists of various β-like components. In the absence of heparin (middle columns), no substantial changes in component distribution were observed after 72 hours of incubation, indicating that the peptides largely retained their disordered or partially helical conformations. In contrast, incubation in the presence of heparin led to a pronounced increase in β-like components (blue–green shades) at specific pH values. At pH conditions where no self-assembly occurred, the bar chart distribution remained comparable to that of the initial state.

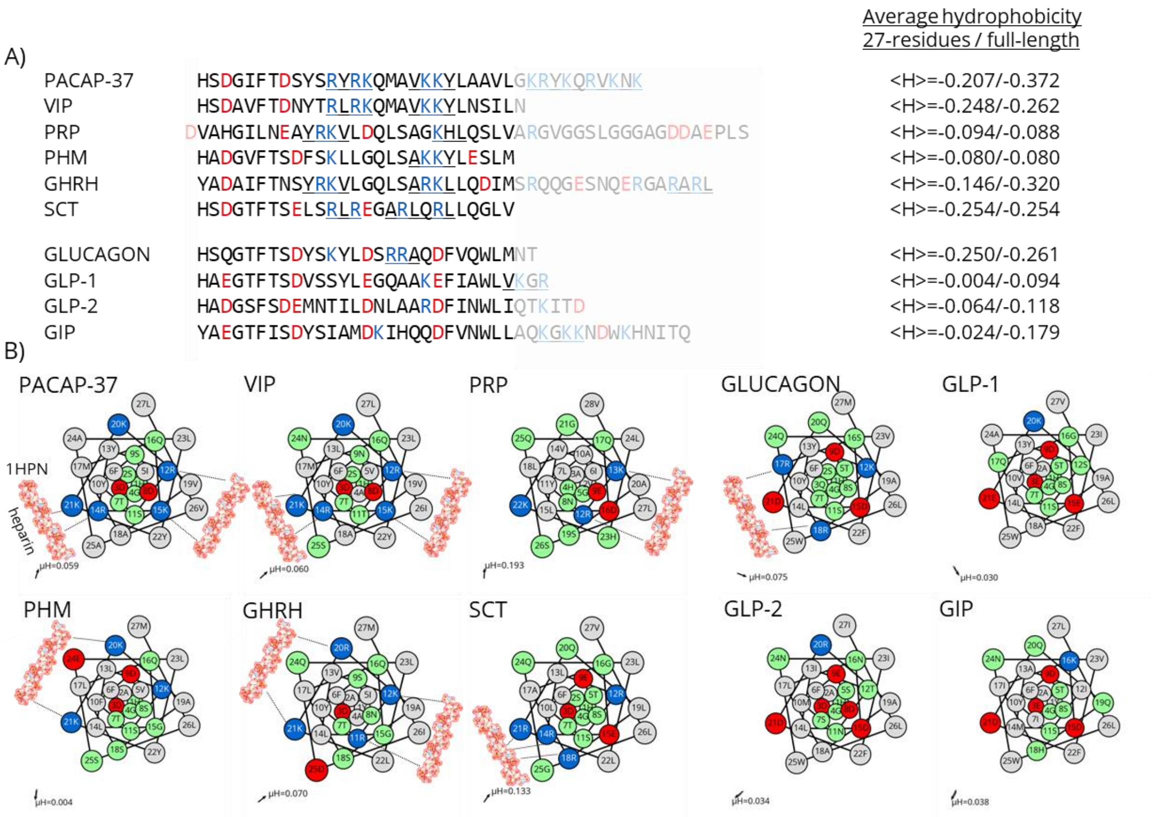


**Supplementary Figure 8. Putative heparin binding sites localised in the PACAP and glucagon families. (A)** The bioactive 1–27 segment of the PACAP and glucagon family peptides is shown in full intensity, whereas the C-terminal extensions are faded. Within the sequences, acidic residues at neutral pH are highlighted in red and basic residues in blue. Regions with potential high heparin affinity are underlined. The residue-normalized average Eisenberg hydrophobicity (higher values indicate greater hydrophobicity) is indicated next to each sequence. Except for PRP, the polar C-terminal extensions decrease the overall average hydrophobicity and, in the case of PACAP-37, GHRH, GLP-1, and GIP, introduce additional putative heparin-binding motifs. **(B)** Helix-wheel representations depict the hypothetical helical arrangement of the 27-residue bioactive cores. Owing to the intrinsic helical propensity of these peptides, it is informative to visualize the putative heparin-binding sites on the helical surface, even though CD data indicate only partial helicity in solution. Lys/Arg residues that are distant in the primary sequence may become spatially proximal upon helix formation, particularly across successive helical turns, thereby forming potential heparin-binding patches. In the helix-wheel diagrams, a heparin molecule (PDB ID: 1HPN) is shown to indicate the possible binding interfaces. The Eisenberg hydrophobic moment vector[2], representing the magnitude and direction of helix amphipathicity, which captures both the magnitude and spatial orientation of hydrophobic segregation along the helical surface, is indicated in the lower-left corner of each helix. The hydrophobic moment vector of an amphipathic helix is informative because it defines the spatial segregation of hydrophobic and charged surfaces. Such amphipathicity is essential for effective GPCR–ligand recognition[3], while also highlighting potential electrostatically driven protein–GAG interaction sites and coiled-coil association interfaces that may serve as stabilized intermediates toward higher-order aggregation[4]. Examination of the peptide helix wheels and their hydrophobic moment vectors shows that the vectors point opposite to the proposed polar heparin-binding surface, thereby defining a complementary hydrophobic face that may promote hydrophobic–hydrophobic interactions.

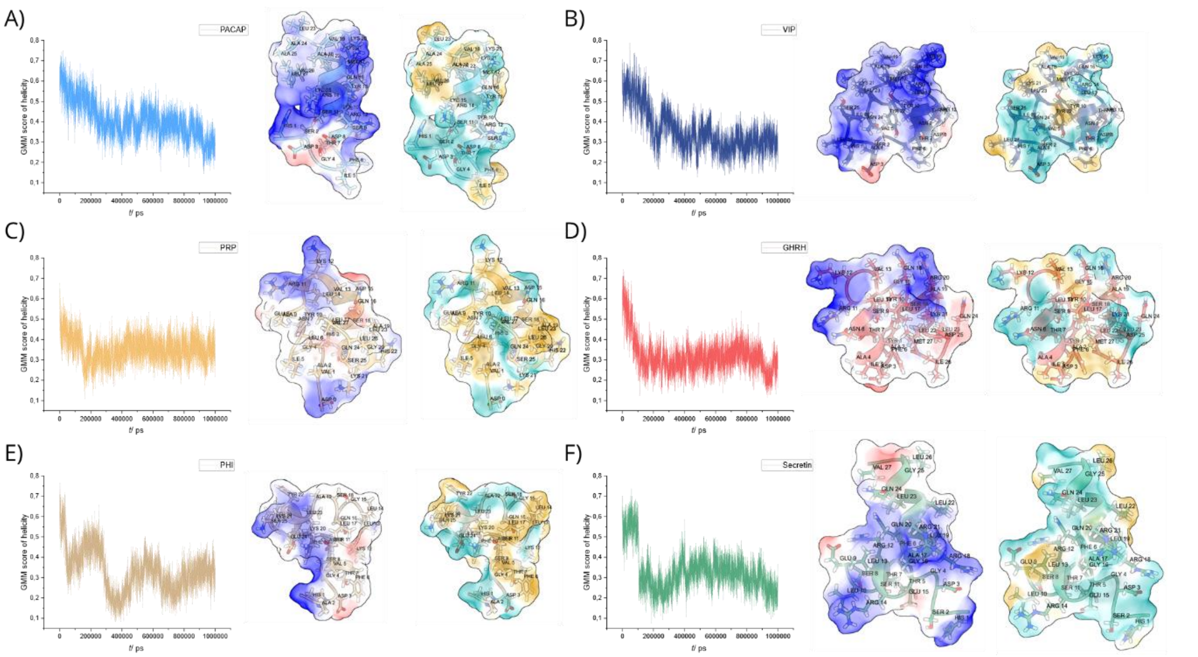


G)

| Peptide | total charge at pH2 | | total charge at pH7 | |
|---|---|---|---|---|
| | Residue 1-27 | Full length | Residue 1-27 | Full length |
| PACAP | 7 | 13 | 4 | 10 |
| VIP | 7 | 7 | 4 | 4 |
| PRP* | 6 | 7 | 1 | -2 |
| PHI | 5 | 5 | 1 | 0 |
| Secretin | 6 | 6 | 2 | 2 |
| GHRH | 4 | 8 | 2 | 3 |

H)

| Peptide | Number of clusters | Top 1 cluster population | H-bond | VdW | Ionic | π-π stack | π-cation | GFN2-xTB dipole moment (D) |
|---|---|---|---|---|---|---|---|---|
| PACAP | 11 | 41% | 22.8 | 10.1 | 0.4 | 1.1 | 0.0 | 45.6 |
| VIP | 5 | 94% | 20.1 | 13.8 | 0.4 | 0.2 | 0.1 | 117.6 |
| PRP | 1 | 100% | 21.0 | 13.7 | 0.6 | 0.0 | 0.0 | 27.3 |
| PHI | 5 | 85% | 22.2 | 12.2 | 0.8 | 0.2 | 0.0 | 29.4 |
| Secretin | 16 | 25 % | 21.5 | 9.7 | 0.5 | 0.0 | 0.1 | 84.7 |
| GHRH | 9 | 70 % | 18.9 | 15.0 | 0.5 | 0.3 | 0.0 | 139.6 |

**Supplementary Figure 9. Characterization of the solvated states of PACAP-family peptides by molecular dynamics simulations. (A)-(F)** Helicity (min: 0, max: 1) as a function of simulation time, quantified by the Gaussian Mahalanobis Mean score (GMMs), calculated from the backbone torsion angles in each frame. Representative structures correspond to the most populated cluster centres from the last 200 ns of each simulation and depict surface charge distribution and hydrophobicity (brown: more hydrophobic; turquoise: more hydrophilic). (G) Approximate overall charge comparison calculated at pH 2 (Asp/Glu fully protonated) and pH 7 (Asp/Glu fully deprotonated) for the full-length peptides used in the screening experiments and for the 27-residue, C-terminally protonated models used in the MD simulations. PRP contains an additional Asp at the N-terminus, designated as position 0 to maintain consistent numbering. (H) Stabilizing intramolecular interaction counts averaged over the last 200 ns of the simulation and the dipole moments of the most representative (Top 1) cluster centres calculated from the same time interval.

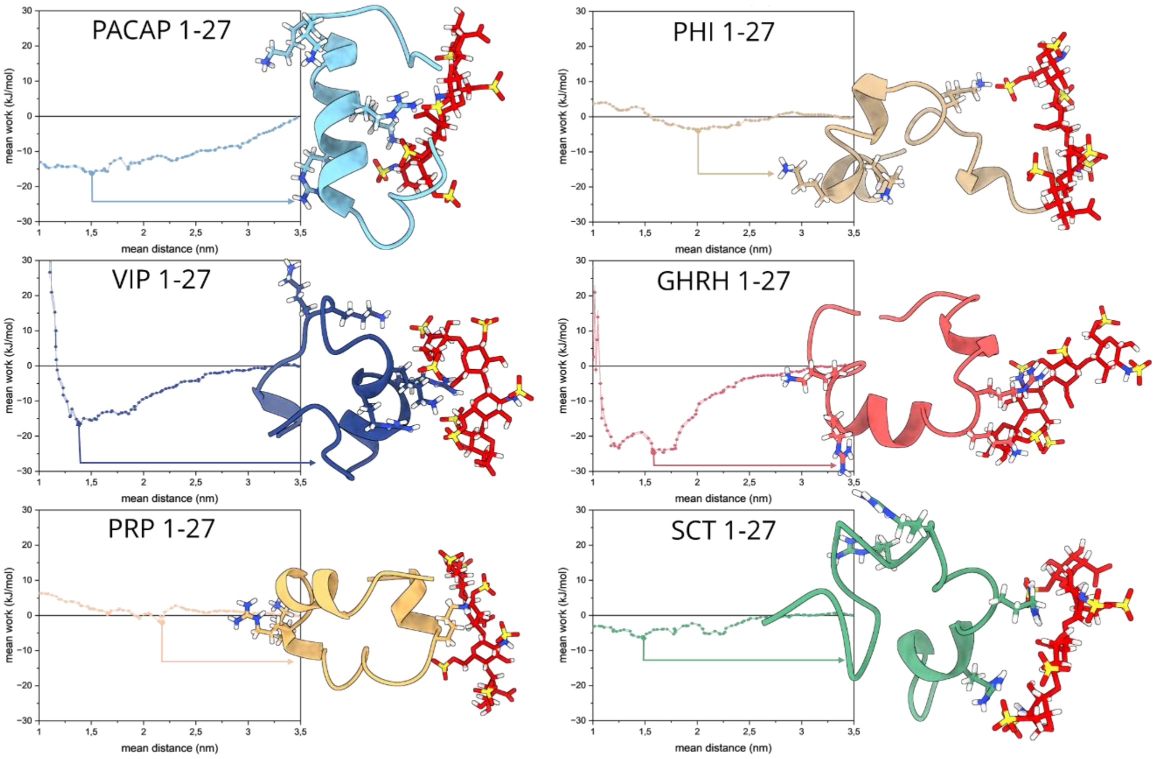


**Supplementary Figure 10. Steered molecular dynamics (SMD) simulation of peptide-HEP complexes.** Steered molecular dynamics (SMD) simulations were performed by applying a time-dependent harmonic bias to the distance (d) between the geometric centres of the peptide and heparin molecules. This approach gradually drives the two molecules together while recording the work along the trajectory. Representative structures at the minima of the work curves are shown.

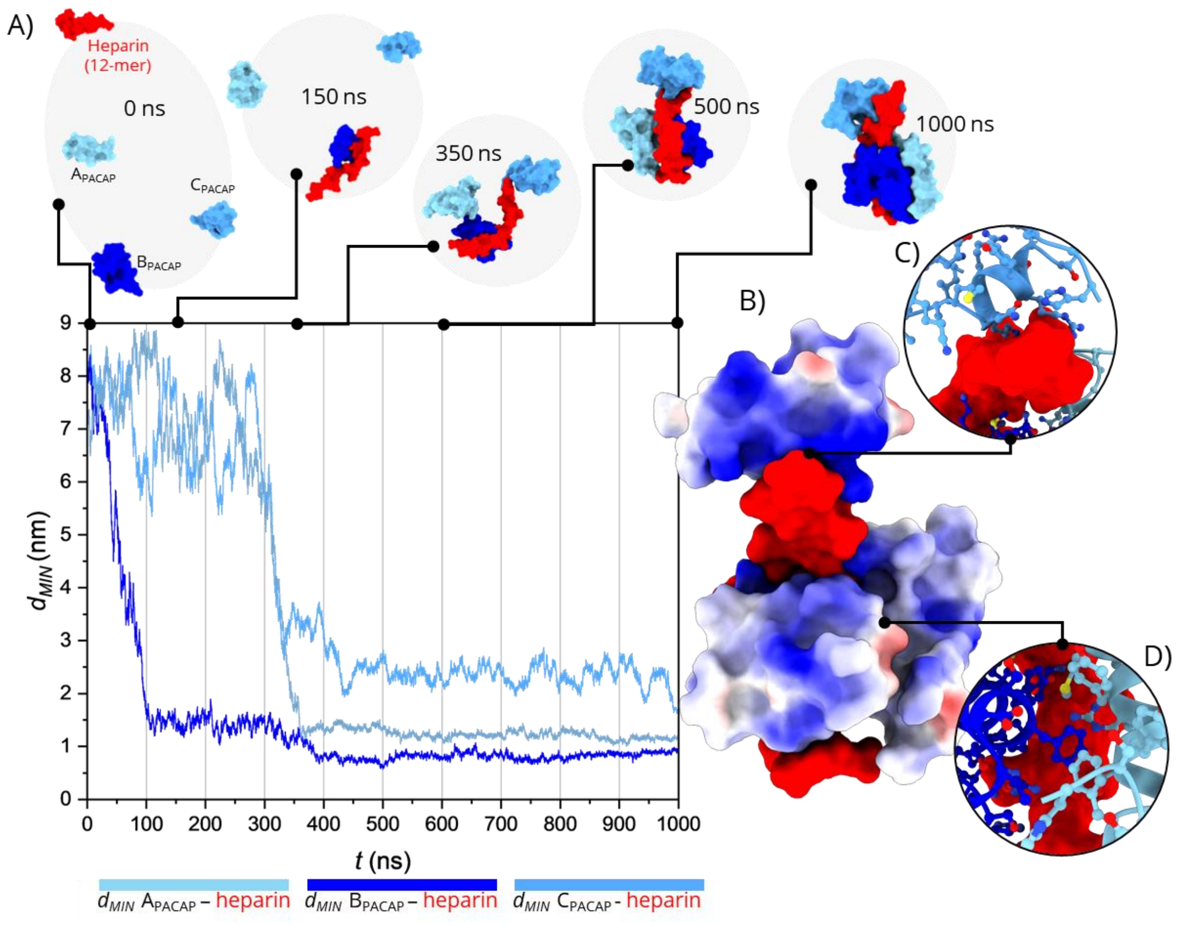


**Supplementary Figure 11. Molecular dynamics (MD) simulation of the self-assembly of three PACAP-27 peptides in the presence of a 12-mer heparin fragment (representative of low-molecular-weight heparin)** **(A)** Time evolution of the minimum distances ($d_{MIN}$) between peptide–heparin pairs, with representative snapshots of the corresponding molecular assemblies. Negatively charged GAG chains recruit positively charged peptides from solution, forming stable electrostatic interactions and increasing their local concentration. **(B)** Electrostatic surface representation of the peptide (blue: positively charged, white: neutral, red: negatively charged) in complex with heparin (red) at 500 ns. **(C)** Lys/Arg-rich gatekeeper regions of PACAP-27 anchor to the negatively charged GAG surface. **(D)** Hydrophobic surfaces of the captured peptides cluster together, facilitating subsequent amyloid self-assembly.

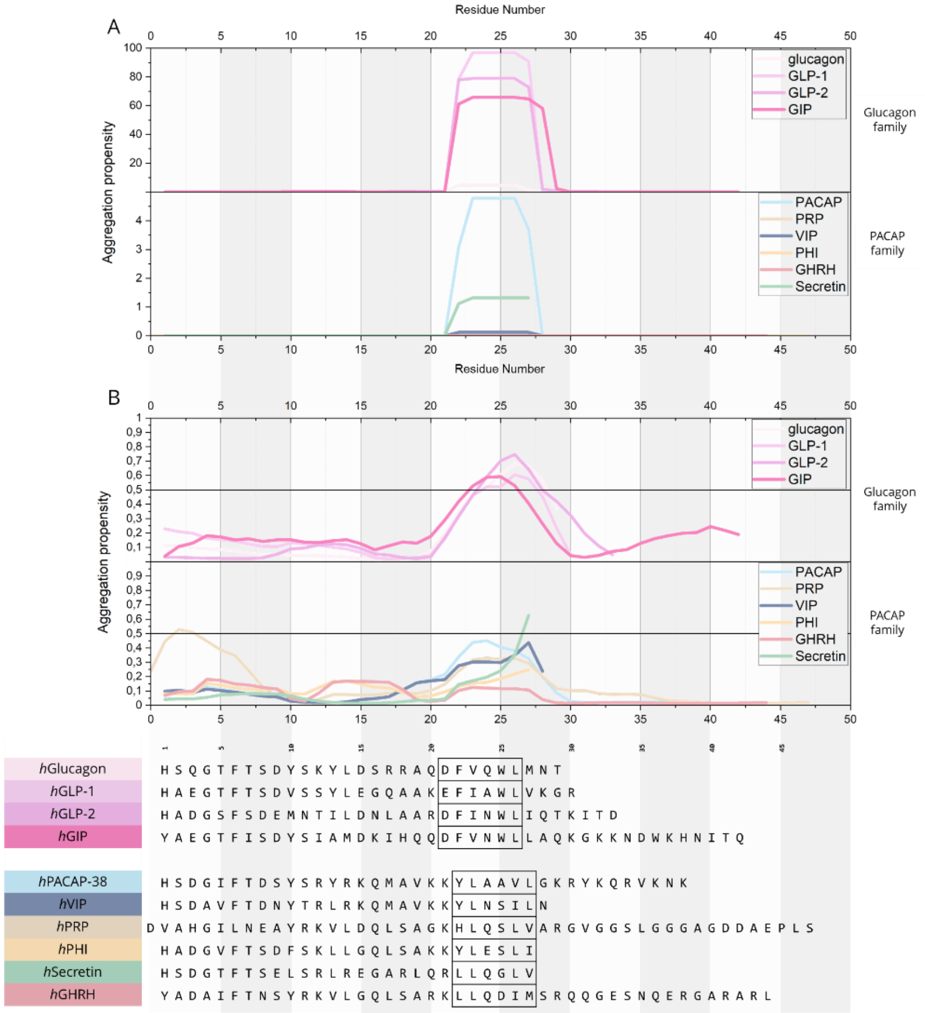


**Supplementary Figure 12. Prediction of aggregation-prone regions along the sequences of the PACAP–glucagon superfamily. (A)** Aggregation propensities predicted by TANGO and **(B)** by AggreProt for grouped PACAP and glucagon family sequences. Boxed regions indicate the APR segments identified in the glucagon and PACAP families. In the glucagon family, position 21 contains a pH-sensitive Asp or Glu that acts as a negatively charged gatekeeper residue. In contrast, the PACAP family features a conserved Lys or Arg at this position, which remain positively charged across physiological pH values. To reduce potential repulsive forces during self-assembly arising from the disproportionately high positive charge of these Arg/Lys residues in the PACAP-related hexapeptides, the APR hexapeptide segment was shifted by one residue relative to the region analysed in the glucagon family.

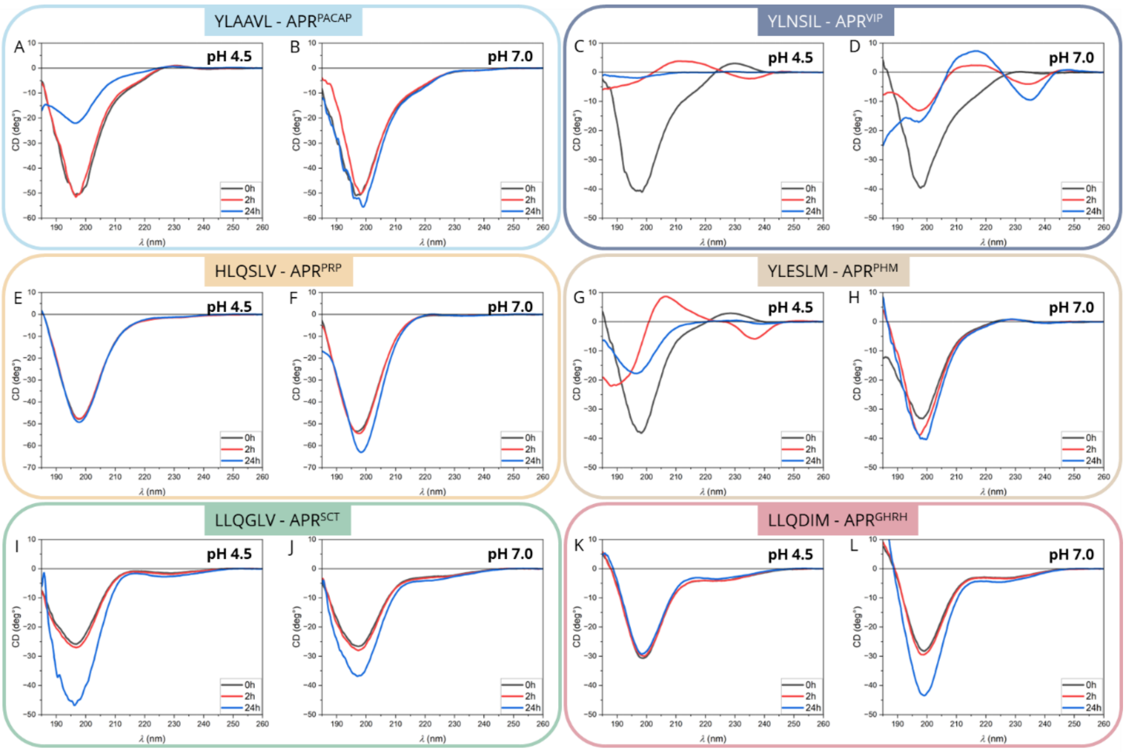


**Supplementary Figure 13. Monitoring the aggregation propensity of the non-terminally protected APR hexapeptide cores by CD spectroscopy.** The initial CD spectra in all cases displayed predominantly disordered conformations characteristic of short hexapeptide sequences (black traces). Upon incubation, only APR$^{VIP}$ (C, D) and APR$^{PHM}$ (G) exhibited marked β-like CD signatures, consistent with β-structure formation but not definitively confirming canonical β-sheet organization.

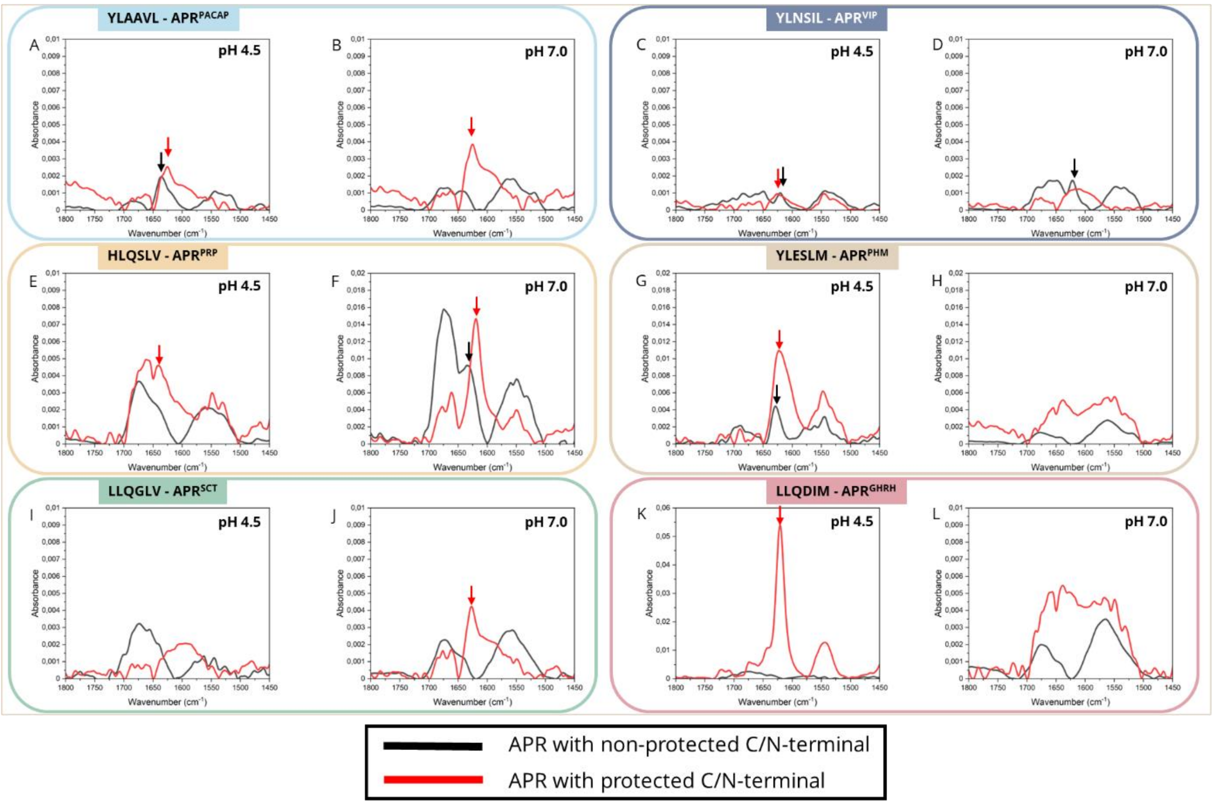


**Supplementary Figure 14. IR spectra of the terminally protected and non-protected APR-s after 3-day long incubation.**

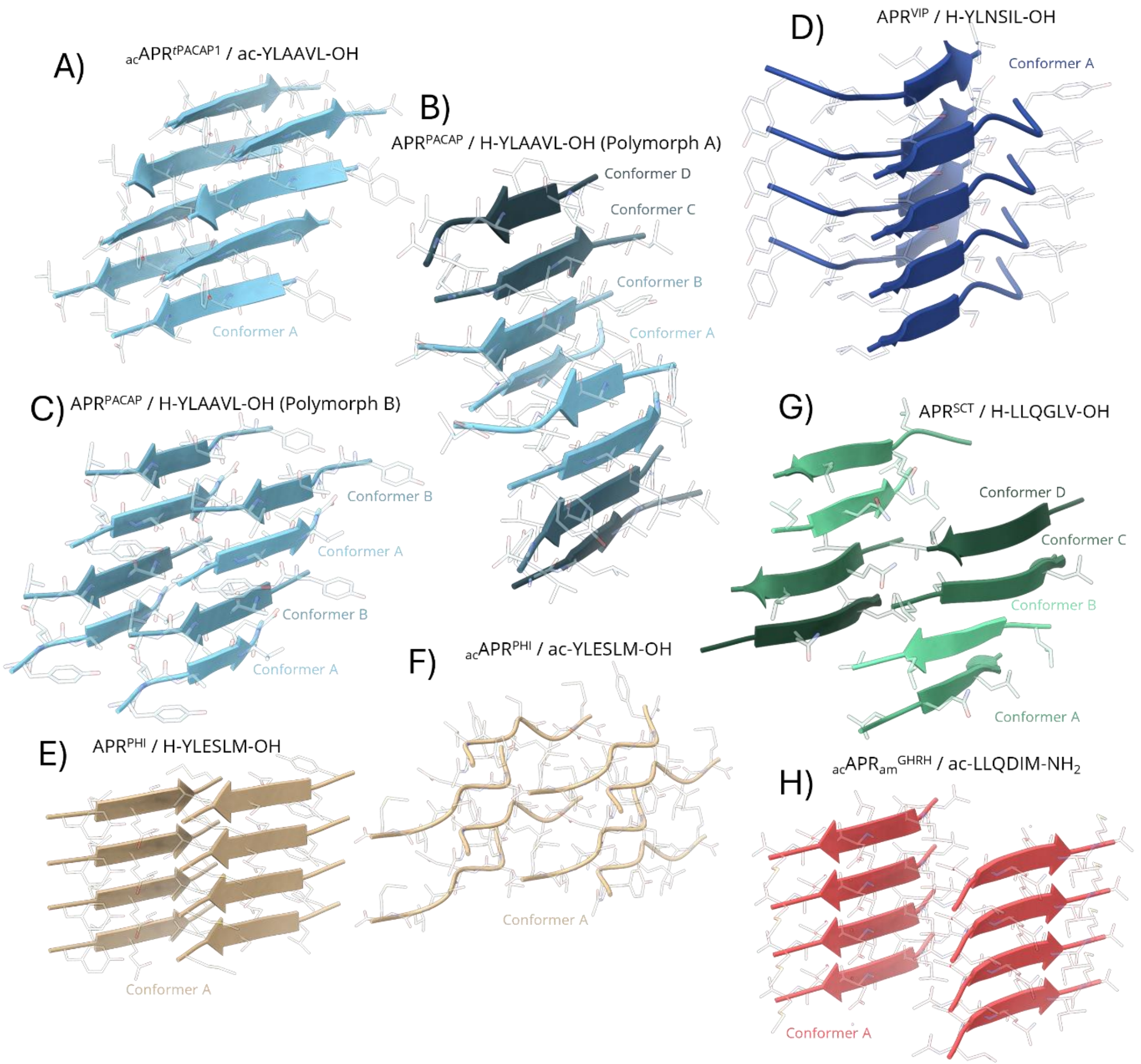


**Supplementary Figure 15. Molecular packing of two adjacent β-sheet planes, each containing four β-strands stacked along the fibril axis.** Identical conformers (labelled A–D) are consistently color-coded within each model.

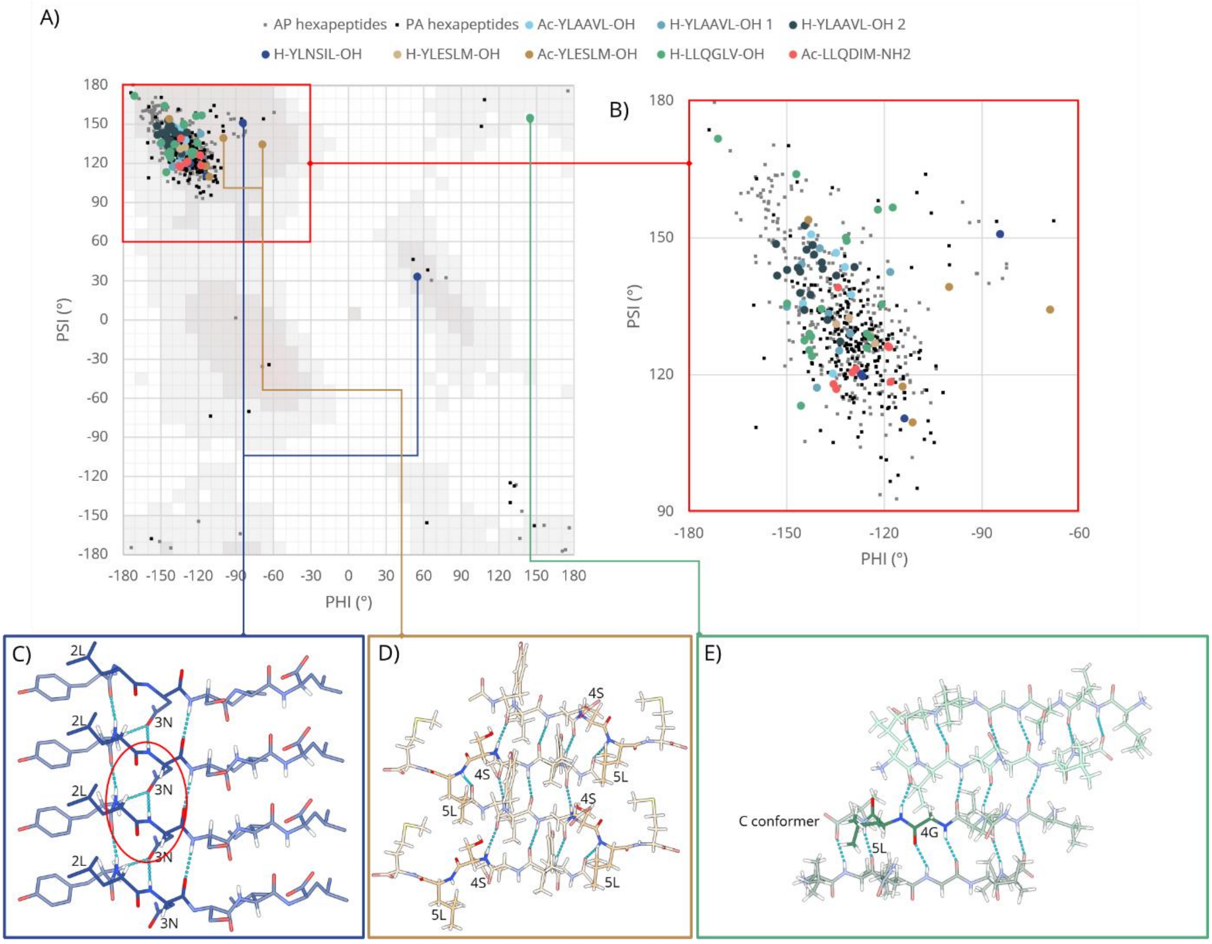


**Supplementary Figure 16. Ramachandran diagram of the amyloid-like crystal structures of APRs in the PACAP family. (A)** Amyloid-like crystal structures available in the PDB are shown in gray (antiparallel) and black (parallel β-sheet assemblies). APR structures of the PACAP family determined in this study are highlighted in distinct colours. **(B)** Enlarged view of the Ramachandran region characteristic of β-sheet backbone torsion angles. **(C–E)** Structural representation of residues whose backbone torsion angles fall outside the canonical β-sheet region. **C)** The torsion angle between Leu2 and Asn3 in $APR^{VIP}$ introduces an unusual twist in the β-strand plane, whereby the backbone amide of Asn3 forms a hydrogen bond with the side-chain amide of an Asn residue from a vertically adjacent β-strand, instead of participating in the canonical backbone–backbone hydrogen bond. **D)** In ${}_{ac}APR^{PHM}$, the peptide bond planes at the Ser4–Leu5 segment rotate by approximately 90° relative to the plane of the β-sheeted peptide chain, deviating from the geometry of a standard β-strand. **E)** In chain C of $APR^{SCT}$, the torsion angle between Gly4 and Leu5 also falls outside the canonical β-sheet region; however, this outlier conformation is consistent with the inherent torsional flexibility of glycine residues.

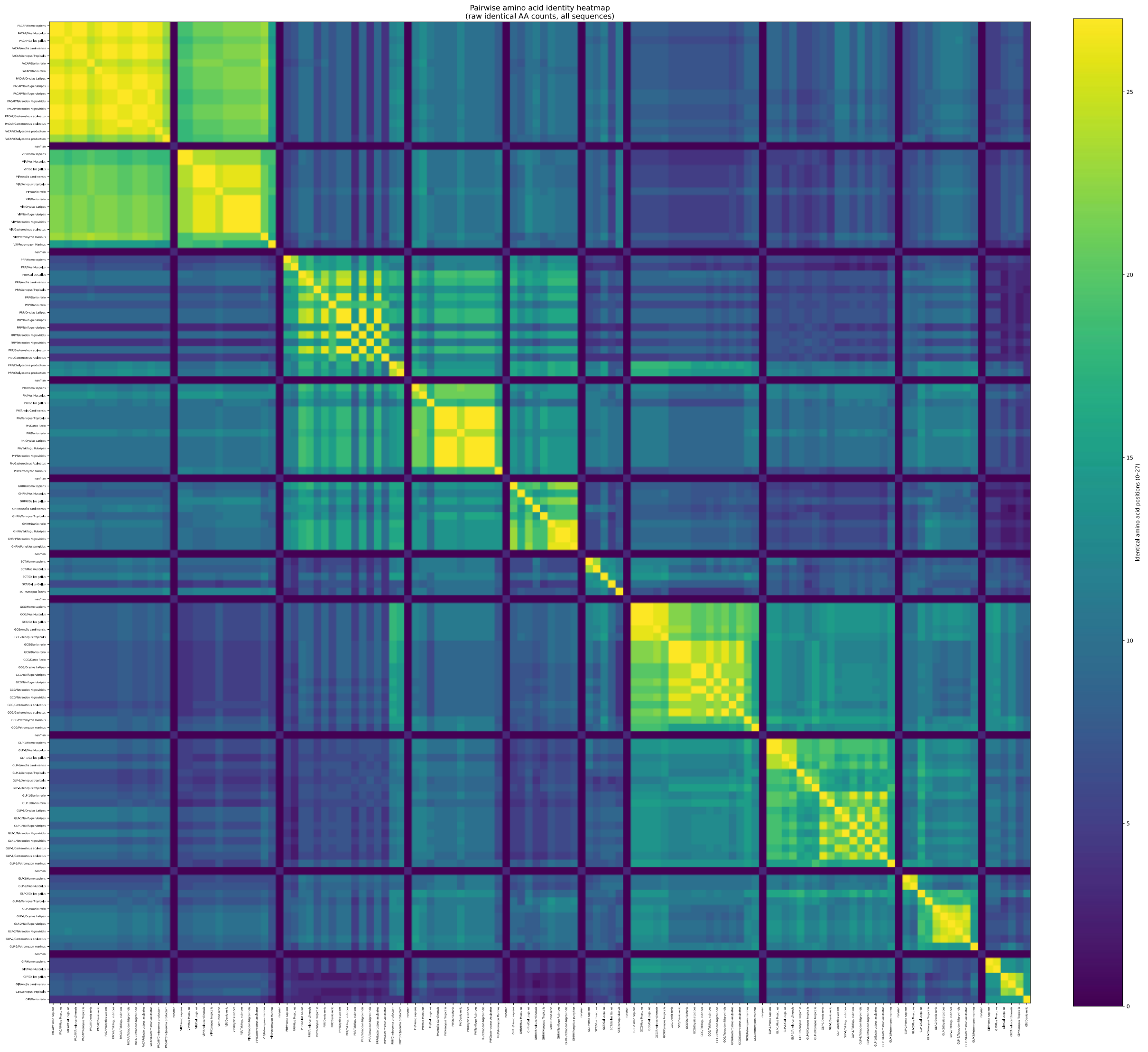
Pairwise amino acid identity heatmap
(raw identical AA counts, all sequences)
Identical amino acid positions (0–27)

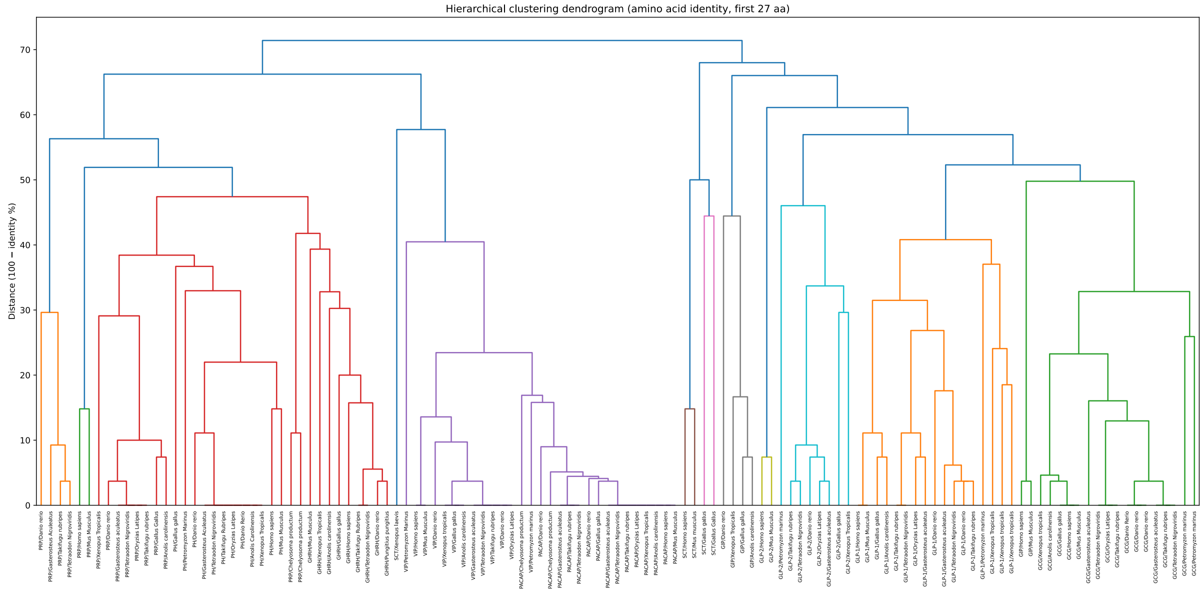
Hierarchical clustering dendrogram (amino acid identity, first 27 aa)
Distance (100 − identity %)

**Supplementary Figure 17. Pairwise amino acid identity heatmap of the PACAP–glucagon peptide family and hierarchical clustering dendrogram based on amino acid sequence identity (first 27 residues). (A)** The heatmap displays pairwise amino acid identity calculated for the N-terminal 27 residues of each peptide. Identity values represent the absolute number of identical residues at identical positions, with a maximum possible value of 27. Sequences are arranged according to their original order in the input dataset (S.Table 2, according to Cardoso et al.[5]) without hierarchical clustering or reordering. Thus, the matrix reflects the predefined taxonomic or family organization of the table rather than similarity-driven grouping. The color scale corresponds to the number of identical residues (range: 0–27). The diagonal represents self-comparisons (27 identical residues). Paralogous peptide copies within the same species often display lower sequence similarity than orthologous sequences from closely related species, producing a characteristic checkerboard-like pattern near the diagonal of the pairwise identity heatmap. This pattern reflects the exon-duplication–driven diversification of peptide copies discussed in the main text. **(B)** Percent identity was defined as the proportion of identical residues over 27 positions. A distance matrix was constructed as: $\text{Distance} = 100 - \text{identity}\,(\%)$ Hierarchical clustering was performed using the average linkage method. The dendrogram represents the relative evolutionary proximity of sequences at the amino acid level, with branch lengths proportional to sequence divergence. Sequences clustering together share higher positional identity across the conserved 27-residue region. The amino acid–based dendrogram reveals clear clustering according to peptide family, reflecting strong conservation at the protein level. PACAP/VIP sequences form a tight cluster characterized by short branch lengths, indicating minimal divergence within the conserved 27-residue region. Similarly, PRP/GHRH-related peptides group together, forming distinct subclusters separated from the PACAP/VIP branch. Mammalian PRP sequences are markedly separated from the other members of the family, which may be related to the emergence of an N-terminal one-residue extension in mammals resulting in the loss of the peptide's original function. Glucagon-related sequences form a separate branch, reflecting their evolutionary divergence from the PACAP lineage at the amino acid level Interestingly, the *Xenopus Laevis* secretin sequence shows higher similarity to PACAP/VIP peptides, whereas avian and mammalian SCTs more closely resemble GIP sequences, leaving the evolutionary origin of the secretin family unclear.

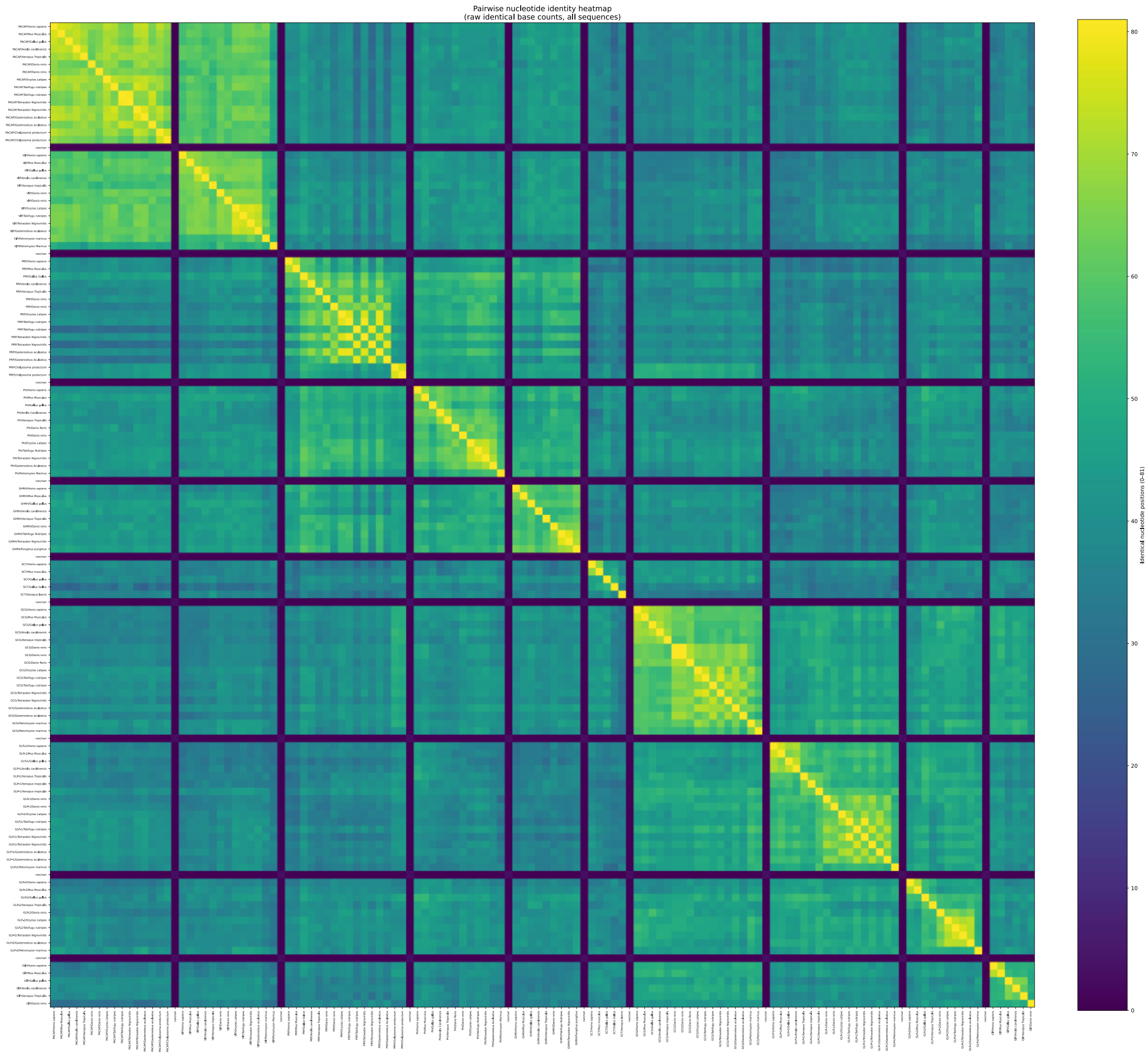
Pairwise nucleotide identity heatmap
(raw identical base counts, all sequences)
Identical nucleotide positions (0–81)
80
70
60
50
40
30
20
10
0

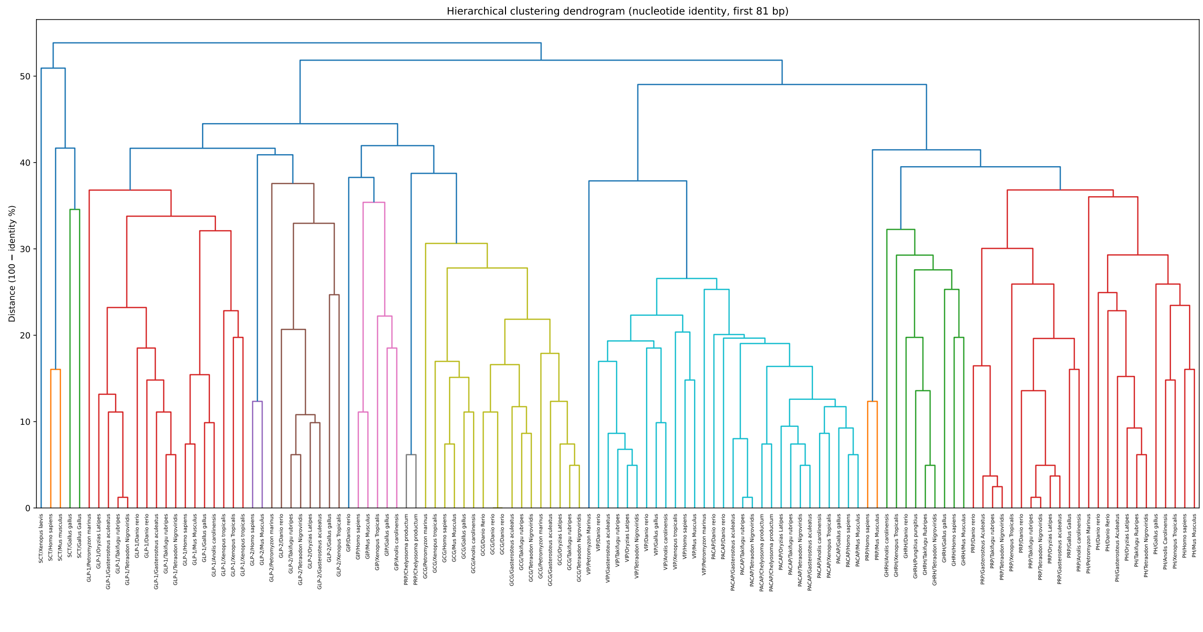
Hierarchical clustering dendrogram (nucleotide identity, first 81 bp)
Distance (100 − identity %)
50
40
30
20
10
0

**Supplementary Figure 18. Pairwise nucleotide identity heatmap of the PACAP–glucagon gene family and hierarchical clustering dendrogram based on nucleotide sequence identity (first 81 bp). (A)** The heatmap shows pairwise nucleotide identity calculated for the first 81 base pairs corresponding to the coding region of the N-terminal 27 amino acids. Identity values represent the absolute number of identical nucleotides at identical positions with a maximum possible value of 81. Sequences are displayed in the original order of the dataset, without hierarchical clustering. Therefore, similarity relationships must be interpreted within the predefined sequence arrangement rather than inferred from reordering. (S.Table 2, according to Cardoso et al.[5]) The color scale indicates the number of identical nucleotide positions (range: 0–81). The diagonal corresponds to self-comparisons (81 identical nucleotides). Unlike amino acid–level comparisons, nucleotide-level identity reflects both synonymous and nonsynonymous substitutions, enabling visualization of codon-level diversification within the PACAP–glucagon gene family. **(B)** Identity values were determined by direct position-by-position comparison without alignment and expressed as percentage identity across 81 positions. A distance matrix was computed as: Distance = 100 − identity (%) Hierarchical clustering was performed using the average linkage algorithm. Branch lengths reflect nucleotide-level divergence, capturing both synonymous and nonsynonymous substitutions. In contrast to amino acid dendogram (SFig.17), the nucleotide-based dendrogram displays greater branch length variability and increased dispersion within clusters. Although major family-level groupings remain observable, internal divergence within PACAP-related sequences is more pronounced at the nucleotide level than at the amino acid level. This pattern reflects extensive synonymous codon diversification: sequences encoding highly conserved peptides show substantial nucleotide-level variation while preserving identical or near-identical amino acid sequences. (SFig. 19) Although the pairwise sequence heatmap (SFig. 17) suggests partial similarity between secretin and other peptide families, nucleotide-based clustering places the five analysed SCT sequences as a distinct lineage, clearly separated from the PACAP–glucagon family. This may indicate that the observed similarities arise from convergent evolution rather than shared ancestry.

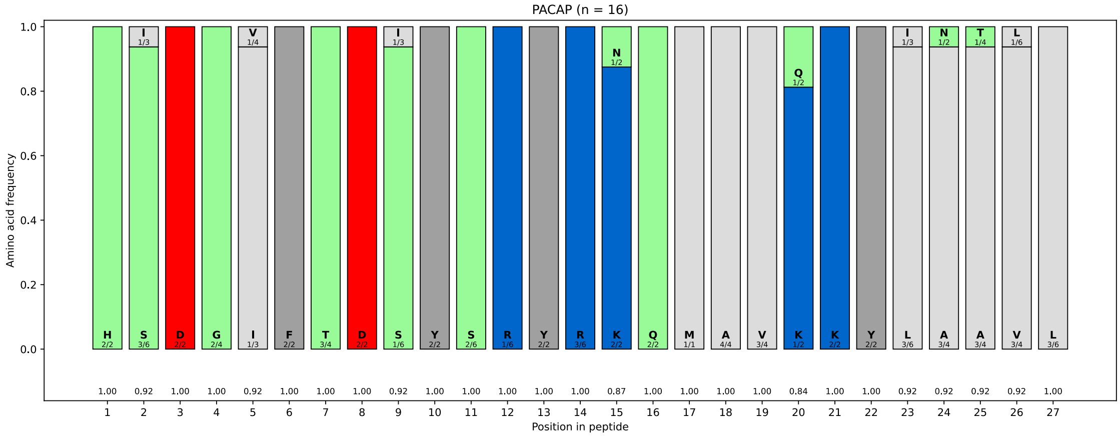


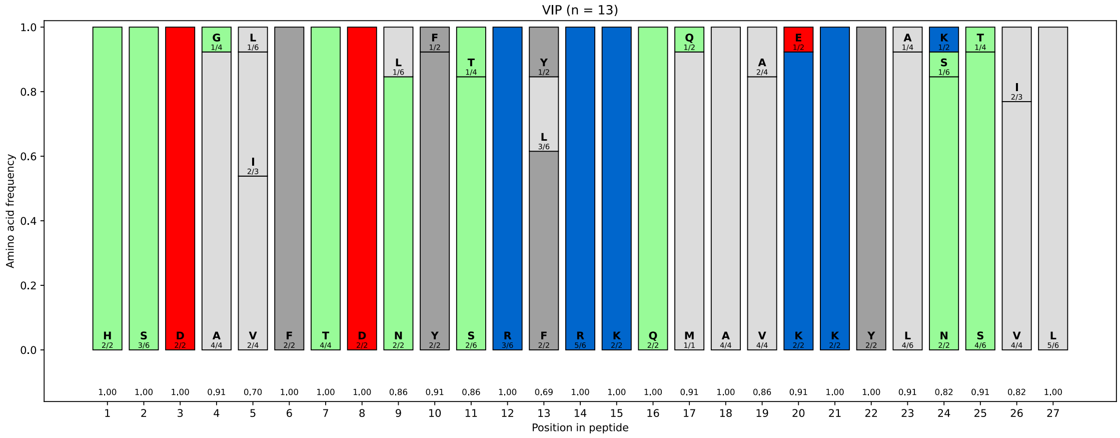

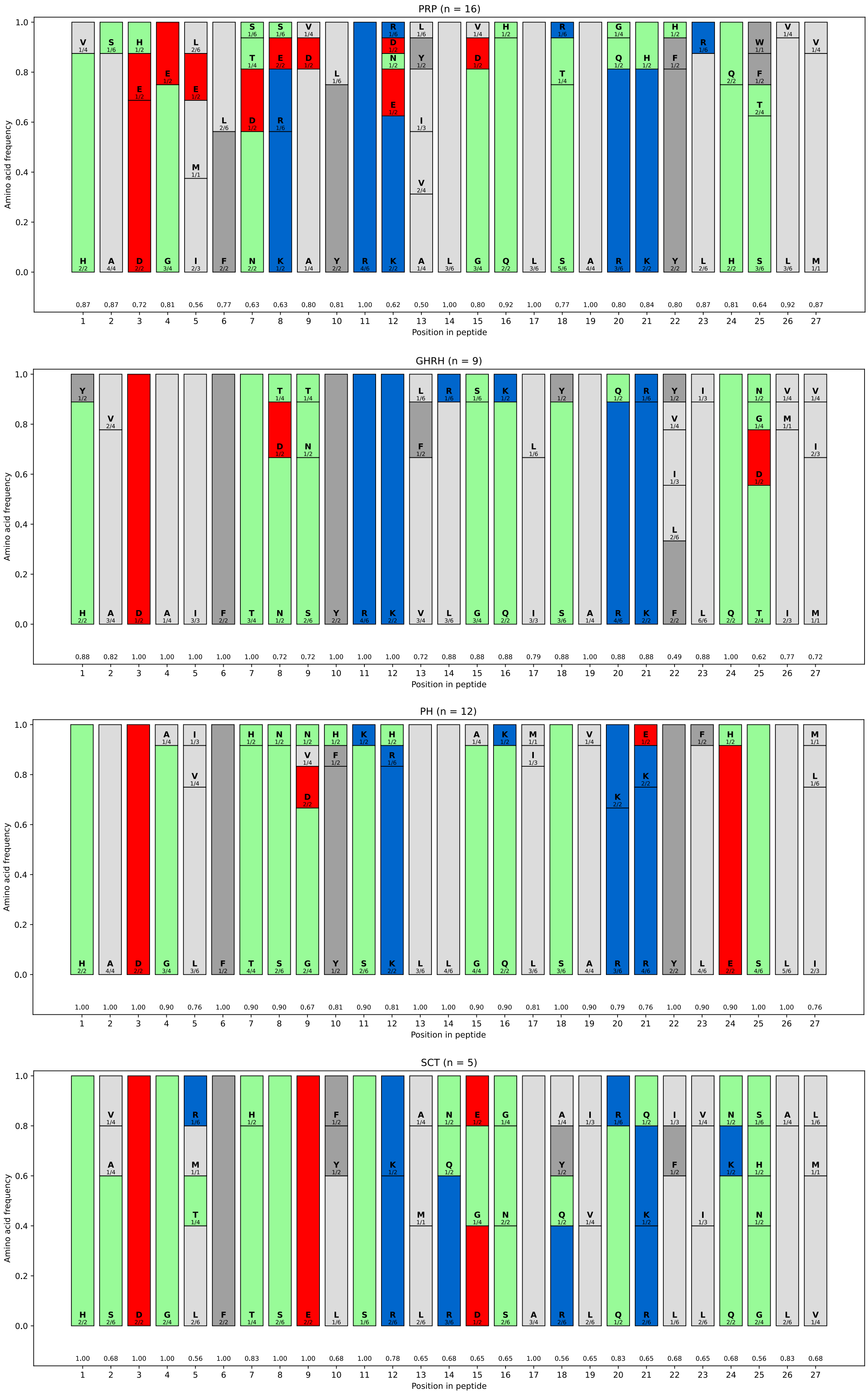
PRP (n = 16)
GHRH (n = 9)
PH (n = 12)
SCT (n = 5)
Amino acid frequency
Position in peptide

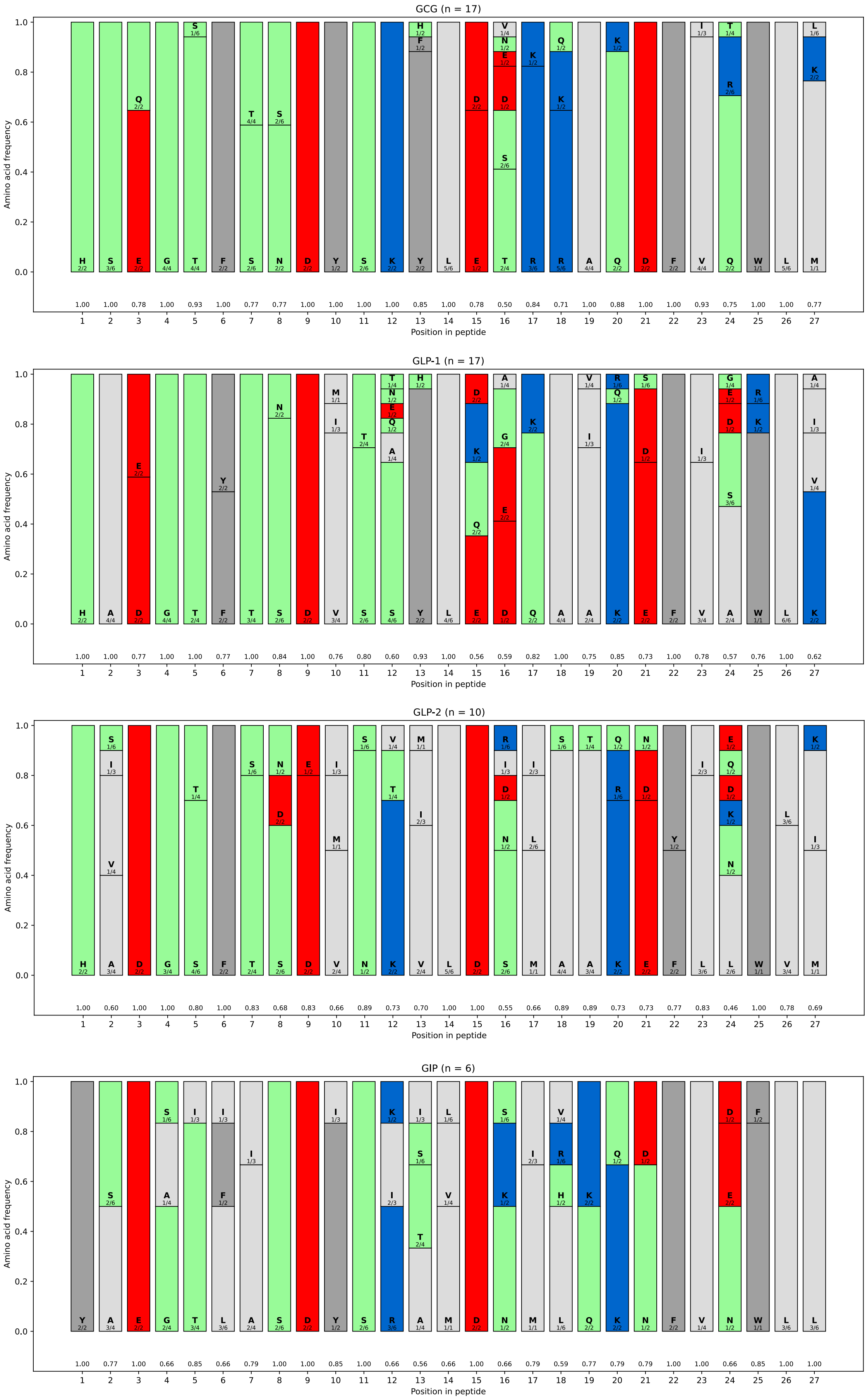
GCG (n = 17)
GLP-1 (n = 17)
GLP-2 (n = 10)
GIP (n = 6)
Amino acid frequency
Position in peptide

**Supplementary Figure 19. Position-specific amino acid conservation and codon usage diversity across peptide families of the PACAP–glucagon superfamily.** These plots simultaneously illustrate position-specific amino acid conservation, physicochemical residue classes, and codon usage diversity across peptide families, enabling comparison of evolutionary constraints at both the protein and coding-sequence levels.For each peptide family, the N-terminal 27 residues were analysed across all selected sequences. (S.Table 2.) The number of sequences analyzed for each peptide family is indicated above the corresponding bar chart (n). At each position, amino acid frequencies were calculated and visualized as stacked bars, where the height of each colored segment represents the relative frequency of a given amino acid among the analysed species. Amino acids are color-coded according to physicochemical properties: acidic residues (D, E) are shown in red, basic residues (K, R) in blue, polar residues in light green, hydrophobic residues (A, V, L, I, M, P) in light gray, and aromatic hydrophobic residues (F, W, Y) in dark gray. Individual amino acid identities are indicated above the corresponding bar segments. For each amino acid observed at a given position, codon usage diversity was evaluated from the corresponding DNA sequences. The annotation shown within each segment indicates the number of distinct codons observed among the selected sequences relative to the theoretical maximum number of synonymous codons for that amino acid (observed codons / maximum codon degeneracy). This representation allows direct visualization of how extensively the synonymous codon space is utilized at each position. To quantify positional conservation, a normalized Shannon conservation index (CI) was calculated based on the observed amino acid frequency distribution at each position. The CI values are displayed below the bars, where values approaching 1 indicate strong conservation and values approaching 0 reflect higher sequence variability.

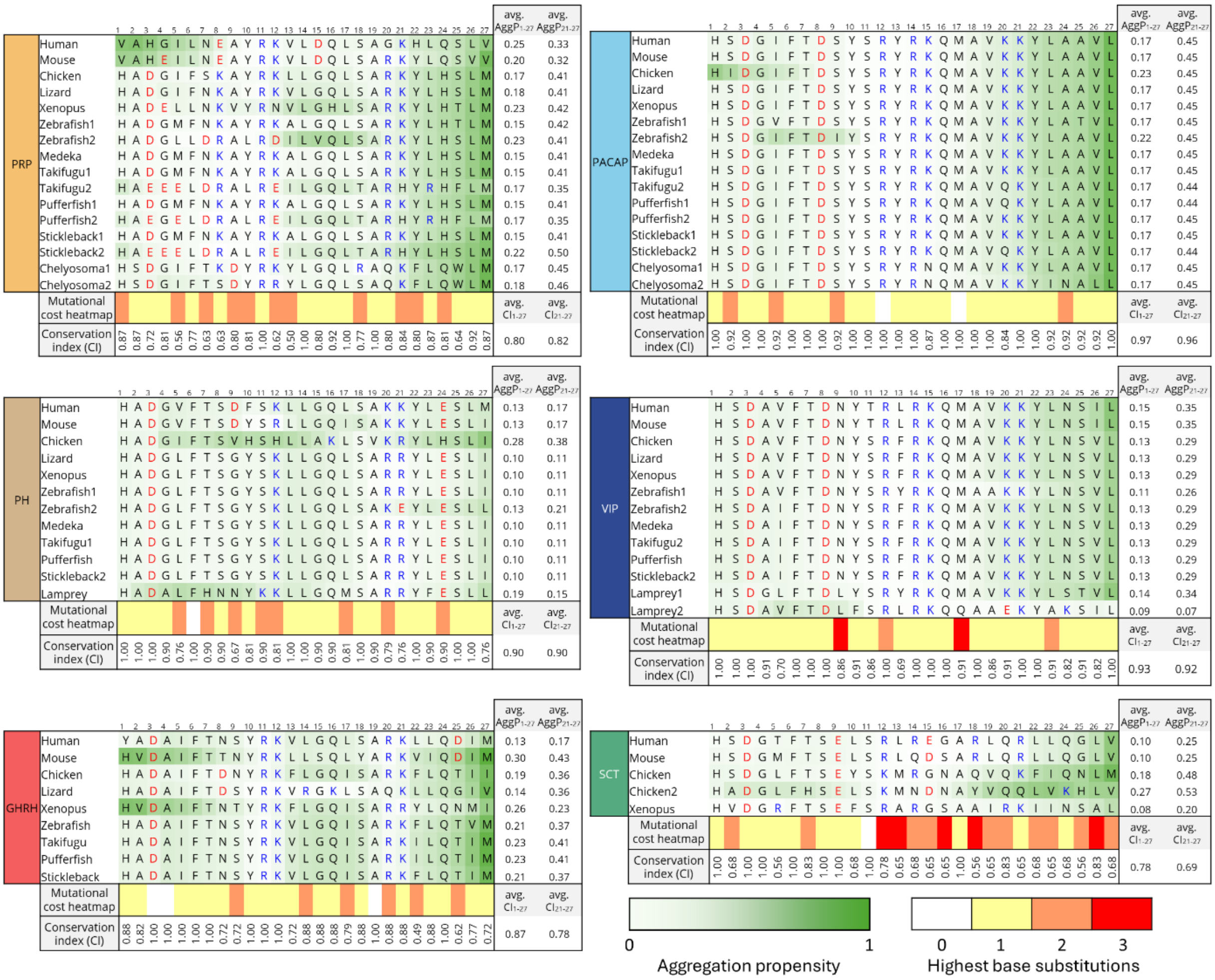


**Supplementary Figure 20. Position-specific conservation and aggregation propensity across species in the PACAP family.**

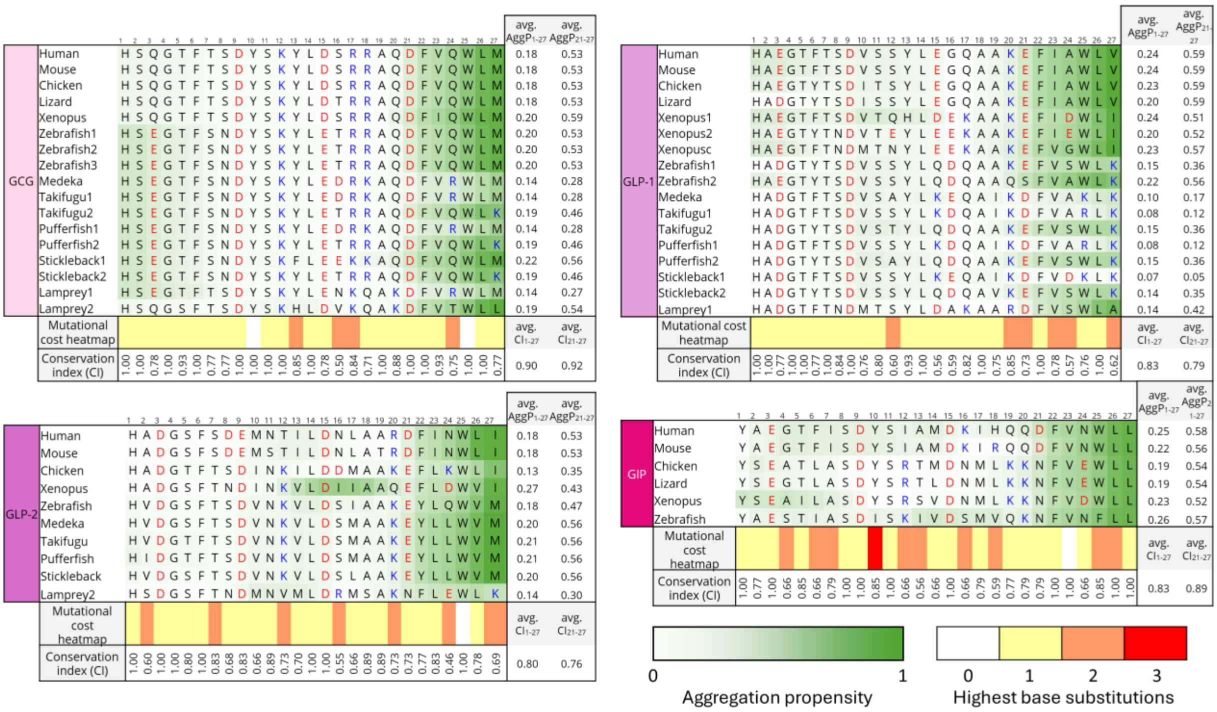


**Supplementary Figure 20. (continued) Position-specific conservation and aggregation propensity across species in the glucagon family.** Species are arranged from bottom to top according to evolutionary order. Position-specific aggregation scores were calculated using AggreProt for each peptide family across the analyzed species. Although the aggregation heatmaps indicate the occasional appearance of local aggregation hotspots in certain species, residues 21–27 consistently represent the characteristic aggregation-prone region (APR) of the PACAP–glucagon peptide family. For each sequence, the residue-normalized mean aggregation propensity of the full peptide (avg. AggP$_{1–27}$) and of the APR region (avg. AggP$_{21–27}$) is indicated at the sequence end. Previous predictive and experimental studies in human sequences showed that the aggregation hotspot of the glucagon family forms a stronger APR than that of the PACAP family. The avg. AggP$_{21–27}$ values demonstrate that this difference is conserved across vertebrate evolution: values for the glucagon family are typically around 0.5, whereas those for the PACAP family are generally below 0.45. In the PH family, avg. AggP$_{21–27}$ values are generally lower than in the other peptide families and do not alone clearly indicate APR character; however, our experimental results with the human $_{ac}$APR$^{PHM}$ peptide confirm the presence of an aggregation-prone nature of this segment. Position-specific conservation (CI) of the most frequent residues across species are shown. Residue-normalized mean CI of the full peptide (avg. CI$_{1–27}$) and of the APR region (avg. CI$_{21–27}$) are shown at the right bottom for each table. Overall, the similarity of the two values indicates that the receptor-binding, aggregation-relevant 21–27 region is not more conserved than the full peptide sequence. Mutational cost heatmap indicating the maximal nucleotide substitution distance among the codons observed at each position; the highest observed distance is highlighted even when most codon pairs differ by fewer substitutions. The heatmap based solely on the codons actually detected in the analysed sequences. A value of 0 indicates complete conservation, whereas values of 1–3 represent increasing mutational cost. The number of the highest observed base substitutions were determined according to the position-specific mutational pathways are listed in S.Table 4/1-10.

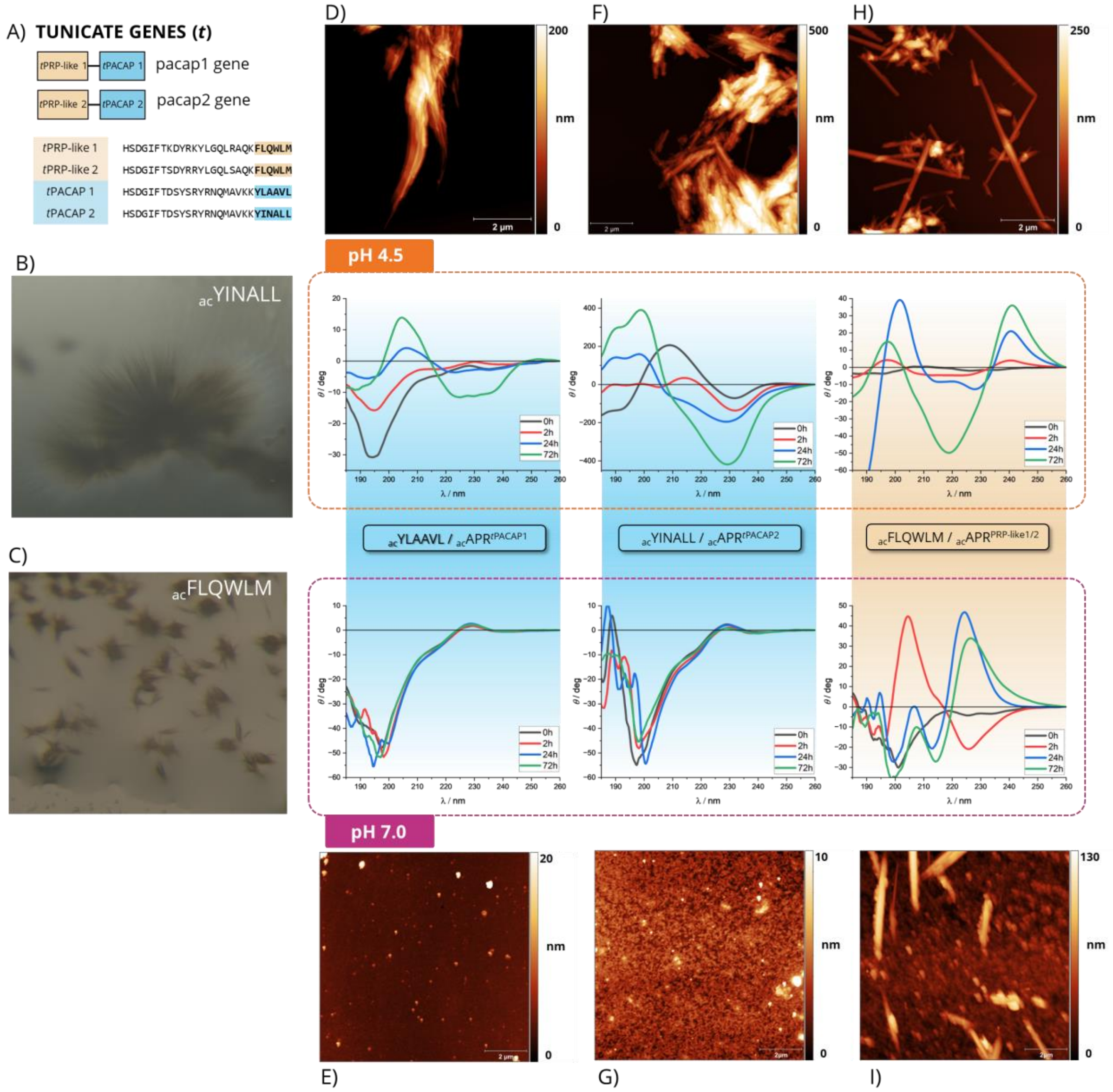


**Supplementary Figure 21. Amyloid aggregation analysis of PACAP-related APR hexapeptides from *Chelyosoma productum*. (A)** In the disk-topped tunicate *Chelyosoma productum*, two full-length cDNAs encode four 27-residue peptides with high similarity to the products of the human ADCYAP1 gene, namely PACAP and PRP. **(B–C)** Microcrystals were obtained from all three APR-derived hexapeptides; however, crystals of $_{ac}$APR$^{tPACAP\ 2}$ and $_{ac}$APR$^{PRP\text{-}like\ 1/2}$ were not suitable for diffraction data collection.**(D–I)** Structural conversion and self-association of the APR segments were monitored by CD and AFM over a 72 h incubation at 37 °C under continuous stirring (magnetic bar) at pH 4.5 (upper panels) and pH 7.0 (lower panels) at a peptide concentration of 5 mg ml$^{-1}$. Panels D–E show the APR of *t*PACAP 1, F–G the APR of *t*PACAP 2, and H–I the APR of *t*PRP 1/2. Amyloid-like self-association was observed at pH 4.5, where the C-termini of the hexapeptides remain largely protonated. Under neutral conditions (pH 7.0), when the C-termini carry a negative charge, the PACAP-related sequences remained largely disordered. For the $_{ac}$YINALL peptide, incubation at pH 4 resulted in a pronounced increase in CD intensity accompanied by structural transitions characteristic of β-type CD spectra. In contrast with the PACAP-related sequences, the FLQWLM peptide displayed partially distinct behaviour. In addition to the β-type CD signature, a pronounced negative–positive band pair around 220 nm appeared, previously reported (HIV) for glucagon-family peptides and attributed to aromatic Phe–Trp interactions. This observation is consistent with our hierarchical clustering analysis, which revealed that the tunicate PRP-like peptide shows substantial similarity to the glucagon peptide family. This relationship becomes even more evident when only the APR segments are compared, as the APR$^{hGCG}$ (FVQWLM) closely resembles the tunicate PRP-like sequence.

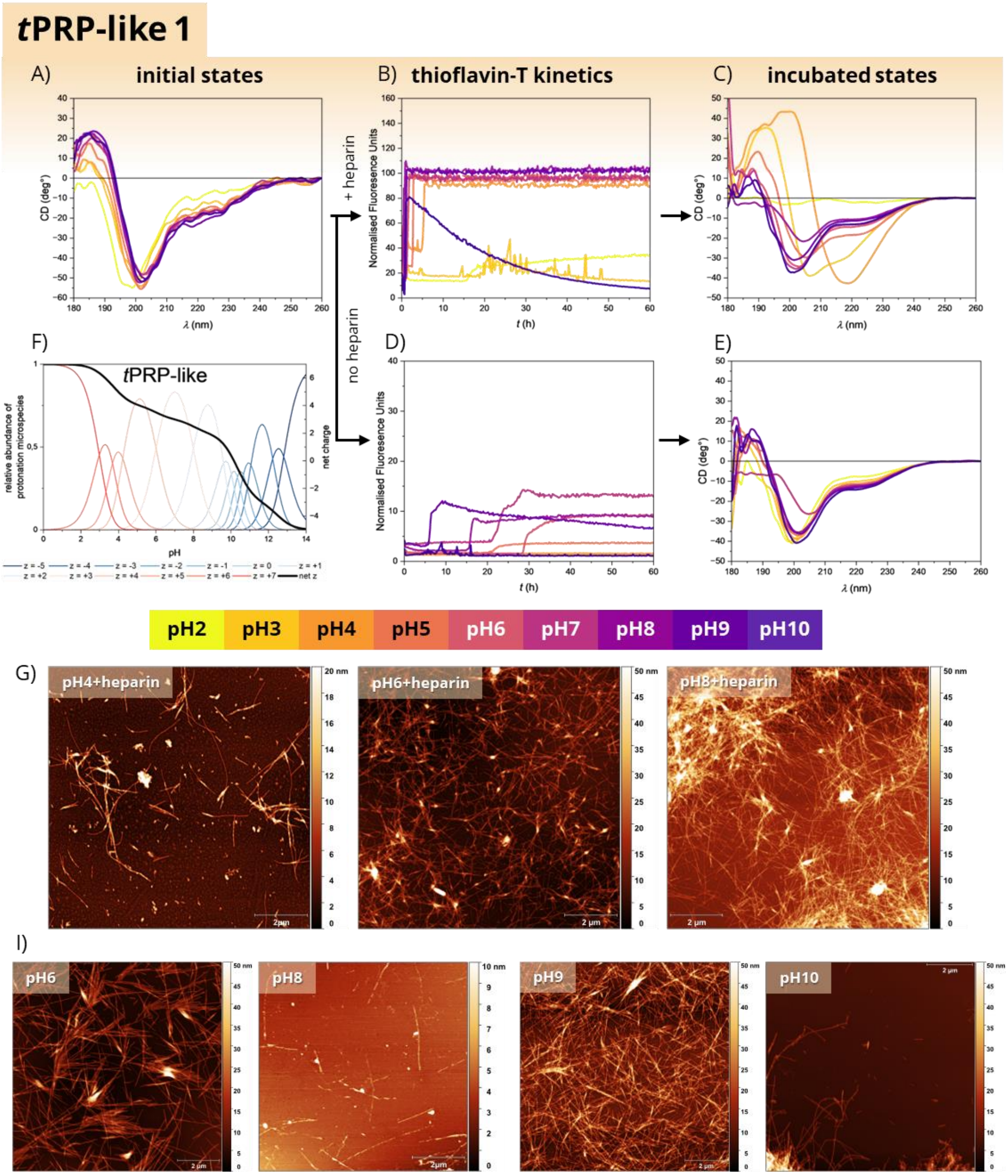


**Supplementary Figure 22. Monitoring of the amyloid formation for tunicate derived PRP-like 1 peptide in presence and absence of heparin**. The experimental setup was identical to that used for the human sequences. A general description of the experimental conditions is provided in the figure legend on page 2 of the Supplementary; here we focus on *t*PRP-like 1 peptide specific observations. AFM images **(G)** confirm that in the pH range 3–9, where the net peptide charge (+z) is positive **(F)**, rapid aggregation (<5 h) occurs in the presence of heparin **(B)**, as indicated by ThT kinetics. During this process, the structural transition from a disordered to a β-sheet–rich state is only partially reflected in the CD spectra and is clearly detectable mainly in the pH 3–6 range **(C)**. In contrast to the human PACAP family sequences, AFM scans demonstrate that amyloid formation can also occur in the absence of heparin within the pH range 5–10 **(I)**. However, according to the ThT kinetics **(D)** this process is slower than in the presence of heparin and the measured fluorescence intensity reaches only a fraction of that observed in heparin-containing samples, while the CD spectra **(E)** do not clearly indicate the corresponding structural transition.

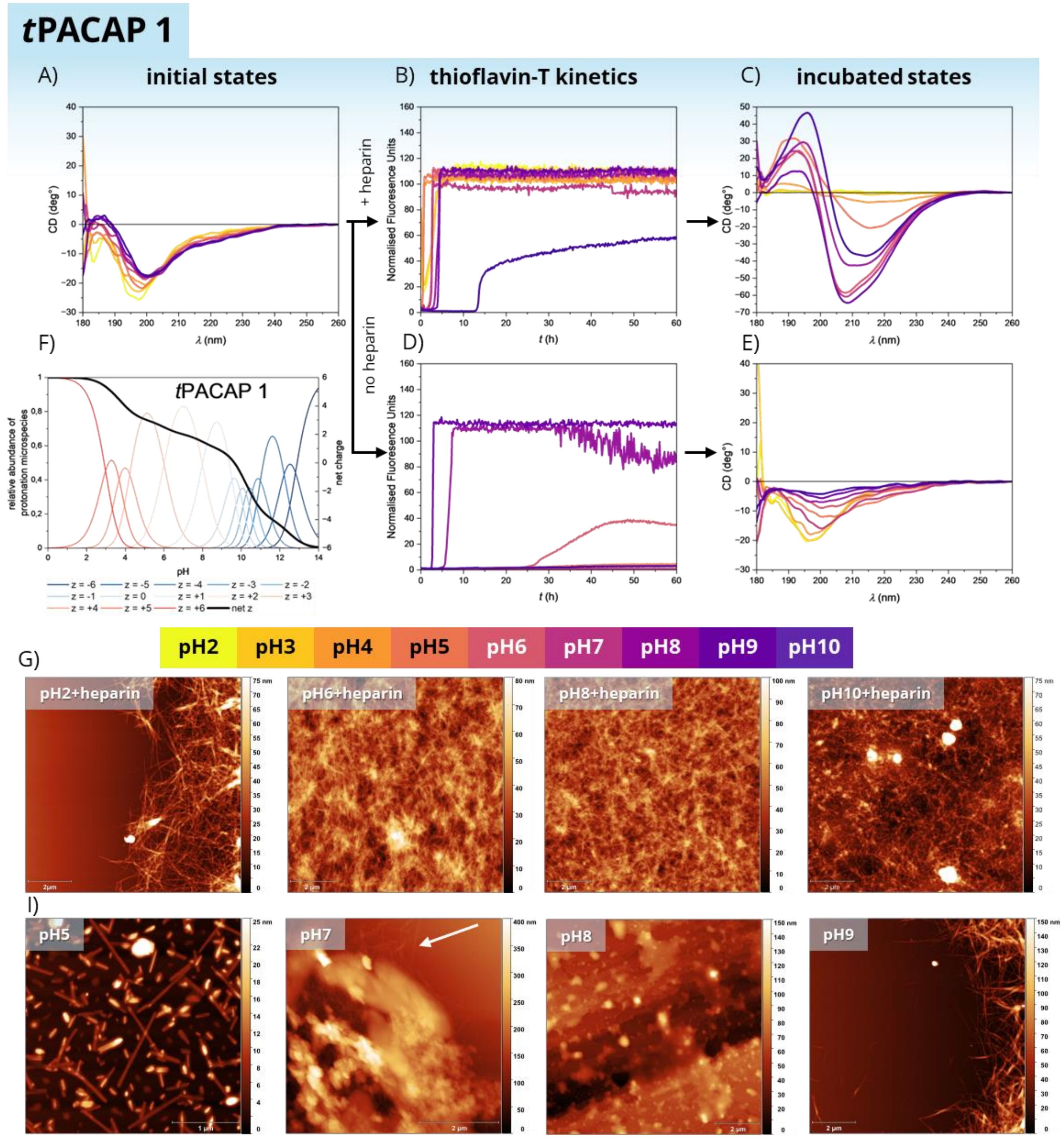


**Supplementary Figure Z4. Monitoring of the amyloid formation for tunicate derived PACAP 1 peptide in presence and absence of heparin**. The experimental setup was identical to that used for the human sequences. A general description of the experimental conditions is provided in the figure legend on page 2 of the Supplementary; here we focus on *t*PACAP 1 peptide specific observations. AFM images **(G)** confirm that in the whole investigated pH range 2–10, where the net peptide charge (+z) remains positive **(F)**, rapid aggregation (<5 h) occurs in the presence of heparin **(B)**, as indicated by ThT kinetics. CD spectroscopy also indicates structural conversion into β-sheet–rich conformations. The 27-residue *t*PACAP1 peptide differs from human PACAP-27 by only a single amino acid and lacks the Lys/Arg-rich C-terminal extension of human PACAP-37. Unlike PACAP-37, the tunicate peptide forms amyloid fibrils even in the absence of heparin. CD spectra reveal a pH-dependent gradient in structural behavior. At acidic pH, the spectra retain a profile characteristic of disordered conformations, whereas toward alkaline conditions the decreasing CD intensity indicates increasing precipitation of the peptide. ThT fluorescence increase was detected only in the pH 6, 7, and 9 samples, where AFM imaging confirmed the presence of fibrillar structures, often partially obscured by amorphous aggregates (arrow). The absence of detectable aggregation at pH 8 remains unexplained. In the heparin-free pH 5 sample, although neither CD nor ThT indicated aggregation, AFM revealed a uniform, fully linear rod-like amyloid morphology, distinct from the fibrillar structures observed at the other pH conditions.

### PACAP

| | |
|---|---|
| human/sheep /rat/mouse | HSDGI FTDSY SRYRK QMAVK KYLAA VLGKR YKQRV KNK |
| chicken | HIDGI FTDSY SRYRK QMAVK KYLAA VLGKR YKQRV KNK |
| lizard | HSDGI FTDSY SRYRK QMAVK KYLAA VLGKR YKQ** *** |
| green frog | HSDGI FTDSY SRYRK QMAVK KYLAA VLGKR YKQRI KNK |
| salmon | HSDGI FTDSY SRYRK QMAVK KYLAA VLGKR YRQRY RNK |
| catfish | HSDGI FTDSY SRYRK QMAVK KYLAA VLGKR YRQRF RNK |
| stargazer | HSDGI FTDSY SRYRK QMAVQ KYLAA VLGKR YRQRV RNK |
| stingray | HSDGI FTDSY SRYRK QMAVK KYLAA VLGKR YKPKV KNS/G RRVFY |
| tunicate-I | HSDGI FTDSY SRYRN QMAVK KYLAA VL |
| tunicate-II | HSDGI FTDSY SRYRN QMAVK KYINA LL |

### SECRETIN

| | |
|---|---|
| human | HSDGT FTSEL SRLRE GARLQ RLLQG LV* |
| pig | HSDGT FTSEL SRLRD SARLQ RLLQG LV* |
| dog | HSDGT FTSEL SRLRE SARLQ RLLQG LV* |
| guinea pig | HSDGT FTSEL SRLRD SARLQ RLLQG LV* |
| rat | HSDGT FTSEL SRLQD SARLQ RLLQG LV* |
| mouse | HSDGM FTSEL SRLQD SARLQ RLLQG LV* |
| chicken | HSDGL FTSEY SKMRG NAQVQ KFIQN LM* |
| X.laevis | HVDGR FTSEF SRARG SAAIR KIINS ALA |
| X.tropicalis | HVDGM FTSEF SRARG SAAIR KIINS ALA |

### PRP

| | |
|---|---|
| human | DVAHGI LNEAY RKVLD QLSAG KHLQS LVARG VGGSL GGGAG DDA*E PLS |
| sheep | DVAHGI LDKAY RKVLD QLSAR RYLQT LMAKG LGGYP GGGAD DDS*E PLS |
| rat | DVAHEI LNEAY RKVLD QLSAR KYLQS MVARG MGENL AAAAV DDR*A PLT |
| mouse | DVAHEI LNEAY RKVLD QLSAR KYLQS VVARG AGDEP RRHAV DDP*A PLT |
| chicken I | *HADGI FSKAY RKLLG QLSAR KYLHS LMAKR VGG** ASSGL GDEAE PLS |
| chicken II | *HADGI FSKAY RKLLG QLSAR KYLHS LMAKR VG*** **SGL GDEAE PLS |
| xenopus | *HADEL LNKVY RNVLG HLSAR KYLHT LMAQR LGT** VSSSL EDESE PLS |
| carp | *HADGM FNKAY RKALG QLSAR KYLHT LMAKR VG*** GGSMI EDDNE PLS |
| catfish | *HADGL LDRAL RDILV QLSAR KYLHS LTAVR VG*** EEEED EEDSE PLS |

| | |
|---|---|
| goldfish (salmon-like) | *HADGM FNKAY RKALG QLSAR KYLHT LMRKR VG*** GGSTI EDDNE PLS |
| goldfish (catfish-like) | *HADGL LDRAL RDILV QLSAR KYLHS LMAVR VG*** GGSSE EDESE PLS |
| salmon | *HADGM FNKAY RKALG QLSAR KYLHS LMAKR VG*** GGSTM EDDSE PLS |
| zebrafish | *HADGL LDRAL RDILV QLSAR KYLHS LMAVR VG*** GGSSE EDESE PLS |
| tunicate-I | *HSDGI FTKDY RKYLG QLRAQ KFLQW LM |
| tunicate-II | *HSDGI FTSDY RRYLG QLSAQ KFLQW LM |

### VIP

| | |
|---|---|
| human/monkey/ pig/cattle/dog/ rat/mouse | HSDAV FTDNY TRLRK QMAVK KYLNS ILN |
| guinea-pig | HSDAL FTDTY TRLRK QMAMK KYLNS VLN |
| opossum | HSDAV FTDSY TRLLK QMAMR KYLDS ILN |
| chicken/turkey/ alligator/frog | HSDAV FTDNY SRFRK QMAVK KYLNS VLT |
| cod | HSDAV FTDNY SRFRK QMAAK KYLNS VLA |
| trout/bowfin | HSDAV FTDNY SRFRK QMAVK KYLNS VLT |
| dogfish | HSDAV FTDNY SRIRK QMAVK KYINS ILA |

### PHI/PHM

| | |
|---|---|
| human | HADGV FTSDF SKLLG QLSAK KYLES LM |
| rat/mouse | HADGV FTSDY SRLLG QISAK KYLES LI |
| guinea-pig | HADGV FTSDY SRLLG QLSAR KYLES LI |
| cow/sheep | HADGV FTSDY SRLLG QLSAK KYLES LI |
| pig/rabbit | HADGV FTSDF SRLLG QLSAK KYLES LI |
| chicken | HADGI FTSVY SHLLA KLAVK RYLHS LI |
| turkey | HADGI FTTVY SHLLA KLAVK RYLHS LI |
| goldfish | HADGL FTSGY SKLLG QLSAK KYLE SLL |

### GHRH

| | |
|---|---|
| human | YADAI FTNSY RKVLG QLSAR KLLQD IMSRQ QGESN QERGA RARL |
| pig | YADAI FTNSY RKVLG QLSAR KLLQD IMSRQ QGERN QEQGA RVRL |
| cattle/goat | YADAI FTNSY RKVLG QLSAR KLLQD IMNRQ QGERN QEQGA KVRL |
| sheep | YADAI FTNSY RKILG QLSAR KLLQD IMNRQ QGERN QEQGA KVRL |

| | |
|---|---|
| hamster | YADAI FTSSY RKVLG QLSAR KLLQD IMSRQ QGERN QEQGP RVRL |
| rat | HADAI FTSSY RRILG QLYAR KLLHE IMNRQ QGERN QEQRS RFN |
| mouse | EVDAI FTTNY RRILS QTYAR KVIQD IMNKQ *GERI QEQRA RLS |
| chicken GRF1-46 | HADGI FSKAY RKTLG QLSAR NYLHS LMAKR VGGAS SGLGD EAEPL S |
| Chicken GRF1-43 | HADGI FSKAY RKTLG QLSAR NYLHS LMAKR VG*** SGLGD EAEPL S |
| atlantic salmon | HADGM FNKAY RKALG QLSAR KYLHS LMAKR VGGGS TMEDD SEPLS |
| chinook salmon | HADGM FNKAY RKALG QLSAR KYLHS LMAKR VGGGS TMEDD SEPLS |
| carp | HADGM FNKAY RKALG QLSAR KYLHT LMAKR VGGGS MIEDD NEPLS |
| catfish | HADGL LDRAL RDILV QLSAR KYLHS LTAVR VGEEE EDEED SEPLS |
| tunicate-I | HSDGI FTKDY RKYLG QLRAQ KFLQW LM |
| tunicate-II | HSDGI FTSDY RRYLG QLSAQ KFLQW LM |

### HUMAN GLUCAGON FAMILY MEMBERS

| | |
|---|---|
| glucagon | HSQGT FTSDY SKYLD SRRAQ DFVQW LMNT |
| GLP-1 | HAEGT FTSDV SSYLE GQAAK EFIAW LVKGR |
| GLP-2 | HADGS FSDEM NTILD NLAAR DFINW LIQTK ITD |
| GIP | YAEGT FISDY SIAMD KIHQQ DFVNW LLAQK GKKND WKHNI TQ |

**Supplementary Table 1. Members of the PACAP peptide family across vertebrate species.** Full-length sequences are arranged for each peptide type in evolutionary order (from bottom to top). For clarity, sequences are segmented into blocks of five amino acids and aligned to the human reference sequence (top row). Positively charged residues are highlighted in blue, and negatively charged residues in red. Asterisks (*) indicate alignment gaps introduced to maintain positional correspondence and do not represent amino acids. Due to their common evolutionary origin, GHRH (also referred to in the literature as GRF, growth hormone–releasing factor) and PRP share identical sequences in non-mammalian species; differences in segmentation reflect alignment to the human reference sequence rather than sequence variation. Sequences were collected from published literature sources.[5–8]

| Peptide | Species | Latin name | Sequence | SeqDNA |
| --- | --- | --- | --- | --- |
| PACAP | Human | Homo Sapiens | HSDGIFTDSYSRYRKQMAVKKYLAAVL | CACTCGGACGGGATCTTCACGGACAGCTACAGCCGCTACCGGAAACAAATGGCTGTCAAGAAATACTTGGCGGCCGTCCTA |
| PACAP | Mouse | Mus Musculus | HSDGIFTDSYSRYRKQMAVKKYLAAVL | CACTCGGACGGCATCTTCACAGATAGCTACAGCCGCTACCGAAAACAAATGGCTGTCAAGAAATACTTGGCGGCCGTGCTA |
| PACAP | Chicken | Gallus Gallus | HIDGIFTDSYSRYRKQMAVKKYLAAVL | CACATAGACGGCATCTTCACGGACAGCTACAGCCGCTACCGGAAACAAATGGCTGTCAAGAAATACTTAGCGGCCGTCCTG |
| PACAP | Lizard | Anolis Carolinensis | HSDGIFTDSYSRYRKQMAVKKYLAAVL | CATTCAGACGGCATCTTCACAGACAGCTACAGCCGCTATCGAAAACAAATGGCTGTCAAGAAATACTTGGCGGCAGTCCTG |
| PACAP | Xenopus | Xenopus Tropicalis | HSDGIFTDSYSRYRKQMAVKKYLAAVL | CATTCAGATGGCATCTTCACCGACAGCTACAGTCGCTACAGAAAACAAATGGCTGTCAAGAAATACTTGGCAGCAGTGCTG |
| PACAP | Zebrafish (a) | Danio Rerio | HSDGVFTDSYSRYRKQMAVKKYLATVL | CACTCGGACGGGGTTTTCACGGACAGCTACAGTCGCTACCGGAAGCAAATGGCCGTGAAGAAGTATCTGGCCACGGTCCTT |
| PACAP | Zebrafish (b) | Danio Rerio | HSDGIFTDIYSRYRKQMAVKKYLAAVL | CATTCGGATGGGATCTTCACCGACATTTACAGTCGCTACCGAAAACAGATGGCCGTCAAGAAGTATTTAGCAGCCGTCCTG |
| PACAP | Medeka | Oryzias Latipes | HSDGIFTDSYSRYRKQMAVKKYLAAVL | CACTCAGACGGGATCTTCACGGACAGCTACAGCCGCTACAGAAAGCAAATGGCCGTCAAGAAATACCTGGCAGCCGTCCTG |
| PACAP | Takifugu (a) | Takifugu Rubripes | HSDGIFTDSYSRYRKQMAVKKYLAAVL | CACTCAGACGGGATCTTTACAGACAGCTACAGCCGCTACCGAAAGCAAATGGCGGTCAAGAAGTACCTGGCGGCAGTCCTT |
| PACAP | Takifugu(b) | Takifugu Rubripes | HSDGIFTDSYSRYRKQMAVQKYLAAVL | CATTCTGATGGCATCTTCACCGACAGCTACAGCCGCTATAGAAAGCAGATGGCCGTGCAGAAATACCTGGCAGCCGTTCTG |
| PACAP | Pufferfish (a) | Tetraodon Nigroviridis | HSDGIFTDSYSRYRKQMAVQKYLAAVL | CATTCTGATGGCATCTTTACCGACAGCTACAGCCGCTATAGAAAGCAGATGGCCGTGCAGAAATACCTGGCAGCCGTTCTG |
| PACAP | Pufferfish (b) | Tetraodon Nigroviridis | HSDGIFTDSYSRYRKQMAVKKYLAAVL | CACTCGGATGGGATCTTCACAGACAGCTACAGCCGCTACCGAAAGCAAATGGCAGTCAAGAAATACCTGGCGGCAGTCCTT |
| PACAP | Stickleback (a) | Gasterosteus Aculeatus | HSDGIFTDSYSRYRKQMAVKKYLAAVL | CACTCGGACGGCATCTTCACGGACAGCTACAGCCGCTACCGAAAGCAAATGGCAGTCAAGAAATACCTGGCGGCGGTCCTT |
| PACAP | Stickleback (b) | Gasterosteus Aculeatus | HSDGIFTDSYSRYRKQMAVQKYLAAVL | CACTCAGATGGGATCTTCACCGACAGCTACAGTCGCTATAGAAAGCAGATGGCCGTGCAGAAATACCTGGCGGCGGTTCTG |
| PACAP | Chelyosoma 1 | Chelyosoma Productum | HSDGIFTDSYSRYRNQMAVKKYLAAVL | CACTCGGATGGGATCTTCACGGACAGCTATAGCCGCTACCGGAATCAAATGGCTGTTAAGAAATACCTGGCGGCAGTGCTT |
| PACAP | Chelyosoma 2 | Chelyosoma Productum | HSDGIFTDSYSRYRNQMAVKKYINALL | CACTCGGATGGGATCTTCACGGACAGCTATAGCCGCTACCGGAATCAAATGGCTGTTAAGAAATACATAAACGCACTGCTT |
| **Peptide** | **Species** | **Latin name** | **Sequence** | **SeqDNA** |
| VIP | Human | Homo Sapiens | HSDAVFTDNYTRLRKQMAVKKYLNSIL | CACTCAGATGCAGTCTTCACTGACAACTATACCCGCCTTAGAAAACAAATGGCTGTAAAGAAATATTTGAACTCAATTCTG |
| VIP | Mouse | Mus Musculus | HSDAVFTDNYTRLRKQMAVKKYLNSIL | CACTCTGATGCCGTCTTCACAGATAACTACACCCGCCTCAGAAAGCAAATGGCTGTGAAGAAATACCTGAACTCCATCCTG |
| VIP | Chicken | Gallus Gallus | HSDAVFTDNYSRFRKQMAVKKYLNSVL | CACTCTGATGCTGTCTTCACTGACAACTACAGCCGCTTTCGAAAGCAAATGGCTGTGAAGAAATACTTAAACTCAGTTTTA |
| VIP | Lizard | Anolis Carolinensis | HSDAVFTDNYSRFRKQMAVKKYLNSVL | CACTCTGATGCTGTCTTCACTGACAATTACAGTCGCTTTCGAAAGCAGATGGCTGTGAAGAAGTATTTGAACTCTGTCTTA |
| VIP | Xenopus | Xenopus Tropicalis | HSDAVFTDNYSRFRKQMAVKKYLNSVL | CACTCTGATGCTGTGTTCACTGATAACTACAGCAGATTCAGGAAACAAATGGCTGTAAAGAAATACTTGAATTCTGTATTG |
| VIP | Zebrafish (a) | Danio Rerio | HSDAVFTDNYSRYRKQMAAKKYLNSVL | CACTCAGATGCCGTCTTTACGGACAACTACAGCCGCTACCGCAAGCAGATGGCGGCCAAGAAATACCTCAACTCAGTTCTG |
| VIP | Zebrafish (b) | Danio Rerio | HSDAIFTDNYSRFRKQMAVKKYLNSVL | CATTCAGATGCAATATTCACAGACAACTACAGTCGCTTTCGCAAGCAGATGGCGGTGAAGAAATATCTGAACTCTGTTCTC |
| VIP | Medeka | Oryzias Latipes | HSDAIFTDNYSRFRKQMAVKKYLNSVL | CACTCAGACGCCATCTTCACAGACAACTACAGCCGCTTCCGCAAACAGATGGCAGTCAAGAAGTACCTGAACTCAGTCTTA |
| VIP | Takifugu (b) | Takifugu Rubripes | HSDAIFTDNYSRFRKQMAVKKYLNSVL | CACTCAGACGCCATCTTTACAGACAACTACAGCCGCTTCCGTAAACAGATGGCTGTCAAGAAATACCTGAACTCGGTCTTA |
| VIP | Pufferfish | Tetraodon Nigroviridis | HSDAIFTDNYSRFRKQMAVKKYLNSVL | CACTCGGACGCCATCTTCACAGACAACTACAGCCGCTTCCGCAAACAGATGGCCGTCAAAAAATACCTGAACTCAGTCTTA |
| VIP | Stickleback (b) | Gasterosteus Aculeatus | HSDAIFTDNYSRFRKQMAVKKYLNSVL | CACTCTGACGCCATCTTCACAGACAACTACAGCCGCTTCCGCAAACAGATGGCCGTGAAGAAGTACCTGAACTCGGTGTTG |
| VIP | Lamprey | Petromyzon Marinus | HSDGLFTDLYSRYRKQMAVKKYLSTVL | CACTCGGACGGACTATTCACCGACCTGTACAGCCGGTACCGCAAACAAATGGCCGTTAAAAAATACCTGTCCACCGTCCTA |
| VIP | Lamprey | Petromyzon Marinus | HSDAVFTDLFSRLRKQQAAEKYAKSIL | CACTCGGACGCGGTGTTCACGGACCTGTTCAGCCGCCTGCGCAAGCAGCAAGCGGCGGAGAAGTACGCCAAGTCCATTCTG |
| **Peptide** | **Species** | **Latin name** | **Sequence** | **SeqDNA** |
| PRP | Human | Homo Sapiens | VAHGILNEAYRKVLDQLSAGKHLQSLV | GTCGCCCACGGGATCCTTAACGAGGCCTACCGCAAAGTGCTGGACCAGCTGTCCGCCGGGAAGCACCTGCAGTCGCTCGTG |
| PRP | Mouse | Mus Musculus | VAHEILNEAYRKVLDQLSARKYLQSVV | GTCGCCCACGAAATCCTTAACGAAGCCTATCGAAAAGTCTTGGACCAGCTGTCCGCCAGGAAGTACCTGCAGTCGGTCGTG |
| PRP | Chicken | Gallus Gallus | HADGIFSKAYRKLLGQLSARKYLHSLM | CACGCCGATGGGATCTTCAGCAAAGCCTACAGGAAACTCCTGGGCCAGCTGTCCGCCAGGAAATACCTGCACTCCCTGATG |
| PRP | Lizard | Anolis Carolinensis | HADGIFNKAYRKVLGQLSARKYLHSLM | CACGCGGATGGGATATTTAATAAAGCCTACCGGAAAGTGCTGGGCCAGCTGTCGGCGAGGAAGTATCTGCATTCCCTGATG |
| PRP | Xenopus | Xenopus Tropicalis | HADELLNKVYRNVLGHLSARKYLHTLM | CATGCTGATGAACTATTAAACAAAGTCTATAGGAATGTGCTGGGGCATTTGTCTGCAAGAAAATACCTGCATACTCTGATG |
| PRP | Zebrafish (a) | Danio Rerio | HADGMFNKAYRKALGQLSARKYLHTLM | CACGCTGACGGGATGTTTAATAAAGCCTACAGGAAAGCGCTCGGCCAGTTATCCGCGAGGAAGTACCTGCATACACTGATG |
| PRP | Zebrafish (b) | Danio Rerio | HADGLLDRALRDILVQLSARKYLHSLM | CATGCAGATGGATTATTAGATAGAGCCTTGAGGGACATCCTGGTTCAGTTATCAGCACGAAAATATCTGCATTCTCTGATG |
| PRP | Medeka | Oryzias Latipes | HADGMFNKAYRKALGQLSARKYLHSLM | CATGCAGACGGCATGTTTAATAAAGCCTACAGGAAAGCGCTGGGTCAGTTATCAGCAAGGAAATATCTTCATTCTCTGATG |
| PRP | Takifugua | Takifugu Rubripes | HADGMFNKAYRKALGQLSARKYLHSLM | CACGCAGACGGCATGTTTAATAAAGCCTACAGGAAGGCGCTGGGTCAGTTATCAGCCAGGAAATATCTGCATTCTCTGATG |
| PRP | Takifugub | Takifugu Rubripes | HAEEELDRALREILGQLTARHYRHFLM | CATGCTGAGGAAGAATTAGATAGAGCCTTGAGGGAGATCCTGGGTCAGTTAACAGCGAGGCATTATCGGCATTTTCTGATG |
| PRP | Pufferfish | Tetraodon Nigroviridis | HADGMFNKAYRKALGQLSARKYLHSLM | CACGCCGACGGCATGTTTAATAAAGCCTACAGGAAGGCGCTGGGTCAGTTATCAGCCAGGAAATATCTGCATTCTCTGATG |
| PRP | Pufferfish | Tetraodon Nigroviridis | HAEGELDRALREILGQLTARHYRHFLM | CATGCTGAGGGAGAATTAGATAGAGCCTTGAGGGAGATCCTGGGTCAGTTAACAGCGAGACATTATCGGCATTTTCTGATG |
| PRP | Stickleback (a) | Gasterosteus Aculeatus | HADGMFNKAYRKALGQLSARKYLHSLM | CATGCAGACGGCATGTTTAATAAAGCCTACAGGAAAGCGCTGGGTCAGTTATCAGCCAGGAAGTATCTGCATTCTCTGATG |
| PRP | Stickleback (b) | Gasterosteus Aculeatus | HAEEELDRALREILGQLTARHYLHSLM | CATGCTGAGGAAGAATTAGATAGAGCCTTGAGGGAGATCCTGGGTCAGTTAACAGCGAGACATTATCTGCATTCTCTGATG |
| PRP | Chelyosoma 1 | Chelyosoma Productum | HSDGIFTKDYRKYLGQLRAQKFLQWLM | CACTCCGATGGGATATTCACAAAAGATTATCGGAAGTACCTCGGGCAACTGCGAGCTCAAAAATTCCTGCAATGGCTTATG |
| PRP | Chelyosoma 2 | Chelyosoma Productum | HSDGIFTSDYRRYLGQLSAQKFLQWLM | CACTCCGATGGGATATTCACAAGTGATTATCGGAGGTACCTCGGGCAACTGAGTGCTCAAAAATTCCTGCAATGGCTTATG |
| **Peptide** | **Species** | **Latin name** | **Sequence** | **SeqDNA** |
| PH | Human | Homo Sapiens | HADGVFTSDFSKLLGQLSAKKYLESLM | CATGCTGATGGAGTTTTCACCAGTGACTTCAGTAAACTCTTGGGTCAACTTTCTGCCAAAAAGTACCTTGAGTCTCTTATG |
| PH | Mouse | Mus Musculus | HADGVFTSDYSRLLGQISAKKYLESLI | CATGCTGATGGAGTTTTCACCAGCGATTACAGCAGACTTCTGGGTCAGATTTCTGCCAAAAAATACCTTGAGTCACTCATT |
| PH | Chicken | Gallus Gallus | HADGIFTSVHSHLLAKLSVKRYLHSLI | CATGCTGATGGAATTTTCACCAGTGTCCACAGCCATCTTTTGGCTAAACTTTCTGTGAAGAGATATCTGCATTCGCTTATT |
| PH | Lizard | Anolis Carolinensis | HADGLFTSGYSKLLGQLSARRYLESLI | CATGCTGATGGACTCTTCACAAGTGGCTACAGCAAACTTTTGGGTCAACTTTCTGCAAGAAGATATTTGGAATCACTTATT |
| PH | Xenopus | Xenopus Tropicalis | HADGLFTSGYSKLLGQLSARRYLESLI | CACGCTGATGGGCTCTTCACTAGTGGATACAGCAAGCTTTTGGGTCAGCTTTCTGCAAGAAGATATCTAGAGTCTTTGATT |
| PH | Zebrafish (a) | Danio Rerio | HADGLFTSGYSKLLGQLSARRYLESLI | CACGCAGACGGGCTCTTCACAAGCGGATACAGTAAACTTCTAGGACAATTATCTGCCAGACGGTACCTGGAGTCATTGATC |
| PH | Zebrafish (b) | Danio Rerio | HADGLFTSGYSKLLGQLSAKEYLESLL | CACGCTGATGGACTGTTCACGAGCGGATACAGCAAACTGCTCGGCCAGCTGTCTGCTAAAGAGTATCTGGAGTCTTTACTG |
| PH | Medeka | Oryzias Latipes | HADGLFTSGYSKLLGQLSARRYLESLI | CACGCAGACGGTTTGTTCACCAGCGGATACAGCAAACTCCTGGGACAGTTATCAGCGCGGAGGTACCTGGAGTCTCTGATC |
| PH | Takifugu (a) | Takifugu Rubripes | HADGLFTSGYSKLLGQLSARRYLESLI | CACGCAGATGGTTTGTTCACCAGCGGCTACAGCAAACTCCTGGGTCAGCTGTCAGCACGGAGATATCTGGAGTCTCTGATC |
| PH | Pufferfish | Tetraodon Nigroviridis | HADGLFTSGYSKLLGQLSARRYLESLI | CACGCAGATGGACTGTTCACCAGCGGCTACAGCAAACTCCTGGGTCAGCTGTCAGCGCGGAGGTACCTGGAGTCTCTGATC |
| PH | Stickleback (b) | Gasterosteus Aculeatus | HADGLFTSGYSKLLGQLSARRYLESLI | CACGCGGACGGGCTGTTCACCAGCGGCTACAGTAAGCTGCTCGGGCAGCTGTCCGCGCGCAGGTACCTGGAGTCTCTGATC |
| PH | Lamprey | Petromyzon Marinus | HADALFHNNYKKLLGQMSARRYFESLL | CACGCCGATGCGCTCTTCCACAACAACTACAAGAAGCTGCTGGGACAGATGTCTGCCCGGCGCTACTTCGAGTCCCTGCTG |
| **Peptide** | **Species** | **Latin name** | **Sequence** | **SeqDNA** |
| GHRH | Human | Homo Sapiens | YADAIFTNSYRKVLGQLSARKLLQDIM | TATGCAGATGCCATCTTCACCAACAGCTACCGGAAGGTGCTGGGCCAGCTGTCCGCCCGCAAGCTGCTCCAGGACATCATG |
| GHRH | Mouse | Mus Musculus | HVDAIFTTNYRKLLSQLYARKVIQDIM | CACGTAGATGCCATCTTCACCACCAACTACAGGAAACTCCTGAGCCAGCTGTATGCCCGGAAAGTGATCCAGGACATCATG |
| GHRH | Chicken | Gallus Gallus | HADAIFTDNYRKFLGQISARKFLQTII | CACGCTGATGCCATTTTCACCGACAACTACCGGAAATTCCTGGGGCAGATTTCTGCCCGCAAATTCTTACAGACCATCATT |
| GHRH | Lizard | Anolis Carolinensis | HADAIFTDSYRKVRGKLSAQKLLQGIV | CATGCAGATGCCATATTCACCGACAGTTACCGTAAAGTCCGAGGCAAGCTGTCTGCCCAGAAGTTATTGCAGGGTATTGTT |
| GHRH | Xenopus | Xenopus Tropicalis | HVDAIFTNTYRKFLGQISARRYLQNMI | CATGTGGATGCCATATTCACTAACACATATAGGAAATTCCTGGGGCAGATTTCAGCCAGGAGGTACCTGCAGAACATGATA |
| GHRH | Zebrafish | Danio Rerio | HADAIFTNSYRKVLGQISARKFLQTVM | CATGCTGATGCCATCTTTACCAACAGCTACAGAAAAGTTCTTGGTCAAATATCTGCCAGAAAATTTCTTCAAACTGTTATG |
| GHRH | Takifugu | Takifugu Rubripes | HADAIFTNSYRKVLGQISARKILQTIM | CACGCCGATGCCATCTTTACAAACAGCTACAGAAAGGTCCTGGGTCAAATCTCAGCCAGGAAGATTCTACAGACCATTATG |
| GHRH | Pufferfish | Tetraodon Nigroviridis | HADAIFTNSYRKVLGQISARKILQTIM | CACGCCGATGCCATCTTTACAAACAGTTACAGGAAAGTCTTGGGTCAAATCTCAGCCAGGAAGATTCTACAGACCATTATG |
| GHRH | Stickleback | Pungitius Pungitius | HADAIFTNSYRKVLGQISARKFLQTIM | CACGCCGATGCCATCTTCACCAACAGTTACAGGAAAGTGCTGGGCCAAATCTCTGCCAGGAAGTTCCTTCAGACCATCATG |
| **Peptide** | **Species** | **Latin name** | **Sequence** | **SeqDNA** |
| SCT | Human | Homo sapiens | HSDGTFTSELSRLREGARLQRLLQGLV | CACTCAGACGGGACGTTCACCAGCGAGCTCAGCCGCCTGCGGGAGGGCGCGCGGCTCCAGCGGCTGCTACAGGGCCTGGTG |
| SCT | Mouse | Mus musculus | HSDGMFTSELSRLQDSARLQRLLQGLV | CACTCAGACGGAATGTTCACCAGCGAGCTCAGCCGCTTGCAGGACAGTGCCAGGCTGCAGCGCCTGCTGCAGGGTCTGGTG |
| SCT | Chicken | Gallus Gallus | HSDGLFTSEYSKMRGNAQVQKFIQNLM | CACTCGGATGGACTGTTCACCAGTGAATACAGCAAGATGAGAGGAAACGCTCAGGTGCAGAAGTTTATCCAAAATCTCATG |
| SCT | Chicken | Gallus Gallus | HADGLFHSELSKMNDNAYVQQLVKHLV | CATGCTGATGGGCTTTTTCACAGTGAGCTCAGCAAGATGAATGACAATGCTTACGTGCAGCAGCTGGTGAAACACCTGGTG |
| SCT | Xenopus | Xenopus Laevis | HVDGRFTSEFSRARGSAAIRKIINSAL | CACGTTGATGGGAGGTTCACCAGTGAATTCAGCCGAGCCAGGGGATCAGCTGCTATACGCAAGATCATCAACTCTGCTCTT |

**Supplementary Table 2. The 27-residue bioactive segments and their corresponding coding sequences from the PACAP peptide family were used to sequence and genomic analyses in this work.** The extended editable version of this table can be found in source data. Sequences and corresponding accession numbers were retrieved and grouped into families based on the classification reported by Cardoso et al.[5]

| Peptide | Species | Latin name | Sequence | SeqDNA |
|---|---|---|---|---|
| GCG | Human | Homo Sapiens | HSQGTFTSDYSKYLDSRRAQDFVQWLM | CATTCACAGGGCACATTCACCAGTGACTACAGCAAGTATCTGGACTCCAGGCGTGCCCAAGATTTTGTGCAGTGGTTGATG |
| GCG | Mouse | Mus Musculus | HSQGTFTSDYSKYLDSRRAQDFVQWLM | CACTCACAGGGCACATTCACCAGCGACTACAGCAAATACCTGGACTCCCGCCGTGCCCAAGATTTTGTGCAGTGGTTGATG |
| GCG | Chicken | Gallus Gallus | HSQGTFTSDYSKYLDSRRAQDFVQWLM | CATTCACAAGGCACATTCACCAGTGACTACAGCAAGTACTTGGACAGCAGACGAGCTCAGGATTTTGTGCAATGGTTAATG |
| GCG | Lizard | Anolis Carolinensis | HSQGTFTSDYSKYLDTRRAQDFVQWLM | CACTCACAAGGTACATTCACCAGCGATTACAGCAAGTACTTGGACACGAGGCGAGCTCAAGATTTTGTGCAGTGGTTAATG |
| GCG | Xenopus | Xenopus Tropicalis | HSQGTFTSDYSKYLDSRRAQDFIQWLM | CACTCACAAGGCACATTTACTAGTGATTACAGCAAGTACCTAGACTCCAGGCGGGCTCAAGACTTTATCCAGTGGTTAATG |
| GCG | Zebrafish (a) | Danio Rerio | HSEGTFSNDYSKYLETRRAQDFVQWLM | CATTCAGAGGGAACTTTCTCCAACGACTACAGCAAATACCTGGAGACCAGAAGAGCACAAGACTTCGTTCAGTGGCTCATG |
| GCG | Zebrafish (b) | Danio Rerio | HSEGTFSNDYSKYLETRRAQDFVQWLM | CATTCAGAGGGAACTTTCTCCAACGACTACAGCAAATACCTGGAGACCAGAAGAGCACAAGACTTCGTTCAGTGGCTCATG |
| GCG | Zebrafish (c) | Danio Rerio | HSEGTFSNDYSKYLETRRAQDFVQWLM | CATTCAGAAGGCACTTTCTCAAATGACTACAGCAAATATCTGGAGACCAGGAGAGCACAAGACTTTGTTCAGTGGTTAATG |
| GCG | Medeka | Oryzias Latipes | HSEGTFSNDYSKYLEDRKAQDFVRWLM | CACTCGGAGGGAACCTTCTCAAACGACTACAGCAAATACCTGGAGGACAGGAAGGCGCAGGACTTCGTTCGATGGCTGATG |
| GCG | Takifugu (a) | Takifugu Rubripes | HSEGTFSNDYSKYLEDRKAQDFVRWLM | CACTCGGAGGGGACTTTCTCCAACGACTACAGCAAATATCTGGAGGACAGGAAGGCGCAGGACTTTGTCCGATGGCTGATG |
| GCG | Takifugu (b) | Takifugu Rubripes | HSEGTFSNDYSKYLETRRAQDFVQWLK | CATTCAGAGGGGACCTTTTCCAACGACTACAGTAAATACCTCGAGACGAGGAGGGCGCAGGACTTTGTCCAGTGGCTGAAG |
| GCG | Pufferfish (a) | Tetraodon Nigroviridis | HSEGTFSNDYSKYLEDRKAQDFVRWLM | CACTCCGAGGGCACGTTCTCAAACGACTACAGCAAATATCTGGAGGACAGGAAGGCGCAGGACTTTGTCCGATGGCTGATG |
| GCG | Pufferfish (b) | Tetraodon Nigroviridis | HSEGTFSNDYSKYLETRRAQDFVQWLK | CATTCAGAGGGGACTTTTTCCAACGACTACAGCAAATATCTTGAGACGAGGAGGGCGCAAGACTTTGTCCAGTGGCTAAAA |
| GCG | Stickleback (a) | Gasterosteus Aculeatus | HSEGTFSNDYSKFLEEKKAQDFVQWLM | CATTCGGAGGGGACCTTCTCCAACGACTACAGCAAATTCCTGGAGGAAAAGAAGGCGCAGGACTTTGTTCAGTGGCTGATG |
| GCG | Stickleback (b) | Gasterosteus Aculeatus | HSEGTFSNDYSKYLETRRAQDFVQWLK | CATTCGGAGGGGACGTTTTCCAACGACTACAGTAAATACCTGGAGACGAGAAGAGCACAAGATTTTGTCCAGTGGCTAAAG |
| GCG | LampreyI (a) | Petromyzon Marinus | HSEGTFTSDYSKYLENKQAKDFVRWLM | CACTCGGAGGGAACCTTCACAAGCGACTACAGCAAGTACCTGGAGAACAAGCAGGCCAAGGACTTTGTACGCTGGCTCATG |
| GCG | Lamprey (b) | Petromyzon Marinus | HSQGSFTSDYSKHLDVKQAKDFVTWLL | CACTCACAGGGCTCCTTCACGAGCGACTACAGCAAACACCTGGATGTGAAGCAGGCCAAGGACTTTGTGACCTGGCTCTTG |
| **Peptide** | **Species** | **Latin name** | **Sequence** | **SeqDNA** |
| GLP-1 | Human | Homo Sapiens | HAEGTFTSDVSSYLEGQAAKEFIAWLV | CATGCTGAAGGGACCTTTACCAGTGATGTAAGTTCTTATTTGGAAGGCCAAGCTGCCAAGGAATTCATTGCTTGGCTGGTG |
| GLP-1 | Mouse | Mus Musculus | HAEGTFTSDVSSYLEGQAAKEFIAWLV | CATGCTGAAGGGACCTTTACCAGTGATGTGAGTTCTTACTTGGAGGGCCAGGCAGCAAAGGAATTCATTGCTTGGCTGGTG |
| GLP-1 | Chicken | Gallus Gallus | HAEGTYTSDITSYLEGQAAKEFIAWLV | CATGCTGAAGGCACCTACACCAGTGATATCACCTCTTATTTGGAAGGTCAAGCTGCCAAAGAATTCATTGCTTGGTTAGTG |
| GLP-1 | Lizard | Anolis Carolinensis | HADGTYTSDISSYLEGQAAKEFIAWLV | CATGCTGATGGCACCTATACCAGTGATATCAGCTCTTACTTGGAAGGTCAAGCTGCCAAGGAGTTCATTGCTTGGCTTGTG |
| GLP-1 | Xenopusa | Xenopus Tropicalis | HAEGTFTSDVTQHLDEKAAKEFIDWLI | CATGCAGAAGGTACATTCACCAGTGATGTGACTCAGCACTTGGATGAAAAAGCAGCCAAAGAATTCATTGACTGGCTAATA |
| GLP-1 | Xenopusb | Xenopus Tropicalis | HAEGTYTNDVTEYLEEKAAKEFIEWLI | CATGCCGAAGGTACATACACTAATGATGTAACTGAATATCTGGAGGAGAAAGCAGCCAAAGAATTTATTGAGTGGTTGATA |
| GLP-1 | Xenopusc | Xenopus Tropicalis | HAEGTFTNDMTNYLEEKAAKEFVGWLI | CACGCAGAAGGAACATTCACTAATGACATGACCAACTACCTGGAAGAGAAGGCAGCCAAAGAATTTGTGGGCTGGCTAATA |
| GLP-1 | Zebrafish (a) | Danio Rerio | HADGTYTSDVSSYLQDQAAKEFVSWLK | CATGCAGACGGCACATACACCAGTGATGTCAGCTCGTACCTGCAGGACCAGGCAGCAAAAGAGTTTGTGTCCTGGCTAAAA |
| GLP-1 | Zebrafish (b) | Danio Rerio | HAEGTYTSDVSSYLQDQAAQSFVAWLK | CATGCAGAGGGCACCTACACCAGCGACGTGAGCTCATACCTGCAAGACCAGGCGGCCCAGAGCTTTGTGGCCTGGCTAAAA |
| GLP-1 | Medeka | Oryzias Latipes | HADGTFTSDVSAYLKEQAIKDFVAKLK | CATGCAGATGGTACCTTCACCAGCGACGTGAGCGCCTACCTCAAGGAGCAGGCAATCAAAGACTTTGTTGCCAAGCTCAAG |
| GLP-1 | Takifugu (a) | Takifugu Rubripes | HADGTFTSDVSSYLKDQAIKDFVARLK | CACGCTGACGGGACCTTCACCAGTGACGTGAGCTCCTACCTCAAGGACCAGGCAATCAAAGACTTTGTGGCCAGGCTCAAA |
| GLP-1 | Takifugu (b) | Takifugu Rubripes | HADGTYTSDVSTYLQDQAAKEFVSWLK | CACGCTGACGGCACCTACACCAGCGACGTCAGCACCTACCTGCAGGACCAGGCAGCCAAGGAGTTTGTGTCCTGGCTGAAG |
| GLP-1 | Pufferfish (a) | Tetraodon Nigroviridis | HADGTFTSDVSSYLKDQAIKDFVARLK | CACGCTGACGGGACCTTCACCAGTGACGTGAGCTCCTACCTCAAGGACCAGGCCATCAAAGACTTTGTGGCCAGGCTCAAA |
| GLP-1 | Pufferfis (b) | Tetraodon Nigroviridis | HADGTYTSDVSAYLQDQAAKEFVSWLK | CACGCAGACGGCACCTACACCAGCGATGTGAGCGCCTACCTGCAGGACCAGGCAGCCAAGGAGTTCGTGTCCTGGCTGAAG |
| GLP-1 | Stickleback (a) | Gasterosteus Aculeatus | HADGTFTSDVSSYLKEQAIKDFVDKLK | CACGCTGACGGGACCTTCACCAGCGACGTGAGCTCGTACCTAAAGGAGCAGGCTATCAAAGACTTTGTGGACAAGCTTAAG |
| GLP-1 | Stickleback (b) | Gasterosteus Aculeatus | HADGTYTSDVSSYLQDQAVKEFVSWLK | CACGCAGACGGCACCTACACCAGCGACGTGAGCTCGTACCTGCAGGACCAAGCGGTCAAGGAGTTTGTGTCTTGGTTGAAG |
| GLP-1 | Lamprey (a) | Petromyzon Marinus | HADGTFTNDMTSYLDAKAARDFVSWLA | CACCCTGACGGCACCTTCACGAACGACATGACCTCCTACCTGGACGCGAAGGCTGCCCGCGACTTTCTCAGCTGGCTCGCC |
| **Peptide** | **Species** | **Latin name** | **Sequence** | **SeqDNA** |
| GLP-2 | Human | Homo Sapiens | HADGSFSDEMNTILDNLAARDFINWLI | CATGCTGATGGTTCTTTCTCTGATGAGATGAACACCATTCTTGATAATCTTGCCGCCAGGGACTTTATAAACTGGTTGATT |
| GLP-2 | Mouse | Mus Musculus | HADGSFSDEMSTILDNLATRDFINWLI | CACGCTGATGGCTCCTTCTCTGACGAGATGAGCACCATTCTGGATAATCTTGCCACCAGGGACTTCATCAACTGGCTGATT |
| GLP-2 | Chicken | Gallus Gallus | HADGTFTSDINKILDDMAAKEFLKWLI | CATGCGGATGGCACCTTCACAAGTGATATCAACAAAATCCTGGATGATATGGCTGCCAAAGAGTTCCTAAAATGGCTGATT |
| GLP-2 | Xenopus | Xenopus Tropicalis | HADGSFTNDINKVLDIIAAQEFLDWVI | CATGCCGATGGCAGTTTTACCAATGATATCAACAAAGTCCTAGATATTATTGCTGCACAGGAATTCTTAGATTGGGTAATT |
| GLP-2 | Zebrafish | Danio Rerio | HVDGSFTSDVNKVLDSIAAKEYLQWVM | CATGTAGACGGCAGTTTCACCAGTGACGTCAACAAGGTGTTGGACAGCATCGCAGCTAAAGAGTATTTACAATGGGTGATG |
| GLP-2 | Medeka | Oryzias Latipes | HVDGSFTSDVNKVLDSMAAKEYLLWVM | CATGTAGACGGAAGCTTCACCAGCGACGTGAACAAAGTCCTGGACTCCATGGCGGCCAAGGAATATTTACTCTGGGTCATG |
| GLP-2 | Takifugu | Takifugu Rubripes | HVDGTFTSDVNKVLDSMAAKEYLLWVM | CATGTAGATGGAACCTTCACCAGCGACGTGAACAAGGTGCTGGACTCCATGGCGGCCAAAGAATATTTACTGTGGGTGATG |
| GLP-2 | Pufferfish | Tetraodon Nigroviridis | HIDGTFTSDVNKVLDSMAAKEYLLWVM | CACATAGATGGAACCTTCACCAGCGATGTGAACAAGGTGCTGGACTCCATGGCCGCCAAGGAATATTTACTGTGGGTGATG |
| GLP-2 | Stickleback | Gasterosteus Aculeatus | HVDGSFTSDVNKVLDSLAAKEYLLWVM | CATGTAGATGGCAGCTTCACCAGTGATGTGAACAAGGTGCTGGACTCCCTGGCTGCCAAGGAATATTTACTCTGGGTCATG |
| GLP-2 | Lamprey (b) | Petromyzon Marinus | HSDGSFTNDMNVMLDRMSAKNFLEWLK | CACTCGGACGGCAGCTTCACCAATGACATGAACCTGATGCTCGACAGAATGTCAGCCAAGAACTTCCTGGAGTGGCTCAAA |
| **Peptide** | **Species** | **Latin name** | **Sequence** | **SeqDNA** |
| GIP | Human | Homo Sapiens | YAEGTFISDYSIAMDKIHQQDFVNWLL | TACGCGGAAGGGACTTTCATCAGTGACTACAGTATTGCCATGGACAAGATTCACCAACAAGACTTTGTGAACTGGCTGCTG |
| GIP | Mouse | Mus Musculus | YAEGTFISDYSIAMDKIRQQDFVNWLL | TACGCAGAGGGGACTTTCATCAGTGATTACAGCATCGCCATGGACAAGATCCGACAACAAGACTTCGTGAACTGGCTGCTG |
| GIP | Chicken | Gallus Gallus | YSEATLASDYSRTMDNMLKKNFVEWLL | TACTCGGAGGCCACGCTGGCGAGCGATTACAGCCGCACGATGGACAACATGCTGAAGAAGAACTTCGTGGAGTGGCTGCTG |
| GIP | Lizard | Anolis Carolinensis | YSEGTLASDYSRTLDNMLKKNFVEWLL | TATTCAGAAGGAACCTTAGCGAGCGATTACAGCCGAACCTTGGACAACATGCTGAAGAAGAACTTTGTGGAATGGCTCTTG |
| GIP | Xenopus | Xenopus Tropicalis | YSEAILASDYSRSVDNMLKKNFVDWLL | TACTCGGAGGCCATTTTGGCCAGCGATTACAGCAGATCAGTGGATAACATGCTGAAAAAGAACTTTGTGGACTGGTTACTT |
| GIP | Zebrafish | Danio Rerio | YAESTIASDISKIVDSMVQKNFVNFLL | TATGCTGAATCAACTATTGCCAGCGACATTAGCAAAATCGTGGACTCGATGGTTCAGAAAAACTTTGTGAACTTTCTGCTG |

**Supplementary Table 2 (continued). The 27-residue bioactive segments and their corresponding coding sequences from the glucagon peptide family were used to sequence and genomic analyses in this work.** The selected species broadly represent major vertebrate lineages, including cyclostomes (*Petromyzon marinus*), teleost fishes (*Danio rerio, Oryzias latipes, Takifugu rubripes, Tetraodon nigroviridis, Gasterosteus aculeatus*, and *Pungitius pungitius*), amphibians (*Xenopus* species), reptiles (*Anolis carolinensis*), birds (*Gallus gallus*), and mammals (*Mus musculus* and *Homo sapiens*), with the tunicate *Chelyosoma productum* included as a protochordate outgroup. The extended editable version of this table can be found in source data. Sequences and corresponding accession numbers were retrieved and grouped into families based on the classification reported by Cardoso et al.

| Peptide / Latin name | Best match / Similarity % DNA (same family, different species) | Best match / Similarity % Amino acid (same family, different species) | Best match / Similarity % DNA (other family) | Best match / Similarity % Amino acid (other family) |
|---|---|---|---|---|
| PACAP/Homo Sapiens | PACAP/Mus Musculus / 93.83% | PACAP/Mus Musculus / 100.0% | VIP/Takifugu Rubripes / 80.25% | VIP/Petromyzon Marinus / 85.19% |
| PACAP/Mus Musculus | PACAP/Homo Sapiens / 93.83% | PACAP/Homo Sapiens / 100.0% | VIP/Takifugu Rubripes / 80.25% | VIP/Petromyzon Marinus / 85.19% |
| PACAP/Gallus Gallus | PACAP/Homo Sapiens / 92.59% | PACAP/Homo Sapiens / 96.3% | VIP/Takifugu Rubripes / 77.78% | VIP/Petromyzon Marinus / 81.48% |
| PACAP/Anolis Carolinensis | PACAP/Mus Musculus / 91.36% | PACAP/Homo Sapiens / 100.0% | VIP/Oryzias Latipes / 80.25% | VIP/Petromyzon Marinus / 85.19% |
| PACAP/Xenopus Tropicalis | PACAP/Anolis Carolinensis / 91.36% | PACAP/Homo Sapiens / 100.0% | VIP/Homo Sapiens / 75.31% | VIP/Petromyzon Marinus / 85.19% |
| PACAP/Danio Rerio | PACAP/Gasterosteus Aculeatus / 86.42% | PACAP/Homo Sapiens / 92.59% | VIP/Petromyzon Marinus / 77.78% | VIP/Petromyzon Marinus / 88.89% |
| PACAP/Danio Rerio | PACAP/Xenopus Tropicalis / 85.19% | PACAP/Homo Sapiens / 96.3% | VIP/Anolis Carolinensis / 75.31% | VIP/Petromyzon Marinus / 85.19% |
| PACAP/Oryzias Latipes | PACAP/Gasterosteus Aculeatus / 91.36% | PACAP/Homo Sapiens / 100.0% | VIP/Mus Musculus / 77.78% | VIP/Petromyzon Marinus / 85.19% |
| PACAP/Takifugu Rubripes | PACAP/Tetraodon Nigroviridis / 93.83% | PACAP/Homo Sapiens / 100.0% | VIP/Oryzias Latipes / 81.48% | VIP/Petromyzon Marinus / 85.19% |
| PACAP/Takifugu Rubripes | PACAP/Tetraodon Nigroviridis / 98.77% | PACAP/Tetraodon Nigroviridis / 100.0% | VIP/Mus Musculus / 76.54% | VIP/Petromyzon Marinus / 81.48% |
| PACAP/Tetraodon Nigroviridis | PACAP/Takifugu Rubripes / 98.77% | PACAP/Takifugu Rubripes / 100.0% | VIP/Danio Rerio / 76.54% | VIP/Petromyzon Marinus / 81.48% |
| PACAP/Tetraodon Nigroviridis | PACAP/Gasterosteus Aculeatus / 95.06% | PACAP/Homo Sapiens / 100.0% | VIP/Oryzias Latipes / 80.25% | VIP/Petromyzon Marinus / 85.19% |
| PACAP/Gasterosteus Aculeatus | PACAP/Tetraodon Nigroviridis / 95.06% | PACAP/Homo Sapiens / 100.0% | VIP/Oryzias Latipes / 80.25% | VIP/Petromyzon Marinus / 85.19% |
| PACAP/Gasterosteus Aculeatus | PACAP/Takifugu Rubripes / 92.59% | PACAP/Takifugu Rubripes / 100.0% | VIP/Danio Rerio / 77.78% | VIP/Petromyzon Marinus / 81.48% |
| PACAP/Chelyosoma Productum | PACAP/Tetraodon Nigroviridis / 91.36% | PACAP/Homo Sapiens / 96.3% | VIP/Danio Rerio / 75.31% | VIP/Petromyzon Marinus / 81.48% |
| PACAP/Chelyosoma Productum | PACAP/Homo Sapiens / 83.95% | PACAP/Homo Sapiens / 85.19% | VIP/Gallus Gallus / 77.78% | VIP/Danio Rerio / 74.07% |
| VIP/Homo Sapiens | VIP/Mus Musculus / 85.19% | VIP/Mus Musculus / 100.0% | PACAP/Anolis Carolinensis / 75.31% | PACAP/Danio Rerio / 74.07% |
| VIP/Mus Musculus | VIP/Homo Sapiens / 85.19% | VIP/Homo Sapiens / 100.0% | PACAP/Oryzias Latipes / 77.78% | PACAP/Danio Rerio / 74.07% |
| VIP/Gallus Gallus | VIP/Anolis Carolinensis / 90.12% | VIP/Anolis Carolinensis / 100.0% | PACAP/Chelyosoma Productum / 77.78% | PACAP/Danio Rerio / 81.48% |
| VIP/Anolis Carolinensis | VIP/Gallus Gallus / 90.12% | VIP/Gallus Gallus / 100.0% | PACAP/Danio Rerio / 75.31% | PACAP/Danio Rerio / 81.48% |
| VIP/Xenopus Tropicalis | VIP/Gallus Gallus / 82.72% | VIP/Gallus Gallus / 100.0% | PACAP/Homo Sapiens / 72.84% | PACAP/Danio Rerio / 81.48% |
| VIP/Danio Rerio | VIP/Oryzias Latipes / 83.95% | VIP/Gallus Gallus / 92.59% | PACAP/Takifugu Rubripes / 79.01% | PACAP/Danio Rerio / 81.48% |
| VIP/Danio Rerio | VIP/Anolis Carolinensis / 82.72% | VIP/Oryzias Latipes / 100.0% | PACAP/Gasterosteus Aculeatus / 77.78% | PACAP/Homo Sapiens / 81.48% |
| VIP/Oryzias Latipes | VIP/Tetraodon Nigroviridis / 95.06% | VIP/Danio Rerio / 100.0% | PACAP/Takifugu Rubripes / 81.48% | PACAP/Homo Sapiens / 81.48% |
| VIP/Takifugu Rubripes | VIP/Oryzias Latipes / 93.83% | VIP/Danio Rerio / 100.0% | PACAP/Homo Sapiens / 80.25% | PACAP/Homo Sapiens / 81.48% |
| VIP/Tetraodon Nigroviridis | VIP/Oryzias Latipes / 95.06% | VIP/Danio Rerio / 100.0% | PACAP/Homo Sapiens / 80.25% | PACAP/Homo Sapiens / 81.48% |
| VIP/Gasterosteus Aculeatus | VIP/Oryzias Latipes / 92.59% | VIP/Danio Rerio / 100.0% | PACAP/Mus Musculus / 77.78% | PACAP/Homo Sapiens / 81.48% |
| VIP/Petromyzon Marinus | VIP/Tetraodon Nigroviridis / 79.01% | VIP/Gallus Gallus / 77.78% | PACAP/Homo Sapiens / 80.25% | PACAP/Danio Rerio / 88.89% |
| VIP/Petromyzon Marinus | VIP/Danio Rerio / 71.6% | VIP/Homo Sapiens / 70.37% | PACAP/Homo Sapiens / 62.96% | PACAP/Danio Rerio / 59.26% |
| PRP/Homo Sapiens | PRP/Mus Musculus / 87.65% | PRP/Mus Musculus / 85.19% | GHRH/Homo Sapiens / 65.43% | GHRH/Homo Sapiens / 51.85% |
| PRP/Mus Musculus | PRP/Homo Sapiens / 87.65% | PRP/Homo Sapiens / 85.19% | GHRH/Pungitius Pungitius / 62.96% | GHRH/Homo Sapiens / 55.56% |
| PRP/Gallus Gallus | PRP/Anolis Carolinensis / 83.95% | PRP/Anolis Carolinensis / 92.59% | PH/Takifugu Rubripes / 76.54% | PH/Homo Sapiens / 70.37% |
| PRP/Anolis Carolinensis | PRP/Danio Rerio / 85.19% | PRP/Gallus Gallus / 92.59% | PH/Tetraodon Nigroviridis / 71.6% | PH/Homo Sapiens / 66.67% |
| PRP/Xenopus Tropicalis | PRP/Oryzias Latipes / 75.31% | PRP/Anolis Carolinensis / 74.07% | GHRH/Danio Rerio / 69.14% | GHRH/Danio Rerio / 55.56% |
| PRP/Danio Rerio | PRP/Gasterosteus Aculeatus / 87.65% | PRP/Oryzias Latipes / 96.3% | PH/Gasterosteus Aculeatus / 71.6% | PH/Homo Sapiens / 62.96% |
| PRP/Danio Rerio | PRP/Gasterosteus Aculeatus / 85.19% | PRP/Gasterosteus Aculeatus / 74.07% | PH/Takifugu Rubripes / 71.6% | PH/Homo Sapiens / 55.56% |
| PRP/Oryzias Latipes | PRP/Gasterosteus Aculeatus / 96.3% | PRP/Takifugu Rubripes / 100.0% | PH/Takifugu Rubripes / 72.84% | PH/Homo Sapiens / 66.67% |
| PRP/Takifugu Rubripes | PRP/Tetraodon Nigroviridis / 98.77% | PRP/Oryzias Latipes / 100.0% | PH/Takifugu Rubripes / 72.84% | PH/Homo Sapiens / 66.67% |
| PRP/Takifugu Rubripes | PRP/Tetraodon Nigroviridis / 97.53% | PRP/Tetraodon Nigroviridis / 96.3% | PH/Tetraodon Nigroviridis / 60.49% | PH/Homo Sapiens / 37.04% |
| PRP/Tetraodon Nigroviridis | PRP/Takifugu Rubripes / 98.77% | PRP/Oryzias Latipes / 100.0% | PH/Takifugu Rubripes / 71.6% | PH/Homo Sapiens / 66.67% |
| PRP/Tetraodon Nigroviridis | PRP/Takifugu Rubripes / 97.53% | PRP/Takifugu Rubripes / 96.3% | PH/Homo Sapiens / 61.73% | PH/Homo Sapiens / 40.74% |
| PRP/Gasterosteus Aculeatus | PRP/Oryzias Latipes / 96.3% | PRP/Oryzias Latipes / 100.0% | PH/Oryzias Latipes / 72.84% | PH/Homo Sapiens / 66.67% |
| PRP/Gasterosteus Aculeatus | PRP/Takifugu Rubripes / 96.3% | PRP/Takifugu Rubripes / 92.59% | PH/Homo Sapiens / 62.96% | PH/Homo Sapiens / 44.44% |
| PRP/Chelyosoma Productum | PRP/Gallus Gallus / 67.9% | PRP/Gallus Gallus / 66.67% | GCG/Anolis Carolinensis / 66.67% | GCG/Homo Sapiens / 66.67% |
| PRP/Chelyosoma Productum | PRP/Gallus Gallus / 64.2% | PRP/Gallus Gallus / 62.96% | GCG/Anolis Carolinensis / 64.2% | PH/Homo Sapiens / 62.96% |
| PH/Homo Sapiens | PH/Mus Musculus / 83.95% | PH/Mus Musculus / 85.19% | PRP/Gallus Gallus / 70.37% | PRP/Gallus Gallus / 70.37% |
| PH/Mus Musculus | PH/Homo Sapiens / 83.95% | PH/Homo Sapiens / 85.19% | GHRH/Gallus Gallus / 70.37% | PRP/Gallus Gallus / 62.96% |
| PH/Gallus Gallus | PH/Anolis Carolinensis / 79.01% | PH/Anolis Carolinensis / 66.67% | PRP/Gallus Gallus / 64.2% | PRP/Gallus Gallus / 55.56% |
| PH/Anolis Carolinensis | PH/Xenopus Tropicalis / 85.19% | PH/Xenopus Tropicalis / 100.0% | GHRH/Danio Rerio / 70.37% | PRP/Gallus Gallus / 70.37% |
| PH/Xenopus Tropicalis | PH/Anolis Carolinensis / 85.19% | PH/Anolis Carolinensis / 100.0% | PRP/Gallus Gallus / 69.14% | PRP/Gallus Gallus / 70.37% |
| PH/Danio Rerio | PH/Oryzias Latipes / 81.48% | PH/Anolis Carolinensis / 100.0% | PRP/Gallus Gallus / 67.9% | PRP/Gallus Gallus / 70.37% |
| PH/Danio Rerio | PH/Tetraodon Nigroviridis / 77.78% | PH/Anolis Carolinensis / 88.89% | PRP/Gallus Gallus / 67.9% | PRP/Gallus Gallus / 66.67% |
| PH/Oryzias Latipes | PH/Tetraodon Nigroviridis / 91.36% | PH/Anolis Carolinensis / 100.0% | PRP/Gasterosteus Aculeatus / 72.84% | PRP/Gallus Gallus / 70.37% |
| PH/Takifugu Rubripes | PH/Tetraodon Nigroviridis / 93.83% | PH/Anolis Carolinensis / 100.0% | PRP/Gallus Gallus / 76.54% | PRP/Gallus Gallus / 70.37% |
| PH/Tetraodon Nigroviridis | PH/Takifugu Rubripes / 93.83% | PH/Anolis Carolinensis / 100.0% | PRP/Gallus Gallus / 76.54% | PRP/Gallus Gallus / 70.37% |
| PH/Gasterosteus Aculeatus | PH/Tetraodon Nigroviridis / 87.65% | PH/Anolis Carolinensis / 100.0% | PRP/Gallus Gallus / 72.84% | PRP/Gallus Gallus / 70.37% |
| PH/Petromyzon Marinus | PH/Tetraodon Nigroviridis / 70.37% | PH/Anolis Carolinensis / 70.37% | PRP/Gallus Gallus / 72.84% | PRP/Gallus Gallus / 59.26% |
| GHRH/Homo Sapiens | GHRH/Mus Musculus / 80.25% | GHRH/Takifugu Rubripes / 85.19% | PRP/Gallus Gallus / 69.14% | PRP/Anolis Carolinensis / 66.67% |
| GHRH/Mus Musculus | GHRH/Homo Sapiens / 80.25% | GHRH/Homo Sapiens / 66.67% | PRP/Gallus Gallus / 71.6% | PRP/Gallus Gallus / 55.56% |
| GHRH/Gallus Gallus | GHRH/Pungitius Pungitius / 79.01% | GHRH/Pungitius Pungitius / 85.19% | PH/Mus Musculus / 70.37% | PRP/Gallus Gallus / 59.26% |
| GHRH/Anolis Carolinensis | GHRH/Homo Sapiens / 72.84% | GHRH/Homo Sapiens / 74.07% | PH/Tetraodon Nigroviridis / 62.96% | PRP/Anolis Carolinensis / 55.56% |
| GHRH/Xenopus Tropicalis | GHRH/Pungitius Pungitius / 75.31% | GHRH/Gallus Gallus / 74.07% | PRP/Gallus Gallus / 70.37% | PH/Anolis Carolinensis / 59.26% |
| GHRH/Danio Rerio | GHRH/Takifugu Rubripes / 81.48% | GHRH/Pungitius Pungitius / 96.3% | PRP/Oryzias Latipes / 71.6% | PRP/Anolis Carolinensis / 66.67% |
| GHRH/Takifugu Rubripes | GHRH/Tetraodon Nigroviridis / 95.06% | GHRH/Tetraodon Nigroviridis / 100.0% | PRP/Tetraodon Nigroviridis / 71.6% | PRP/Anolis Carolinensis / 66.67% |
| GHRH/Tetraodon Nigroviridis | GHRH/Takifugu Rubripes / 95.06% | GHRH/Takifugu Rubripes / 100.0% | PRP/Gallus Gallus / 69.14% | PRP/Anolis Carolinensis / 66.67% |
| GHRH/Pungitius Pungitius | GHRH/Tetraodon Nigroviridis / 87.65% | GHRH/Danio Rerio / 96.3% | PRP/Gallus Gallus / 75.31% | PRP/Anolis Carolinensis / 66.67% |
| SCT/Homo Sapiens | SCT/Mus Musculus / 83.95% | SCT/Mus Musculus / 85.19% | GCG/Gasterosteus Aculeatus / 58.02% | PRP/Chelyosoma Productum / 44.44% |
| SCT/Mus Musculus | SCT/Homo Sapiens / 83.95% | SCT/Homo Sapiens / 85.19% | GCG/Mus Musculus / 56.79% | GCG/Homo Sapiens / 48.15% |
| SCT/Gallus Gallus | SCT/Mus Musculus / 58.02% | SCT/Homo Sapiens / 51.85% | GCG/Petromyzon Marinus / 62.96% | PRP/Chelyosoma Productum / 55.56% |
| SCT/Gallus Gallus | SCT/Mus Musculus / 61.73% | SCT/Mus Musculus / 51.85% | GIP/Homo Sapiens / 54.32% | PH/Anolis Carolinensis / 37.04% |
| SCT/Xenopus Laevis | SCT/Gallus Gallus / 56.79% | SCT/Gallus Gallus / 48.15% | VIP/Xenopus Tropicalis / 55.56% | PACAP/Chelyosoma Productum / 48.15% |

**Supplementary Table 3. Nearest-neighbour sequence similarity analysis of PACAP family peptides at the nucleotide and amino acid levels. (For detailed description see the next page)**

| Peptide / Latin name | Best match / Similarity % DNA (same family, different species) | Best match / Similarity % Amino acid (same family, different species) | Best match / Similarity % DNA (other family) | Best match / Similarity % Amino acid (other family) |
|---|---|---|---|---|
| GCG/Homo Sapiens | GCG/Mus Musculus / 92.59% | GCG/Mus Musculus / 100.0% | GLP-2/Takifugu Rubripes / 69.14% | PRP/Chelyosoma Productum / 66.67% |
| GCG/Mus Musculus | GCG/Homo Sapiens / 92.59% | GCG/Homo Sapiens / 100.0% | GLP-1/Takifugu Rubripes / 66.67% | PRP/Chelyosoma Productum / 66.67% |
| GCG/Gallus Gallus | GCG/Anolis Carolinensis / 88.89% | GCG/Homo Sapiens / 100.0% | GLP-2/Danio Rerio / 70.37% | PRP/Chelyosoma Productum / 66.67% |
| GCG/Anolis Carolinensis | GCG/Gallus Gallus / 88.89% | GCG/Homo Sapiens / 96.3% | GIP/Mus Musculus / 67.9% | PRP/Chelyosoma Productum / 66.67% |
| GCG/Xenopus Tropicalis | GCG/Anolis Carolinensis / 85.19% | GCG/Homo Sapiens / 96.3% | PRP/Chelyosoma Productum / 64.2% | PRP/Chelyosoma Productum / 66.67% |
| GCG/Danio Rerio | GCG/Gasterosteus Aculeatus / 86.42% | GCG/Takifugu Rubripes / 96.3% | GIP/Mus Musculus / 66.67% | PRP/Chelyosoma Productum / 62.96% |
| GCG/Danio Rerio | GCG/Gasterosteus Aculeatus / 86.42% | GCG/Takifugu Rubripes / 96.3% | GIP/Mus Musculus / 66.67% | PRP/Chelyosoma Productum / 62.96% |
| GCG/Danio Rerio | GCG/Tetraodon Nigroviridis / 83.95% | GCG/Takifugu Rubripes / 96.3% | GLP-1/Xenopus Tropicalis / 65.43% | PRP/Chelyosoma Productum / 62.96% |
| GCG/Oryzias Latipes | GCG/Takifugu Rubripes / 92.59% | GCG/Takifugu Rubripes / 100.0% | GLP-1/Takifugu Rubripes / 66.67% | PRP/Chelyosoma Productum / 55.56% |
| GCG/Takifugu Rubripes | GCG/Tetraodon Nigroviridis / 95.06% | GCG/Oryzias Latipes / 100.0% | GLP-1/Takifugu Rubripes / 66.67% | PRP/Chelyosoma Productum / 55.56% |
| GCG/Takifugu Rubripes | GCG/Tetraodon Nigroviridis / 91.36% | GCG/Tetraodon Nigroviridis / 100.0% | GLP-1/Danio Rerio / 61.73% | PRP/Chelyosoma Productum / 59.26% |
| GCG/Tetraodon Nigroviridis | GCG/Takifugu Rubripes / 95.06% | GCG/Oryzias Latipes / 100.0% | GLP-1/Takifugu Rubripes / 66.67% | PRP/Chelyosoma Productum / 55.56% |
| GCG/Tetraodon Nigroviridis | GCG/Takifugu Rubripes / 91.36% | GCG/Takifugu Rubripes / 100.0% | GLP-1/Xenopus Tropicalis / 61.73% | PRP/Chelyosoma Productum / 59.26% |
| GCG/Gasterosteus Aculeatus | GCG/Oryzias Latipes / 88.89% | GCG/Danio Rerio / 85.19% | GLP-1/Takifugu Rubripes / 66.67% | GLP-1/Xenopus Tropicalis / 59.26% |
| GCG/Gasterosteus Aculeatus | GCG/Takifugu Rubripes / 88.89% | GCG/Takifugu Rubripes / 100.0% | GIP/Homo Sapiens / 61.73% | PRP/Chelyosoma Productum / 59.26% |
| GCG/Petromyzon Marinus | GCG/Oryzias Latipes / 85.19% | GCG/Oryzias Latipes / 77.78% | GLP-1/Takifugu Rubripes / 72.84% | GLP-1/Homo Sapiens / 62.96% |
| GCG/Petromyzon Marinus | GCG/Mus Musculus / 72.84% | GCG/Homo Sapiens / 70.37% | GLP-1/Takifugu Rubripes / 69.14% | GLP-1/Xenopus Tropicalis / 55.56% |
| GLP-1/Homo Sapiens | GLP-1/Mus Musculus / 92.59% | GLP-1/Mus Musculus / 100.0% | GLP-2/Gallus Gallus / 61.73% | GCG/Petromyzon Marinus / 62.96% |
| GLP-1/Mus Musculus | GLP-1/Homo Sapiens / 92.59% | GLP-1/Homo Sapiens / 100.0% | GLP-2/Gallus Gallus / 61.73% | GCG/Petromyzon Marinus / 62.96% |
| GLP-1/Gallus Gallus | GLP-1/Anolis Carolinensis / 90.12% | GLP-1/Anolis Carolinensis / 92.59% | GLP-2/Gallus Gallus / 66.67% | GLP-2/Gallus Gallus / 59.26% |
| GLP-1/Anolis Carolinensis | GLP-1/Gallus Gallus / 90.12% | GLP-1/Gallus Gallus / 92.59% | GLP-2/Gallus Gallus / 69.14% | GLP-2/Gallus Gallus / 62.96% |
| GLP-1/Xenopus Tropicalis | GLP-1/Mus Musculus / 74.07% | GLP-1/Homo Sapiens / 70.37% | GLP-2/Gallus Gallus / 69.14% | GLP-2/Gallus Gallus / 66.67% |
| GLP-1/Xenopus Tropicalis | GLP-1/Gallus Gallus / 71.6% | GLP-1/Gallus Gallus / 74.07% | GCG/Danio Rerio / 65.43% | GCG/Petromyzon Marinus / 55.56% |
| GLP-1/Xenopus Tropicalis | GLP-1/Danio Rerio / 72.84% | GLP-1/Petromyzon Marinus / 70.37% | GCG/Petromyzon Marinus / 70.37% | GCG/Petromyzon Marinus / 62.96% |
| GLP-1/Danio Rerio | GLP-1/Takifugu Rubripes / 86.42% | GLP-1/Takifugu Rubripes / 96.3% | GLP-2/Gallus Gallus / 65.43% | GLP-2/Gallus Gallus / 62.96% |
| GLP-1/Danio Rerio | GLP-1/Takifugu Rubripes / 81.48% | GLP-1/Takifugu Rubripes / 81.48% | GCG/Petromyzon Marinus / 66.67% | GCG/Homo Sapiens / 55.56% |
| GLP-1/Oryzias Latipes | GLP-1/Takifugu Rubripes / 87.65% | GLP-1/Gasterosteus Aculeatus / 92.59% | GCG/Petromyzon Marinus / 64.2% | GCG/Petromyzon Marinus / 55.56% |
| GLP-1/Takifugu Rubripes | GLP-1/Tetraodon Nigroviridis / 98.77% | GLP-1/Tetraodon Nigroviridis / 100.0% | GCG/Petromyzon Marinus / 65.43% | GCG/Petromyzon Marinus / 55.56% |
| GLP-1/Takifugu Rubripes | GLP-1/Tetraodon Nigroviridis / 93.83% | GLP-1/Danio Rerio / 96.3% | GCG/Petromyzon Marinus / 72.84% | GLP-2/Gallus Gallus / 62.96% |
| GLP-1/Tetraodon Nigroviridis | GLP-1/Takifugu Rubripes / 98.77% | GLP-1/Takifugu Rubripes / 100.0% | GCG/Petromyzon Marinus / 65.43% | GCG/Petromyzon Marinus / 55.56% |
| GLP-1/Tetraodon Nigroviridis | GLP-1/Takifugu Rubripes / 93.83% | GLP-1/Danio Rerio / 96.3% | GCG/Petromyzon Marinus / 67.9% | GLP-2/Gallus Gallus / 62.96% |
| GLP-1/Gasterosteus Aculeatus | GLP-1/Takifugu Rubripes / 88.89% | GLP-1/Oryzias Latipes / 92.59% | GCG/Petromyzon Marinus / 65.43% | GCG/Petromyzon Marinus / 55.56% |
| GLP-1/Gasterosteus Aculeatus | GLP-1/Takifugu Rubripes / 88.89% | GLP-1/Danio Rerio / 96.3% | GCG/Petromyzon Marinus / 67.9% | GLP-2/Gallus Gallus / 59.26% |
| GLP-1/Petromyzon Marinus | GLP-1/Xenopus Tropicalis / 71.6% | GLP-1/Xenopus Tropicalis / 70.37% | GCG/Petromyzon Marinus / 65.43% | GCG/Homo Sapiens / 55.56% |
| GLP-2/Homo Sapiens | GLP-2/Mus Musculus / 87.65% | GLP-2/Mus Musculus / 92.59% | GLP-1/Xenopus Tropicalis / 60.49% | GLP-1/Petromyzon Marinus / 55.56% |
| GLP-2/Mus Musculus | GLP-2/Homo Sapiens / 87.65% | GLP-2/Homo Sapiens / 92.59% | GLP-1/Takifugu Rubripes / 62.96% | GLP-1/Petromyzon Marinus / 51.85% |
| GLP-2/Gallus Gallus | GLP-2/Xenopus Tropicalis / 75.31% | GLP-2/Xenopus Tropicalis / 70.37% | GLP-1/Anolis Carolinensis / 69.14% | GLP-1/Xenopus Tropicalis / 66.67% |
| GLP-2/Xenopus Tropicalis | GLP-2/Gallus Gallus / 75.31% | GLP-2/Gallus Gallus / 70.37% | GLP-1/Anolis Carolinensis / 64.2% | GLP-1/Xenopus Tropicalis / 55.56% |
| GLP-2/Danio Rerio | GLP-2/Takifugu Rubripes / 81.48% | GLP-2/Oryzias Latipes / 92.59% | GCG/Gallus Gallus / 70.37% | PH/Homo Sapiens / 51.85% |
| GLP-2/Oryzias Latipes | GLP-2/Takifugu Rubripes / 91.36% | GLP-2/Takifugu Rubripes / 96.3% | GCG/Petromyzon Marinus / 69.14% | PH/Homo Sapiens / 51.85% |
| GLP-2/Takifugu Rubripes | GLP-2/Tetraodon Nigroviridis / 93.83% | GLP-2/Oryzias Latipes / 96.3% | GCG/Homo Sapiens / 69.14% | GLP-1/Xenopus Tropicalis / 55.56% |
| GLP-2/Tetraodon Nigroviridis | GLP-2/Takifugu Rubripes / 93.83% | GLP-2/Takifugu Rubripes / 96.3% | GCG/Petromyzon Marinus / 66.67% | GLP-1/Xenopus Tropicalis / 55.56% |
| GLP-2/Gasterosteus Aculeatus | GLP-2/Oryzias Latipes / 90.12% | GLP-2/Oryzias Latipes / 96.3% | GCG/Homo Sapiens / 65.43% | PH/Homo Sapiens / 55.56% |
| GLP-2/Petromyzon Marinus | GLP-2/Gallus Gallus / 65.43% | GLP-2/Gallus Gallus / 59.26% | GLP-1/Xenopus Tropicalis / 62.96% | PRP/Chelyosoma Productum / 51.85% |
| GIP/Homo Sapiens | GIP/Mus Musculus / 88.89% | GIP/Mus Musculus / 96.3% | GCG/Anolis Carolinensis / 64.2% | GCG/Homo Sapiens / 51.85% |
| GIP/Mus Musculus | GIP/Homo Sapiens / 88.89% | GIP/Homo Sapiens / 96.3% | GCG/Anolis Carolinensis / 67.9% | GCG/Homo Sapiens / 55.56% |
| GIP/Gallus Gallus | GIP/Anolis Carolinensis / 81.48% | GIP/Anolis Carolinensis / 92.59% | GCG/Petromyzon Marinus / 62.96% | GCG/Petromyzon Marinus / 48.15% |
| GIP/Anolis Carolinensis | GIP/Gallus Gallus / 81.48% | GIP/Gallus Gallus / 92.59% | GCG/Petromyzon Marinus / 67.9% | GCG/Petromyzon Marinus / 55.56% |
| GIP/Xenopus Tropicalis | GIP/Gallus Gallus / 80.25% | GIP/Gallus Gallus / 85.19% | GCG/Petromyzon Marinus / 65.43% | GCG/Petromyzon Marinus / 44.44% |
| GIP/Danio Rerio | GIP/Anolis Carolinensis / 64.2% | GIP/Gallus Gallus / 55.56% | GLP-2/Gallus Gallus / 58.02% | GLP-2/Gallus Gallus / 44.44% |

**Supplementary Table 3 (continued). Nearest-neighbour sequence similarity analysis of glucagon family peptides at the nucleotide and amino acid levels.** For each peptide, the closest matching sequence was identified separately at the DNA and amino acid levels based on pairwise percentage identity calculated by direct position-by-position comparison. Similarity was determined across the available coding sequence (DNA) and corresponding peptide sequence (amino acids). (S.Table 2) Within-family comparisons were restricted to sequences originating from different species, thereby excluding paralogous peptides derived from the same organism. For each peptide, the table reports the best match within the same family (different species) and the best match outside the family, together with the corresponding percentage identity.

1: CAC(H) –1→ CAT(H)
2: TCG(S) –1→ TCT(S) –1→ TCA(S) –2→ ATA(I)
3: GAC(D) –1→ GAT(D)
4: GGG(G) –1→ GGC(G)
5: ATC(I) –2→ GTT(V)
6: TTC(F) –1→ TTT(F)
7: ACG(T) –1→ ACA(T) –1→ ACC(T)
8: GAC(D) –1→ GAT(D)
9: AGC(S) –2→ ATT(I)
10: TAC(Y) –1→ TAT(Y)
11: AGC(S) –1→ AGT(S)
12: CGC(R)
13: TAC(Y) –1→ TAT(Y)
14: CGG(R) –1→ CGA(R) –1→ AGA(R)
15: AAA(K) –1→ AAG(K) –1→ AAT(N)
16: CAA(Q) –1→ CAG(Q)
17: ATG(M)
18: GCT(A) –1→ GCC(A) –1→ GCG(A) –1→ GCA(A)
19: GTC(V) –1→ GTG(V) –1→ GTT(V)
20: AAG(K) –1→ CAG(Q)
21: AAA(K) –1→ AAG(K)
22: TAC(Y) –1→ TAT(Y)
23: CTG(L) –1→ TTG(L) –1→ TTA(L) –1→ ATA(I)
24: GCG(A) –1→ GCA(A) –1→ GCC(A) –2→ AAC(N)
25: GCC(A) –1→ GCA(A) –1→ GCG(A) –1→ ACG(T)
26: GTC(V) –1→ GTT(V) –1→ GTG(V) –1→ CTG(L)
27: CTA(L) –1→ CTG(L) –1→ CTT(L)

**Supplementary Table 4/1. Position-specific mutational pathways in PACAP sequences.** For each residue position, the codons observed across the analysed species are connected by the minimal mutational routes based exclusively on the detected codons. Numbers above the arrows indicate the minimal number of nucleotide substitutions required for each transition. Amino acids are colored according to residue class: acidic (red), basic (blue), polar (light green), and hydrophobic (gray).

1: CAC(H) –1→ CAT(H)
2: TCA(S) –1→ TCT(S) –1→ TCG(S)
3: GAT(D) –1→ GAC(D)
4: GCC(A) –1→ GCT(A) –1→ GCG(A) –1→ GCA(A) –1→ GGA(G)
5: GTG(V) –1→ GTC(V) –1→ ATC(I) –1→ ATA(I) –1→ CTA(L)
6: TTC(F) –1→ TTT(F)
7: ACT(T) –1→ ACA(T) –1→ ACG(T) –1→ ACC(T)
8: GAC(D) –1→ GAT(D)
9: AAC(N) –1→ AAT(N) –3→ CTG(L)
10: TAT(Y) –1→ TAC(Y) –1→ TTC(F)
11: ACC(T) –1→ AGC(S) –1→ AGT(S)
12: CGC(R) –1→ CGG(R) –2→ AGA(R)
13: CTC(L) –1→ CTG(L) –1→ CTT(L) –1→ TTT(F) –1→ TTC(F) –1→ TAC(Y)
14: AGG(R) –1→ AGA(R) –1→ CGA(R) –1→ CGC(R) –1→ CGT(R)
15: AAA(K) –1→ AAG(K)
16: CAA(Q) –1→ CAG(Q)
17: ATG(M) –3→ CAA(Q)
18: GCT(A) –1→ GCG(A) –1→ GCA(A) –1→ GCC(A)
19: GTA(V) –1→ GTG(V) –1→ GTT(V) –1→ GTC(V) –1→ GCC(A) –1→ GCG(A)
20: AAA(K) –1→ AAG(K) –1→ GAG(E)
21: AAA(K) –1→ AAG(K)
22: TAT(Y) –1→ TAC(Y)
23: TTA(L) –1→ TTG(L) –1→ CTG(L) –1→ CTC(L) –2→ GCC(A)
24: AAT(N) –1→ AAG(K) –1→ AAC(N) –2→ TCC(S)
25: TCA(S) –1→ TCT(S) –1→ TCG(S) –1→ TCC(S) –1→ ACC(T)
26: ATT(I) –1→ ATC(I) –1→ GTC(V) –1→ GTT(V) –1→ GTA(V) –1→ GTG(V)
27: CTG(L) –1→ TTG(L) –1→ TTA(L) –1→ CTA(L) –1→ CTC(L)

**Supplementary Table 4/2. Position-specific mutational pathways in VIP sequences.** For each residue position, the codons observed across the analysed species are connected by the minimal mutational routes based exclusively on the detected codons. Numbers above the arrows indicate the minimal number of nucleotide substitutions required for each transition. Amino acids are colored according to residue class: acidic (red), basic (blue), polar (light green), and hydrophobic (gray).

1: GTC(V) –2→ CAC(H) –1→ CAT(H)
2: GCG(A) –1→ GCT(A) –1→ GCA(A) –1→ GCC(A) –1→ TCC(S)
3: CAC(H) –1→ GAC(D) –1→ GAT(D) –1→ GAG(E)
4: GGG(G) –1→ GGC(G) –1→ GGA(G) –1→ GAA(E)
5: ATC(I) –1→ ATG(M) –1→ ATA(I) –1→ CTA(L) –1→ TTA(L) –2→ GAA(E)
6: CTT(L) –1→ TTT(F) –1→ TTC(F) –1→ TTA(L)
7: GAT(D) –1→ AAT(N) –1→ AAC(N) –1→ AGC(S) –2→ ACA(T)
8: GAG(E) –1→ GAA(E) –1→ AAA(K) –1→ AGA(R) –1→ AGT(S)
9: GCC(A) –1→ GTC(V) –2→ GAT(D)
10: TAC(Y) –1→ TAT(Y) –2→ TTG(L)
11: CGC(R) –1→ CGA(R) –1→ CGG(R) –1→ AGG(R)
12: AAA(K) –1→ AAT(N) –2→ GAC(D) –1→ GAG(E) –1→ AAG(K) –1→ AGG(R)
13: GCG(A) –1→ GTG(V) –1→ GTC(V) –1→ CTC(L) –1→ ATC(I) –2→ TAC(Y)
14: TTG(L) –1→ CTG(L) –1→ CTC(L)
15: GAC(D) –1→ GGC(G) –1→ GGG(G) –1→ GGT(G) –1→ GTT(V)
16: CAG(Q) –1→ CAT(H) –1→ CAA(Q)
17: CTG(L) –1→ TTG(L) –1→ TTA(L)
18: TCC(S) –1→ TCG(S) –1→ TCT(S) –1→ TCA(S) –1→ ACA(T) –2→ CGA(R)
–2→ AGT(S)
19: GCC(A) –1→ GCG(A) –1→ GCA(A) –1→ GCT(A)
20: GGG(G) –1→ AGG(R) –1→ AGA(R) –1→ CGA(R) –1→ CAA(Q)
21: AAG(K) –1→ AAA(K) –2→ CAT(H)
22: CAC(H) –1→ TAC(Y) –1→ TAT(Y) –2→ TTC(F)
23: CTT(L) –1→ CTG(L) –1→ CGG(R)
24: CAG(Q) –1→ CAC(H) –1→ CAT(H) –1→ CAA(Q)
25: TCC(S) –1→ TCG(S) –1→ TGG(W) –2→ TTT(F) –1→ TCT(S) –1→ ACT(T)
–1→ ACA(T)
26: GTC(V) –1→ CTC(L) –1→ CTG(L) –1→ CTT(L)
27: GTG(V) –1→ ATG(M)

**Supplementary Table 4/3. Position-specific mutational pathways in PRP sequences.** For each residue position, the codons observed across the analysed species are connected by the minimal mutational routes based exclusively on the detected codons. Numbers above the arrows indicate the minimal number of nucleotide substitutions required for each transition. Amino acids are colored according to residue class: acidic (red), basic (blue), polar (light green), and hydrophobic (gray).

1: CAT(H) –1→ CAC(H)
2: GCT(A) –1→ GCA(A) –1→ GCG(A) –1→ GCC(A)
3: GAT(D) –1→ GAC(D)
4: GGA(G) –1→ GGT(G) –1→ GGG(G) –1→ GCG(A)
5: GTT(V) –1→ ATT(I) –2→ CTC(L) –1→ CTG(L) –1→ TTG(L)
6: TTC(F)
7: ACA(T) –1→ ACT(T) –1→ ACG(T) –1→ ACC(T) –2→ CAC(H)
8: AGT(S) –1→ AGC(S) –1→ AAC(N)
9: GAT(D) –1→ GAC(D) –1→ AAC(N) –2→ GTC(V) –1→ GGC(G) –1→ GGA(G)
10: TTC(F) –1→ TAC(Y) –1→ CAC(H)
11: AGT(S) –1→ AGC(S) –2→ AAG(K)
12: AGA(R) –1→ AAA(K) –1→ AAG(K) –2→ CAT(H)
13: CTC(L) –1→ CTT(L) –1→ CTG(L)
14: TTG(L) –1→ CTG(L) –1→ CTA(L) –1→ CTC(L)
15: GCT(A) –1→ GGT(G) –1→ GGA(G) –1→ GGC(G) –1→ GGG(G)
16: CAG(Q) –1→ CAA(Q) –1→ AAA(K)
17: CTT(L) –1→ ATT(I) –1→ ATG(M) –1→ CTG(L) –2→ TTA(L)
18: TCT(S) –1→ TCA(S) –1→ TCC(S)
19: GCC(A) –1→ GCA(A) –1→ GCT(A) –1→ GCG(A) –1→ GTG(V)
20: AAG(K) –1→ AAA(K) –1→ AGA(R) –2→ CGG(R) –1→ CGC(R)
21: GAG(E) –1→ AAG(K) –1→ AAA(K) –1→ AGA(R) –1→ AGG(R) –1→ CGG(R) –1→ CGC(R)
22: TAC(Y) –1→ TAT(Y)
23: CTT(L) –1→ CTA(L) –1→ CTG(L) –1→ TTG(L) –1→ TTC(F)
24: GAG(E) –1→ GAA(E) –2→ CAT(H)
25: TCT(S) –1→ TCA(S) –1→ TCG(S) –1→ TCC(S)
26: CTT(L) –1→ CTC(L) –1→ CTG(L) –1→ TTG(L) –1→ TTA(L)
27: ATT(I) –1→ ATC(I) –1→ ATG(M) –1→ CTG(L)

**Supplementary Table 4/4. Position-specific mutational pathways in PH sequences.** For each residue position, the codons observed across the analysed species are connected by the minimal mutational routes based exclusively on the detected codons. Numbers above the arrows indicate the minimal number of nucleotide substitutions required for each transition. Amino acids are colored according to residue class: acidic (red), basic (blue), polar (light green), and hydrophobic (gray).

1: TAT(Y) –1→ CAT(H) –1→ CAC(H)
2: GCT(A) –1→ GCC(A) –1→ GCA(A) –1→ GTA(V) –1→ GTG(V)
3: GAT(D)
4: GCC(A)
5: ATC(I) –1→ ATT(I) –1→ ATA(I)
6: TTC(F) –1→ TTT(F)
7: ACC(T) –1→ ACT(T) –1→ ACA(T)
8: ACC(T) –1→ AAC(N) –1→ GAC(D)
9: AAC(N) –1→ AGC(S) –1→ AGT(S) –2→ ACA(T)
10: TAC(Y) –1→ TAT(Y)
11: CGT(R) –1→ CGG(R) –1→ AGG(R) –1→ AGA(R)
12: AAG(K) –1→ AAA(K)
13: GTG(V) –1→ GTT(V) –1→ GTC(V) –1→ CTC(L) –1→ TTC(F)
14: CGA(R) –2→ CTT(L) –1→ CTG(L) –1→ TTG(L)
15: AGC(S) –1→ GGC(G) –1→ GGG(G) –1→ GGT(G)
16: AAG(K) –1→ CAG(Q) –1→ CAA(Q)
17: CTG(L) –2→ ATT(I) –1→ ATA(I) –1→ ATC(I)
18: TCC(S) –1→ TCA(S) –1→ TCT(S) –1→ TAT(Y)
19: GCC(A)
20: CGC(R) –1→ CGG(R) –1→ CAG(Q) –2→ AGG(R) –1→ AGA(R)
21: AAA(K) –1→ AAG(K) –1→ AGG(R)
22: CTG(L) –1→ GTG(V) –2→ ATT(I) –1→ TTT(F) –1→ TTA(L) –1→ TTC(F)
–1→ TAC(Y)
23: ATC(I) –1→ CTC(L) –1→ CTG(L) –1→ TTG(L) –1→ TTA(L) –1→ CTA(L)
–1→ CTT(L)
24: CAG(Q) –1→ CAA(Q)
25: GAC(D) –1→ AAC(N) –1→ ACC(T) –1→ ACT(T) –2→ GGT(G)
26: ATC(I) –1→ ATG(M) –1→ ATT(I) –1→ GTT(V)
27: ATG(M) –1→ ATA(I) –1→ ATT(I) –1→ GTT(V)

**Supplementary Table 4/5. Position-specific mutational pathways in GHRH sequences.** For each residue position, the codons observed across the analysed species are connected by the minimal mutational routes based exclusively on the detected codons. Numbers above the arrows indicate the minimal number of nucleotide substitutions required for each transition. Amino acids are colored according to residue class: acidic (red), basic (blue), polar (light green), and hydrophobic (gray).

1: CAC(H) –1→ CAT(H)
2: TCA(S) –1→ TCG(S) –2→ GCT(A) –1→ GTT(V)
3: GAC(D) –1→ GAT(D)
4: GGG(G) –1→ GGA(G)
5: ACG(T) –1→ AGG(R) –1→ ATG(M) –1→ CTG(L) –1→ CTT(L)
6: TTC(F) –1→ TTT(F)
7: ACC(T) –2→ CAC(H)
8: AGC(S) –1→ AGT(S)
9: GAG(E) –1→ GAA(E)
10: CTC(L) –1→ TTC(F) –1→ TAC(Y)
11: AGC(S)
12: CGC(R) –1→ CGA(R) –3→ AAG(K)
13: CTG(L) –1→ TTG(L) –1→ ATG(M) –3→ GCC(A)
14: CAG(Q) –1→ CGG(R) –1→ AGG(R) –1→ AGA(R) –2→ AAT(N)
15: GAG(E) –1→ GAC(D) –2→ GGA(G)
16: GGC(G) –2→ AGT(S) –1→ AAT(N) –1→ AAC(N) –3→ TCA(S)
17: GCG(A) –1→ GCC(A) –1→ GCT(A)
18: AGG(R) –1→ CGG(R) –1→ CAG(Q) –2→ TAC(Y) –3→ GCT(A)
19: CTC(L) –1→ CTG(L) –1→ GTG(V) –2→ ATA(I)
20: CAG(Q) –2→ CGC(R)
21: CGC(R) –1→ CGG(R) –1→ CAG(Q) –1→ AAG(K)
22: CTG(L) –2→ TTT(F) –2→ ATC(I)
23: CTA(L) –1→ CTG(L) –1→ GTG(V) –2→ ATC(I)
24: CAG(Q) –1→ CAA(Q) –1→ AAA(K) –1→ AAC(N)
25: GGC(G) –1→ GGT(G) –2→ TCT(S) –2→ AAT(N) –2→ CAC(H)
26: CTG(L) –1→ CTC(L) –3→ GCT(A)
27: GTG(V) –1→ ATG(M) –2→ CTT(L)

**Supplementary Table 4/6. Position-specific mutational pathways in SCT sequences.** For each residue position, the codons observed across the analysed species are connected by the minimal mutational routes based exclusively on the detected codons. Numbers above the arrows indicate the minimal number of nucleotide substitutions required for each transition. Amino acids are colored according to residue class: acidic (red), basic (blue), polar (light green), and hydrophobic (gray).

1: CAT(H) –1→ CAC(H)
2: TCA(S) –1→ TCG(S) –1→ TCC(S)
3: CAG(Q) –1→ CAA(Q) –1→ GAA(E) –1→ GAG(E)
4: GGC(G) –1→ GGT(G) –1→ GGA(G) –1→ GGG(G)
5: ACA(T) –1→ ACT(T) –1→ ACG(T) –1→ ACC(T) –1→ TCC(S)
6: TTC(F) –1→ TTT(F)
7: ACC(T) –1→ ACT(T) –1→ ACG(T) –1→ ACA(T) –1→ TCA(S) –1→ TCC(S)
8: AGT(S) –1→ AGC(S) –1→ AAC(N) –1→ AAT(N)
9: GAC(D) –1→ GAT(D)
10: TAC(Y)
11: AGC(S) –1→ AGT(S)
12: AAG(K) –1→ AAA(K)
13: TAT(Y) –1→ TAC(Y) –1→ TTC(F) –2→ CAC(H)
14: TTG(L) –1→ CTG(L) –1→ CTA(L) –1→ CTC(L) –1→ CTT(L)
15: GAC(D) –1→ GAG(E) –1→ GAT(D)
16: TCC(S) –2→ ACG(T) –1→ ACC(T) –1→ AGC(S) –1→ AAC(N) –1→ GAC(D)
–1→ GAA(E) –2→ GTG(V)
17: CGC(R) –2→ AGA(R) –1→ AGG(R) –1→ AAG(K)
18: CGT(R) –1→ CGA(R) –1→ CGG(R) –1→ CAG(Q) –1→ AAG(K) –1→ AGG(R)
–1→ AGA(R)
19: GCC(A) –1→ GCT(A) –1→ GCA(A) –1→ GCG(A)
20: CAA(Q) –1→ CAG(Q) –1→ AAG(K)
21: GAT(D) –1→ GAC(D)
22: TTT(F) –1→ TTC(F)
23: GTG(V) –1→ GTT(V) –1→ GTA(V) –1→ GTC(V) –1→ ATC(I)
24: CAG(Q) –1→ CAA(Q) –1→ CGA(R) –1→ CGC(R) –2→ ACC(T)
25: TGG(W)
26: TTG(L) –1→ TTA(L) –1→ CTA(L) –1→ CTC(L) –1→ CTG(L)
27: AAA(K) –1→ AAG(K) –1→ ATG(M) –1→ TTG(L)

**Supplementary Table 4/7. Position-specific mutational pathways in GCG sequences.** For each residue position, the codons observed across the analysed species are connected by the minimal mutational routes based exclusively on the detected codons. Numbers above the arrows indicate the minimal number of nucleotide substitutions required for each transition. Amino acids are colored according to residue class: acidic (red), basic (blue), polar (light green), and hydrophobic (gray).

1: CAT(H) –1→ CAC(H)
2: GCA(A) –1→ GCC(A) –1→ GCT(A) –1→ CCT(A)
3: GAA(E) –1→ GAT(D) –1→ GAC(D) –1→ GAG(E)
4: GGG(G) –1→ GGC(G) –1→ GGT(G) –1→ GGA(G)
5: ACC(T) –1→ ACA(T)
6: TTT(F) –1→ TAT(Y) –1→ TAC(Y) –1→ TTC(F)
7: ACC(T) –1→ ACT(T) –1→ ACG(T)
8: AGT(S) –1→ AAT(N) –1→ AAC(N) –1→ AGC(S)
9: GAT(D) –1→ GAC(D)
10: GTA(V) –1→ GTG(V) –1→ ATG(M) –1→ ATC(I) –1→ GTC(V)
11: AGT(S) –1→ AGC(S) –1→ ACC(T) –1→ ACT(T)
12: TCT(S) –1→ TCG(S) –1→ TCA(S) –1→ TCC(S) –1→ GCC(A) –1→ ACC(T)
–1→ AAC(N) –2→ CAG(Q) –2→ GAA(E)
13: TAT(Y) –1→ TAC(Y) –1→ CAC(H)
14: TTG(L) –1→ CTG(L) –1→ CTC(L) –1→ CTA(L)
15: GAA(E) –1→ GAT(D) –1→ GAC(D) –1→ GAG(E) –1→ AAG(K) –1→ CAG(Q)
–1→ CAA(Q)
16: GGT(G) –1→ GGC(G) –1→ GAC(D) –1→ GAA(E) –1→ GAG(E) –1→ GCG(A)
17: CAA(Q) –1→ CAG(Q) –1→ AAG(K) –1→ AAA(K)
18: GCT(A) –1→ GCA(A) –1→ GCG(A) –1→ GCC(A)
19: GCA(A) –1→ GCC(A) –1→ GTC(V) –1→ ATC(I)
20: AAA(K) –1→ AAG(K) –1→ CAG(Q) –2→ CGC(R)
21: GAA(E) –1→ GAG(E) –1→ GAC(D) –2→ AGC(S)
22: TTC(F) –1→ TTT(F)
23: ATT(I) –1→ GTT(V) –1→ GTG(V) –2→ CTC(V)
24: GCT(A) –1→ TCT(S) –1→ TCC(S) –1→ GCC(A) –1→ GAC(D) –1→ GAG(E)
–2→ GGC(G) –1→ AGC(S)
25: TGG(W) –1→ AGG(R) –1→ AAG(K)
26: CTG(L) –1→ CTT(L) –1→ CTC(L) –1→ CTA(L) –1→ TTA(L) –1→ TTG(L)
27: ATA(I) –1→ AAA(K) –1→ AAG(K) –2→ GTG(V) –2→ GCC(A)

**Supplementary Table 4/8. Position-specific mutational pathways in GLP-1 sequences.** For each residue position, the codons observed across the analysed species are connected by the minimal mutational routes based exclusively on the detected codons. Numbers above the arrows indicate the minimal number of nucleotide substitutions required for each transition. Amino acids are colored according to residue class: acidic (red), basic (blue), polar (light green), and hydrophobic (gray).

1: CAT(H) –1→ CAC(H)
2: ATA(I) –1→ GTA(V) –2→ GCT(A) –1→ GCC(A) –1→ GCG(A) –1→ TCG(S)
3: GAT(D) –1→ GAC(D)
4: GGT(G) –1→ GGC(G) –1→ GGA(G)
5: TCT(S) –1→ TCC(S) –1→ ACC(T) –1→ AGC(S) –1→ AGT(S)
6: TTC(F) –1→ TTT(F)
7: TCT(S) –2→ ACA(T) –1→ ACC(T)
8: GAC(D) –1→ GAT(D) –1→ AAT(N) –1→ AGT(S) –1→ AGC(S)
9: GAG(E) –1→ GAT(D) –1→ GAC(D)
10: ATG(M) –1→ ATC(I) –1→ GTC(V) –1→ GTG(V)
11: AAC(N) –1→ AGC(S)
12: ACC(T) –2→ AAA(K) –1→ AAG(K) –2→ CTG(V)
13: ATT(I) –1→ ATC(I) –1→ GTC(V) –1→ GTG(V) –1→ ATG(M)
14: CTT(L) –1→ CTA(L) –1→ CTC(L) –1→ CTG(L) –1→ TTG(L)
15: GAT(D) –1→ GAC(D)
16: GAT(D) –1→ AAT(N) –1→ ATT(I) –2→ AGA(R) –1→ AGC(S) –2→ TCC(S)
17: CTT(L) –1→ ATT(I) –1→ ATC(I) –1→ ATG(M) –1→ CTG(L)
18: GCC(A) –1→ GCT(A) –1→ GCG(A) –1→ GCA(A) –1→ TCA(S)
19: ACC(T) –1→ GCC(A) –1→ GCA(A) –1→ GCT(A)
20: AGG(R) –2→ AAA(K) –1→ AAG(K) –1→ CAG(Q)
21: GAG(E) –1→ GAA(E) –1→ GAC(D) –1→ AAC(N)
22: TTC(F) –1→ TTT(F) –1→ TAT(Y)
23: ATC(I) –1→ ATA(I) –1→ TTA(L) –1→ CTA(L) –1→ CTG(L)
24: AAC(N) –1→ AAA(K) –1→ CAA(Q) –2→ GAT(D) –1→ GAG(E) –2→ CTG(L) –1→ CTC(L)
25: TGG(W)
26: TTG(L) –1→ CTG(L) –1→ GTG(V) –1→ GTA(V) –1→ GTC(V) –1→ CTC(L)
27: ATT(I) –1→ ATG(M) –2→ AAA(K)

**Supplementary Table 4/9. Position-specific mutational pathways in GLP-2 sequences.** For each residue position, the codons observed across the analysed species are connected by the minimal mutational routes based exclusively on the detected codons. Numbers above the arrows indicate the minimal number of nucleotide substitutions required for each transition. Amino acids are colored according to residue class: acidic (red), basic (blue), polar (light green), and hydrophobic (gray).

1: TAC(Y) –1→ TAT(Y)
2: GCG(A) –1→ TCG(S) –1→ TCA(S) –1→ GCA(A) –1→ GCT(A)
3: GAA(E) –1→ GAG(E)
4: GGG(G) –1→ GGA(G) –2→ GCC(A) –2→ TCA(S)
5: ACG(T) –1→ ACC(T) –1→ ACT(T) –1→ ATT(I)
6: TTC(F) –1→ TTA(L) –1→ TTG(L) –1→ CTG(L) –2→ ATT(I)
7: ATC(I) –2→ GCC(A) –1→ GCG(A)
8: AGT(S) –1→ AGC(S)
9: GAC(D) –1→ GAT(D)
10: TAC(Y) –3→ ATT(I)
11: AGT(S) –1→ AGC(S)
12: ATT(I) –1→ ATC(I) –2→ CGC(R) –1→ CGA(R) –1→ AGA(R) –1→ AAA(K)
13: GCC(A) –1→ ACC(T) –1→ ATC(I) –2→ ACG(T) –2→ TCA(S)
14: ATG(M) –1→ TTG(L) –1→ GTG(V)
15: GAC(D) –1→ GAT(D)
16: AAC(N) –1→ AAG(K) –2→ TCG(S)
17: ATT(I) –1→ ATC(I) –1→ ATG(M)
18: CAC(H) –2→ CGA(R) –2→ CTG(L) –2→ GTT(V)
19: CAA(Q) –1→ AAA(K) –1→ AAG(K) –1→ CAG(Q)
20: CAA(Q) –1→ AAA(K) –1→ AAG(K)
21: GAC(D) –1→ AAC(N)
22: TTT(F) –1→ TTC(F)
23: GTG(V)
24: AAC(N) –1→ GAC(D) –1→ GAG(E) –1→ GAA(E)
25: TGG(W) –2→ TTT(F)
26: CTG(L) –1→ CTC(L) –2→ TTA(L)
27: TTG(L) –1→ CTG(L) –1→ CTT(L)

**Supplementary Table 4/10. Position-specific mutational pathways in GIP sequences.** For each residue position, the codons observed across the analysed species are connected by the minimal mutational routes based exclusively on the detected codons. Numbers above the arrows indicate the minimal number of nucleotide substitutions required for each transition. Amino acids are colored according to residue class: acidic (red), basic (blue), polar (light green), and hydrophobic (gray).

| | H-YLAAVL-OH (polymorph A) | H-YLAAVL-OH (polymorph B) | Ac-YLAAVL-OH |
|---|---|---|---|
| Data collection | | | |
| Unit cell: a,b,c (Å) | 9.527, 11.616, 16.76 | 20.035, 18.703, 20.616 | 42.039, 8.614, 11.654 |
| α,β,γ (°) | 83.93, 85.63, 81,98 | 90, 92.69, 90 | 90, 96.76, 90 |
| Space group | P1 | $P2_1$ | C2 |
| Resolution range (Å) | 16.63 – 1.50 (1.55 – 1.50) | 14.70 – 1.15 (1.19 – 1.15) | 11.57 – 1.25 (1.29 – 1.25) |
| No. of unique refl. / observed refl. | 1139 (125) / 4489 (326) | 5520 (547) / 21813 (1694) | 1148 (54) / 4821 (91) |
| <I / σI> | 5.3 (1.3) | 5.3 (1.8) | 17.2 (4.2) |
| $R_{meas}$ | 0.186 (0.948) | 0.167 (0.591) | 0.056 (0.033) |
| Completeness (%) | 99.56 (98.43) | 99.19 (98.92) | 91.62 (41.54) |
| CC(1/2) | 0.989 (0.766) | 0.998 (0.882) | 0.999 (0.986) |
| Refinement | | | |
| Resolution range (Å) | 16.63 – 1.50 | 14.70 – 1.15 | 11.57 – 1.25 |
| R / $R_{free}$ (No. of obs.) | 0.1517 (1018) / 0.2038 (113) | 0.2130 (4883) / 0.2417 (552) | 0.0615 (1028) / 0.0846 (115) |
| B factor of peptide / solvent (Å$^2$) | | | |
| RMS dev. bond length (Å) | 0.014 | 0.008 | 0.017 |
| RMS dev. bond angles (°) | 1.774 | 0.953 | 1.633 |
| Ramachandran fav/all/disall | | | |
| No. of non hydrogen atoms: peptide / solvent | | | |
| PDB code | 29VP | 29VQ | 29VO |

| | H-YLNSIL-OH | H-YLESLM-OH | Ac-YLESLM-OH |
|---|---|---|---|
| Data collection | | | |
| Unit cell: a,b,c (Å) | 4.919, 17.005, 45.529 | 4.833, 18.483, 43.363 | 18.957, 9.492, 24.654 |
| α,β,γ (°) | 90, 90, 90 | 90, 90, 90 | 90, 103.94, 90 |
| Space group | $P2_12_12_1$ | $P2_12_12_1$ | I2 |
| Resolution range (Å) | 22.764 – 1.070 (1.088 – 1.070) | 21.69 – 1.00 (1.04 – 1.00) | 16.65 – 1.24 (1.28 – 1.24) |
| No. of unique refl. / observed refl. | 1565 (19) / 16218 (42) | 2436 (221) / 10681 (774) | 1206 (35) / 4035 (43) |
| <I / σI> | 7.5 (1.0) | 4.9 (1.6) | 10.0 (1.8) |
| $R_{meas}$ | 0.174 (0.741) | 0.249 (1.820) | 0.086 (1.284) |
| Completeness (%) | 78.5 (20.9) | 99.96 (100.00) | 92.13 (27.13) |
| CC(1/2) | 0.997 (0.939) | 0.992 (0160) | 0.996 (0.148) |
| Refinement | | | |
| Resolution range (Å) | 22.76 – 1.20 | 21.67 – 1.00 | 16.65 – 1.24 |
| R / $R_{free}$ (No. of obs.) | 0.0822 (1293) / 0.0709 (55) | 0.2002 (2143) / 0.2236 (234) | 0.0938 (1083) / 0.1020 (121) |
| B factor of peptide / solvent (Å$^2$) | | | |
| RMS dev. bond length (Å) | 0.009 | 0.023 | 0.012 |
| RMS dev. bond angles (°) | 1.476 | 1.764 | 1.312 |
| Ramachandran fav/all/disall | | | |
| No. of non hydrogen atoms: peptide / solvent | | | |
| PDB code | 29VR | 29VS | 29VT |

| | H-LLQGLV-OH | Ac-LLQDIM-$NH_2$ |
|---|---|---|
| Data collection | | |
| Unit cell: a,b,c (Å) | 19.156, 19.739, 22.459 | 38.76, 4.83, 24.40 |
| α,β,γ (°) | 90, 99.43, 90 | 90, 112.008, 90 |
| Space group | $P2_1$ | C2 |
| Resolution range (Å) | 22.16 – 0.86 (0.89 – 0.86) | 17.968 – 1.379 (1.518 – 1.379) |
| No. of unique refl. / observed refl. | 13773 (1052) / 49910 (1823) | 945 (164) / 3285 (296) |
| <I / σI> | 7.2 (1.0) | 6.5 (0.9) |
| $R_{meas}$ | 0.094 (0.822) | 0.119 (0.825) |
| Completeness (%) | 97.34 (75.30) | 92.5 (68.6) |
| CC(1/2) | 0.999 (0.693) | 0.996 (0.398) |
| Refinement | | |
| Resolution range (Å) | 18.90 – 0.86 | 17.97 – 1.41 |
| R / $R_{free}$ (No. of obs.) | 0.1693 (12336) / 0.1811 (1374) | 0.1378 (862) / 0.1732 (56) |
| B factor of peptide / solvent (Å$^2$) | | |
| RMS dev. bond length (Å) | 0.015 | 0.014 |
| RMS dev. bond angles (°) | 1.910 | 1.492 |
| Ramachandran fav/all/disall | | |
| No. of non hydrogen atoms: peptide / solvent | | |
| PDB code | 29WR | 29VU |

**Supplementary Table 5. X-ray data collection and refinement statistics.** Data for the highest resolution shell are given in parentheses.

1. Micsonai, A. *et al.* Accurate secondary structure prediction and fold recognition for circular dichroism spectroscopy. *Proceedings of the National Academy of Sciences* **112**, E3095–E3103 (2015).
2. Eisenberg, D., Weiss, R. M. & Terwilliger, T. C. The helical hydrophobic moment: a measure of the amphiphilicity of a helix. *Nature* **299**, 371–374 (1982).
3. Cong, Z. *et al.* Structural perspective of class B1 GPCR signaling. *Trends in Pharmacological Sciences* **43**, 321–334 (2022).
4. Chang, X., Keller, D., O'Donoghue, S. I. & Led, J. J. NMR studies of the aggregation of glucagon-like peptide-1: formation of a symmetric helical dimer. *FEBS Letters* **515**, 165–170 (2002).
5. Cardoso, J. C., Vieira, F. A., Gomes, A. S. & Power, D. M. The serendipitous origin of chordate secretin peptide family members. *BMC Evolutionary Biology* **10**, 135 (2010).
6. Handbook of Hormones - 2nd Edition. https://www.elsevier.com/books/handbook-of-hormones/ando/978-0-12-820649-2.
7. Sherwood, N. M., Krueckl, S. L. & McRory, J. E. The Origin and Function of the Pituitary Adenylate Cyclase-Activating Polypeptide (PACAP)/Glucagon Superfamily*. *Endocr Rev* **21**, 619–670 (2000).
8. Tam, J. K. V., Lee, L. T. O. & Chow, B. K. C. PACAP-related peptide (PRP)—Molecular evolution and potential functions. *Peptides* **28**, 1920–1929 (2007).
